# Transcriptional networks underlying tumour plasticity in small-cell lung cancer

**DOI:** 10.64898/2026.09.26.754526

**Authors:** Debadrita Bhattacharya, Sarah M. Groves, Cameron Walker, Gina N. Duronio, Marcus Hsieh, Yamina Chtourou, Griffin G. Hartmann, Wen-Hao Hsu, Candace C. Liu, Michael Angelo, Vito Quaranta, Julien Sage

## Abstract

Across human malignancies, the ability of cancer cells to switch states (“plasticity”) drives progression, metastasis and therapy resistance ^1–3^. Here we investigated the transcriptional and epigenetic basis of plasticity in small-cell lung cancer (SCLC), an aggressive, treatment-refractory cancer ^4, 5^. SCLC tumours display plasticity along a neuroendocrine (NE) to nonNE axis, however the trajectories of lineage transition and the gene regulatory network underlying it, remain poorly defined ^6^. Using single-cell multi-omics, we resolved distinct transcriptional SCLC cell states, including a previously unrecognised Intermediate state comprised of low-identity immunogenic SCLC cells that serve as an obligatory route in NE to nonNE lineage transition. Using single-cell gene expression and chromatin accessibility data, we next developed a <u>Bo</u>olean <u>Ba</u>yes model for transcription factor networks (BoBa-T) to predict regulators of SCLC cell states. We validated RORB as a novel gatekeeper of NE state that represses the nonNE identity in SCLC. RORB inactivation drives NE cells into the Intermediate state, upregulates immunogenic programs and inhibits tumour growth in immunocompetent hosts. Our work identifies a plastic, immunogenic waypoint for SCLC lineage switching and reveals strategies to target it therapeutically.

## INTRODUCTION

Intratumoural heterogeneity is a hallmark of human tumours ^7^. A central question in cancer biology is how cancer cells switch between cell states without new genetic alterations, allowing them to adapt rapidly to their microenvironment during tumour progression and in response to treatment. In SCLC, tumours are commonly classified by transcriptional subtypes: tumours driven by the transcription factors (TFs) ASCL1 (Achaete-scute homolog 1) and NEUROD1 (Neurogenic differentiation 1) are typically considered NE tumours, while POU2F3-driven (POU domain, class 2, transcription factor 3) and immune-inflamed tumours (which can express YAP1, Yes-associated protein 1) are considered nonNE tumours ^8–10^. Intratumoural heterogeneity mirrors this intertumoural classification, with individual SCLC tumours often comprising cancer cells displaying mixed features of NE and nonNE cell states ^11–19^. Activation of the Notch signalling pathway, the REST (Repressor Element 1 Silencing Transcription factor) transcriptional regulator, and the YAP1 TF can promote this transition from NE to nonNE ^20–25^. Epigenetic regulators such as EZH2 and LSD1 (a Notch signalling repressor) can also control SCLC cell states ^26–28^. However, the regulatory networks governing these transitions and the cellular intermediates that enable them remain poorly characterised, with most studies focusing on terminally differentiated rather than transitional states.

Here, we employed single-cell multi-omics to systematically dissect SCLC heterogeneity. We performed our initial analysis of SCLC cell states in a well-validated mouse model (*RPR2*, *Rb/p53/Rbl2* mutant mice) in which transcriptional and epigenetic heterogeneity is prominent, and where cancer cells occupy both the NE and nonNE lineages ^22, 29–31^, before validation in human tumours. Our studies describe SCLC cell states and their regulators, revealing mechanisms underlying plasticity and therapeutic opportunities in this fatal cancer.

## RESULTS

### Archetype analysis defines distinct gene programs in SCLC cell populations

We isolated pure populations of live cancer cells by flow cytometry from end-stage primary lung tumours (24 weeks post cancer initiation) in *RPR2* mutant mice ^23^ (**Supplementary Fig. 1a**) and performed combined single-cell RNA and ATAC sequencing (scRNA-seq and scATAC-seq) using the 10x Genomics platform (**Fig. 1a**). In Uniform Manifold Approximation and Projection (UMAP)-based dimensionality reduction, the ∼9,000 high-quality cancer cells segregated primarily by neuroendocrine (NE) or non-neuroendocrine (nonNE) identity (**Supplementary Fig. 1b,c**), consistent with previous work ^22, 32^. To map the continuum of cancer cell states, we used Archetype Analysis (AA) to identify the most divergent gene-expression programs in the scRNA-seq data ^33, 34^. Unlike graph-based clustering, archetype analysis identifies the most extreme and mutually irreducible transcriptional states and assigns each cell a probability of belonging to each. Cells with high probability for a single state are termed “Specialists” (∼3,500 cells), while “Generalists” co-opt programs from multiple archetypes within the NE state or the nonNE state (**Fig. 1b** and **Supplementary Fig. 2a**). This analysis revealed six archetypal cell states, defined by their marker genes and associated pathways (**Fig. 1b-g** and **Supplementary Fig. 1d**): two NE states (NE1 and NE2), both marked by high levels of *Ascl1* and an NE gene signature (akin to human ASCL1^high^ tumours, SCLC-A), but differing in expression of the pro-metastatic transcription factor *Nfib*^30^; two nonNE states (nonNE1 and nonNE2) with elevated Notch and Hippo signalling and no expression of *Ascl1* ^22, 32^; and two previously uncharacterised low-NE states (Secretory and Intermediate) with moderate to low expression of *Ascl1* and its target genes. The Secretory archetype upregulated gene programs for MAPK signalling, cell-cell communication, and protein secretion, suggesting a paracrine role within the tumour microenvironment (**Supplementary Fig. 2a-e**). These features resemble a specialised NE archetype (SCLC-A2) in human SCLC previously associated with chemotherapy resistance ^35^.

**Figure 1:**
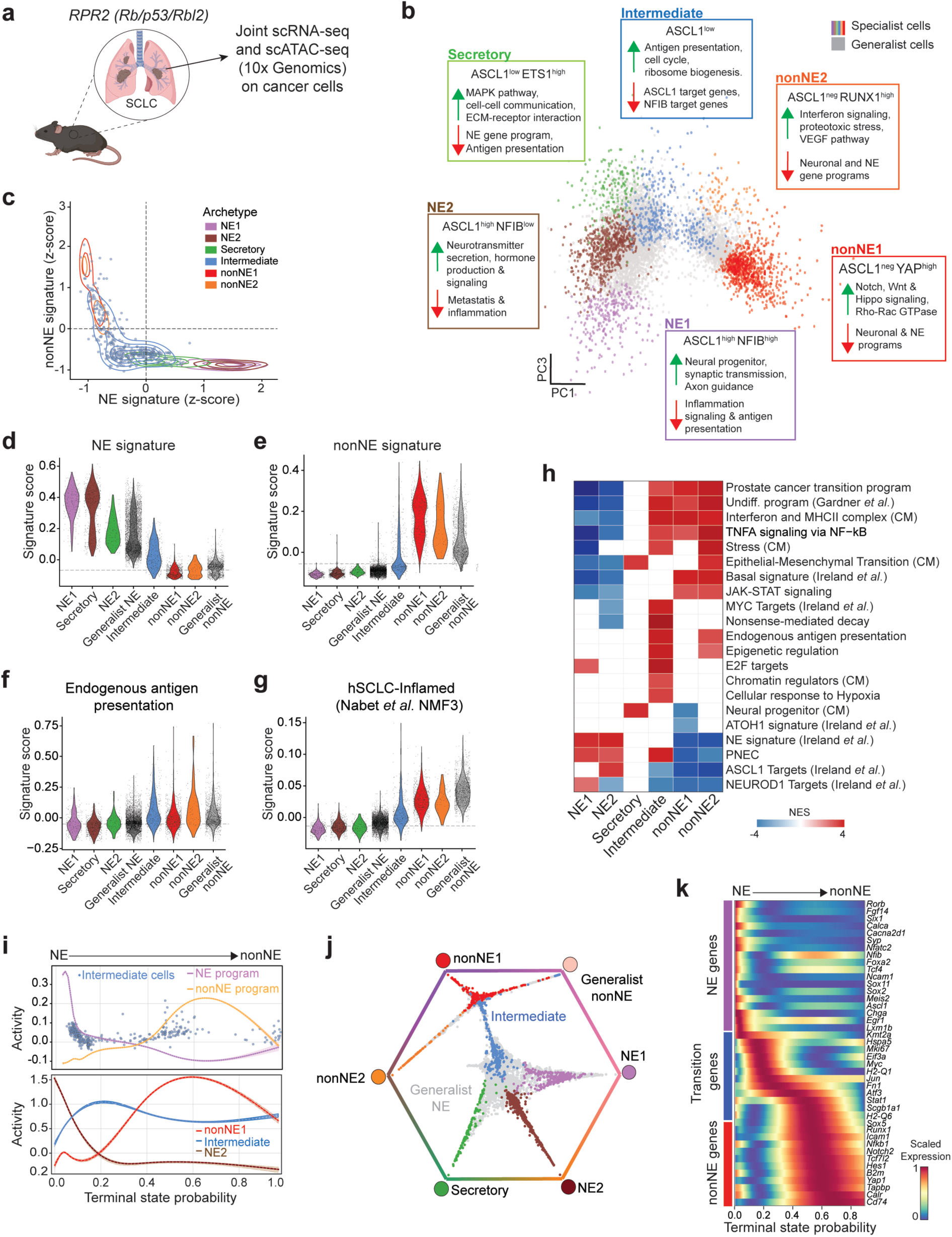
Single-cell multi-omics reveals SCLC cell state diversity. **a.** Experimental workflow for paired single-cell RNA sequencing (scRNA-seq) and single-cell assay for transposase-accessible chromatin sequencing (scATAC-seq) profiling of *RPR2* mutant (*Rb/p53/Rbl2*) mouse SCLC cancer cells isolated from end-stage primary lung tumours by flow cytometry (n = 6 mice, 3 males and 3 females). **b.** Principal component analysis (PCA) rendering of the scRNA-seq modality showing the six identified archetypes and associated gene programs. “Specialist” cancer cells are coloured according to maximum lineage probability, and archetypes are overlaid as coloured nodes. “Generalist” cells are shown in grey. **c.** Contour plots showing the spread of cancer cells in each archetype based on the activity of NE versus nonNE signatures in individual cells. Kernel density contours depict the local density of cells in this space, with inner contours indicating regions of highest cell density and outer contours enclosing progressively lower-density regions. Colours indicate archetype assignment as in (b). Individual Intermediate cells are shown in blue. **d-g.** Violin plots comparing the activity of the NE gene signature (d), nonNE gene signature (e), endogenous antigen presentation pathway (f), and the human inflamed NE SCLC signature (g) across *RPR2* archetypes. **h.** Heatmap showing normalised enrichment score (NES) of select cancer-relevant gene signatures across *RPR2* archetypes. Only significant enrichments (FDR < 0.05) are shown. CM, Cancer Metaprogram. **i.** Spline fits across pseudotime for the NE and nonNE gene programs (top) and for the NE2, Intermediate and nonNE1 archetype signatures (bottom). Intermediate cells, shown in blue (top), predominantly occupy the minimum between the NE and nonNE identities. **j.** Circular projection of single cells according to their fate probabilities. Fate-biased cells are placed next to their corresponding colours, while lineage-naïve cells are placed in the middle. Cells are coloured based on their respective archetypes. **k.** Heatmap of gene trends of select putative driver genes ordered by NE to nonNE transition pseudotime (scale gene trends of imputed expression).

The Intermediate cells mapped between the NE and nonNE arms in principal component analysis (PCA) space, representing a low-identity, cycling population (**Fig. 1b,c** and **Supplementary Fig. 2a**). Despite low expression of lineage genes, Intermediate cells were highly enriched for translational control, ribosome biogenesis, nonsense-mediated decay (NMD), antigen presentation, MHC expression, hypoxia and TNFα-NFκB signalling (**Fig. 1d-g** and **Supplementary Fig. 2b-d,f,g**). These features distinguished Intermediate from canonical NE cells, which were dominated by neuronal and synapse formation pathways, strongly downregulated translational control and NMD, and lacked antigen presentation genes (**Fig. 1h**, **Supplementary Fig. 2h,i**). NonNE cells upregulated Notch signalling, cytokine-associated immune and JAK-STAT pathways but were not enriched for antigen presentation or translation/NMD pathways (**Fig. 1h**, **Supplementary Fig. 2j**).

We addressed three potential confounders in our characterisation of the Intermediate state. First, to exclude this state as a heterotypic doublet of NE and nonNE cells, we examined DoubletFinder pANN scores, which are high for technical doublets; Intermediate had the lowest median of all archetypes (**Supplementary Fig. 3a)**. Second, to exclude it as an artificial split of a single archetype, we varied the number of archetypes (k) fitted (k = 3-8); the Intermediate was consistently recovered as a distinct state across this range (**Supplementary Fig. 3b**). Third, since roughly 25% of Intermediate cells are mitotic, we tested whether its enriched pathways simply reflect its proliferative state. Only chromatin organisation and ribosome biogenesis varied with cell-cycle phase, rising modestly in G2/M cells but far less than E2F targets, while antigen presentation, NMD and inflammatory signalling were cell-cycle phase-independent (**Supplementary Fig. 3c**); irrespective of archetype, a cell-cycle phase-specific gene set enrichment analysis (GSEA) confirmed that Intermediate and G2/M cells shared only cell-cycle pathways (**Supplementary Fig. 3d**).

We next performed single-cell spatial proteomics of ∼10^6^ cells from *RPR2* primary tumour sections using MIBI (Multiplexed Ion Beam Imaging). Cancer cells co-expressing NE and nonNE protein markers (ASCL1, Synaptophysin, and ICAM1), albeit at levels lower than their differentiated counterparts, were identified as candidate Intermediate cells; these cells also displayed high levels of c-JUN, EpCAM, and Vimentin, consistent with the scRNA-seq data (**Supplementary Fig. 4a-d**). Notably, Intermediate cells were found throughout the tumours but were enriched at the tumour border (**Supplementary Fig. 4e,f**) (see below).

Together, these results establish the Intermediate state in advanced *RPR2* tumours as a stable cycling state with distinct transcriptional identity.

### The Intermediate state represents a transition state between NE and nonNE SCLC cells

We next asked how these archetypes relate to low-identity cancer cell states described in other systems, including “inflamed basal” SCLC cells, and to human SCLC subtypes. The SCLC Basal (Ireland *et al.* ^12^) and prostate adenocarcinoma-to-neuroendocrine transition (Chan *et al.* ^35^) signatures peaked in nonNE2 cells, whereas the lung adenocarcinoma (LUAD) to SCLC transformation program (Gardner *et al.* ^36^) was enriched in Intermediate as well as nonNE1/2 cells (**Fig. 1h and Supplementary Fig. 5a,b**). A human ASCL1^high^ signature (Nabet *et al.*, ^37^) was highly active in NE1/2 and Secretory states, low in Intermediate, and absent from nonNE1/2 (**Supplementary Fig. 5c**). NEUROD1^high^ (SCLC-N) ^37^ and POU2F3^high^ (SCLC-P) (Gay *et al.*, ^8^) signatures scored low across all *RPR2* archetypes, as expected in this SCLC-A mouse model (**Supplementary Fig. 5d,e**); the slight SCLC-N elevation in NE1 may reflect NFIB-driven neuronal programs ^38^. Inflamed subtype (SCLC-I) signatures ^8, 37^ were strongly active in Intermediate and nonNE1/2 states (**Fig. 1h** and **Supplementary Fig. 5f**). Finally, cross-signature correlation placed Intermediate apart from NE and nonNE, closest to the human SCLC-I subtype and the undifferentiated LUAD-SCLC program (**Supplementary Fig. 5g**). The inflamed program itself, however, correlated more strongly with nonNE archetypes, indicating that the Intermediate state partly recapitulates the human SCLC-I state and is not uniquely comparable to other plastic cancer cell states.

We used CellRank ^39^ to model archetype dynamics in the *RPR2* dataset. The predominant trajectory followed the NE (“early”) to nonNE (“late”) lineage switch, with early downregulation of NE genes and eventual upregulation of the nonNE program (**Fig. 1i**). Of the seven terminal states recovered along this trajectory, six corresponded to the archetypes and one to Generalist nonNE. The NE1, NE2 and Secretory states diverged early in this trajectory while the nonNE1, nonNE2 and Generalist nonNE states appeared later. This trajectory passed through a bottleneck composed entirely of Intermediate cells and marked by Intermediate-associated genes (**Fig. 1j,k**), positioning this low-identity population as a transition state between the NE and nonNE lineages.

We next asked how this trajectory evolves during tumorigenesis. We performed scRNA-seq on sorted cancer cells at 4 months post cancer initiation, when microscopic tumours first appear, and at 5 months, when tumours are macroscopically distinct but well below end-stage size (6 months). Clustering showed greater transcriptional heterogeneity in cancer cells from 5-month compared to 4-month tumours (**Supplementary Fig. 6a**), and archetype labels transferred from the end-stage reference (at 6 months) recovered all six archetypes at these two early timepoints with high confidence (**Supplementary Fig. 6b-d**). 4-month tumours held a balanced NE/nonNE mix, whereas by 5 months they had shifted to an NE-enriched composition resembling end-stage tumours (**Supplementary Fig. 6b**).

We confirmed that label transfer mapped equivalent states across timepoints. The NE and nonNE archetype-labelled cells scored highest for their respective lineage programs, while Intermediate-labelled cells were low for both and, like nonNE cells, elevated for antigen presentation and inflammation (**Supplementary Fig. 6e-h**). Pathway enrichment showed high concordance between early (query) and late (reference) timepoint cells for each archetype (**Supplementary Fig. 6i**). The Intermediate state was the sole exception: its G2/M and cell-cycle programs were under-enriched early while its other programs remained similar, indicating that its proliferative feature emerges only in late-stage tumours and further supporting the notion that the definition of this state does not depend on its cell-cycle status.

We next integrated the Specialist archetype cells from the 6-month tumours with the 4-month and 5-month datasets and used moscot (multi-omics single-cell optimal transport, ^40^) to trace cell movement between states across the three timepoints (**Fig. 2a**). To determine the transcriptional basis of these transitions, we compared pathways upregulated in cells that remained in their state versus those that left it at each timepoint. Across all three timepoints, the terminal archetypes were highly self-renewing, most cells giving rise to cells in the same state at the next timepoint (**Fig. 2b-e** and **Supplementary Fig. 7a-d**).

**Figure 2:**
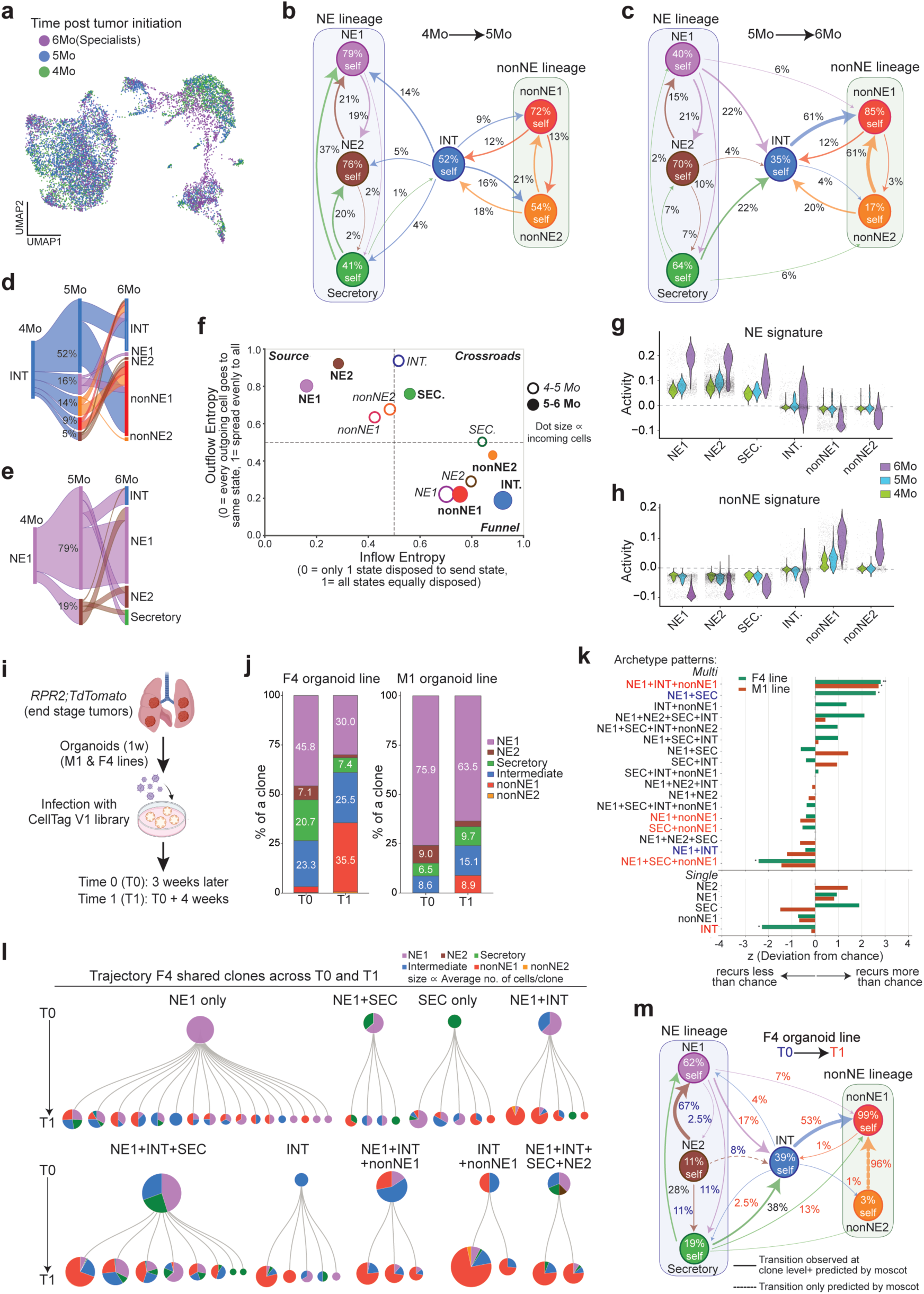
Temporal and clonal analysis of SCLC cell-state transitions. **a.** Uniform manifold approximation and projection (UMAP) of combined scRNA-seq data from sorted cancer cells from 4-, 5- and 6-month *RPR2* tumours post initiation (only “Specialist” archetype cells are shown for the 6-month timepoint). **b,c.** moscot transition couplings for the 4–5-month (b) and 5–6-month (c) intervals, with archetypes grouped into NE (NE1, NE2, Secretory) and nonNE (nonNE1, nonNE2) lineages and Intermediate between them. Percentage values are the fraction of the source state’s cells reaching each destination (node self-renewal + outgoing arrows = 100%), as determined by moscot. Transitions shown above 1%, or ≥ 5% for cross-lineage arrows. **d,e.** Alluvial flow plots from 4-month Intermediate (d) and NE1 (e) cells. Ribbon width is proportional to cell mass, and percentages at 5-month show the fraction of the source’s 4-month cells taking each route; routes below 5% are omitted at both hops. **f.** Log-normalised Shannon diversity (self-transitions excluded) of inflow (x) versus outflow (y) diversity per state. Open circles, 4–5-month interval; filled circles, 5–6-month interval. **g,h.** Violin plots comparing the activity of the NE gene signature (g) and nonNE gene signature (h) in the *RPR2* cancer cells profiled across the 4-, 5- and 6-month timepoints. **i.** Schematic of the CellTag timepoint experiment in *RPR2* organoid lines F4 and M1. **j.** Stacked bar plots comparing the archetype composition across timepoints for organoid lines F4 and M1. **k.** Recurrence of archetype combinations within CellTag clones in F4 and M1. Bars represent the deviation of the observed number of clones carrying each combination from chance expectation, in standard deviations (z); combinations to the right are enriched, those to the left disenriched. Asterisks denote significance (* p < 0.05, ** p < 0.01, *** p < 0.001). **l.** F4 clones shared across timepoints T0 and T1 (57 clones), grouped by the set of archetypes they contained at T0 and ordered left to right by the proportion of nonNE cells they reached at T1; groups represented by a single clone are omitted. The upper pie of each group gives the pooled composition of its founding clones, and each lower pie one individual clone at T1. Pie area is proportional to cell number at T0 but drawn on a compressed scale at T1, so that clones of only a few cells remain visible. **m.** Cell-state flux in F4 organoids, combining moscot-inferred transitions with the CellTag lineage evidence. Every percentage is moscot flux, i.e. the fraction of the source state’s T0 cells reaching that destination. Line style carries the clonal evidence separately: solid where barcoded lineages witnessed the edge, and broken arrow where only moscot predicts it. Label colour indicates whether that clonal evidence came mainly from T0 (blue), mainly from T1 (red), or both (black).

During the 4–5-month interval, the Intermediate state was by contrast the least stable: roughly half of its cells remained Intermediate, while the half that departed split almost evenly between the NE and nonNE lineages (48% toward NE versus 52% toward nonNE) (**Fig. 2b-d**). Thus, in early tumours, the Intermediate state functions as a “crossroad” state, through which predominantly nonNE cells pass before transitioning to NE, remaining Intermediate, or returning to nonNE (**Fig. 2d** and **Supplementary Fig. 7c,d)**. Outgoing Intermediate cells fated for NE versus nonNE lineage were transcriptionally distinct: Intermediate-to-NE cells upregulated neuronal programs, while Intermediate-to-nonNE cells strongly upregulated immune-related signalling (**Supplementary Fig. 7e**).

During the 5–6-month interval, the outgoing flux of Intermediate cells became markedly directional, with most cells resolving toward nonNE (60.8%) and essentially none returning to NE (0.5%) (**Fig. 2c,d,f**). Despite this directed outflow, all other archetypes contributed cells to the Intermediate state during this period, and the inflow was dominated by the NE and Secretory states rather than nonNE (79% versus 21% of cells entering Intermediate). The Intermediate state therefore switches from a crossroad to a funnel during tumour progression, channelling cells progressively toward nonNE as tumours advance, while the NE and nonNE states consolidate as self-maintaining endpoints (**Fig. 2f**). Consistent with this consolidation, scoring NE and nonNE gene signatures across the combined dataset showed that the two opposing lineages became mutually exclusive with progression, with each set of lineage genes being more strongly expressed in its own archetype and more repressed in the other by end stage (**Fig. 2g,h**). As before, Intermediate cells transitioning to nonNE states upregulated the same inflammatory and immune-related programs (**Supplementary Fig. 7f**). Interestingly, NE cells that moved to the Intermediate state downregulated neuronal gene signatures and upregulated translational regulation and ribosome biogenesis alongside some interferon-related inflammatory programs (**Supplementary Fig. 7g**), thus implicating these transcriptional changes as early events mediating the NE to nonNE lineage switch.

### Lineage tracing establishes the Intermediate state as an obligate route between NE-to-nonNE

To experimentally validate the cell state transitions described above, we performed lineage tracing in organoids derived from end-stage *RPR2* tumours, which retain natural variation in lineage composition (**Fig. 2i**). We transduced two organoid lines, F4 and M1, with the CellTag barcoding system ^41^ and profiled them by scRNA-seq at T0 (3 weeks after transduction and selection) and T1 (one month later), assigning archetypes by anchor-based label transfer from the end-stage tumour dataset as described above. We recovered 232 clones in F4 and 441 in M1, including 57 F4 and 172 M1 clones shared across the two timepoints. Consistent with the initial flow cytometry, F4 was more heterogeneous than M1 (**Fig. 2j**) and had fewer single-archetype or single-cluster clones at both timepoints (**Supplementary Fig. 7h,i**). In both lines, the NE1+Intermediate+nonNE1 combination was over-represented, whereas cross-lineage clones lacking Intermediate (NE1+nonNE1 and NE1+Secretory+nonNE1) were under-represented, suggesting that passing through the Intermediate state may be the “preferred” route in the NE to nonNE lineage transition (**Fig. 2k**). To test whether clones containing Intermediate cells at T0 progress further toward nonNE, we grouped the shared clones of each line by their T0 composition and measured what fraction of each group’s cells had become nonNE by T1 (**Fig. 2l** and **Supplementary Fig. 7j**). In F4, founder clones containing Intermediate cells at T0 reached a substantially higher proportion of nonNE cells at T1 than those without (> 43% versus < 30%). NE1+Intermediate founder clones reached the highest proportion (84%). The same relationship held in M1, although nonNE cells remained below 10% in all clones (**Supplementary Fig. 7j**). Thus, the presence of Intermediate cells in a founder clone predicted how far that clone progressed toward nonNE, while the magnitude of progression was set by the heterogeneity of these organoid lines.

Combining clonal evidence with moscot-inferred transition probabilities revealed that the transition structure in the F4 organoid closely recapitulated that observed *in vivo* between 5 and 6 months: the Intermediate state showed some self-renewal (39.2% versus 34.9% *in vivo*) and directed its output predominantly towards nonNE1 (53.5% versus 60.8%), while returning almost nothing to NE1 (**Fig. 2m**). Cells in the NE1 state in turn fed the Intermediate state at a comparable rate in F4 and *in vivo* (17.0% versus 21.9%), and nonNE1 cells behaved as a near-absorbing state in each (99.5% versus 85.3%) (**Fig. 2m and 2c**). The M1 organoid departed from this pattern: its NE1 compartment had reduced outflux and generated far fewer Intermediate cells, while those that did arise were more self-renewing **(Supplementary Fig. 7k**). Intermediate output was inverted, returning more cells to NE1 than to nonNE1 (19.3% versus 7.0%) (**Fig. 2m and Supplementary Fig. 7k**). In M1, the NE1-to-Intermediate-to-nonNE1 axis is thus attenuated at every step, indicating that the extent of nonNE conversion depends on flux through Intermediate.

Together, our *in vivo* and *ex vivo* analyses identify the Intermediate state as a low-identity state and the principal route between the NE and nonNE lineages.

### Intermediate cells exist in several mouse SCLC contexts

We next asked whether the Intermediate state exists in SCLC contexts with altered NE-nonNE dynamics, such as *RPR2* liver metastases, which lack nonNE cells ^22^. We performed single-cell multi-omics on sorted cancer cells from these metastases and scored archetype signatures compiled from primary tumours (**Fig. 3a**). The NE2 signature showed high but variable activity across all liver clusters (**Fig. 3b**). In contrast, no cluster showed activity for the nonNE1 signature, consistent with an absence of the nonNE lineage (**Supplementary Fig. 8a**). The inflamed basal signature (Ireland *et al.*, ^12^), high in nonNE2 cells from primary tumours, was likewise absent (**Supplementary Fig. 8b**). The Intermediate signature was active specifically in cluster 6 (**Fig. 3c**), which also showed wide variance in NE2 score, consistent with the low-NE character of these cells (**Fig. 3b**). Top markers of cluster 6 included Intermediate genes (**Supplementary Fig. 8c**), and every pathway upregulated in this cluster was shared with the Intermediate archetype, including ribosome biogenesis, translational control, nonsense-mediated decay, antigen presentation and TNFα-NFκB signalling (**Fig. 3d**). Thus, the Intermediate state persists even in purely NE tumours that lack nonNE cells and do not undergo lineage transition.

**Figure 3:**
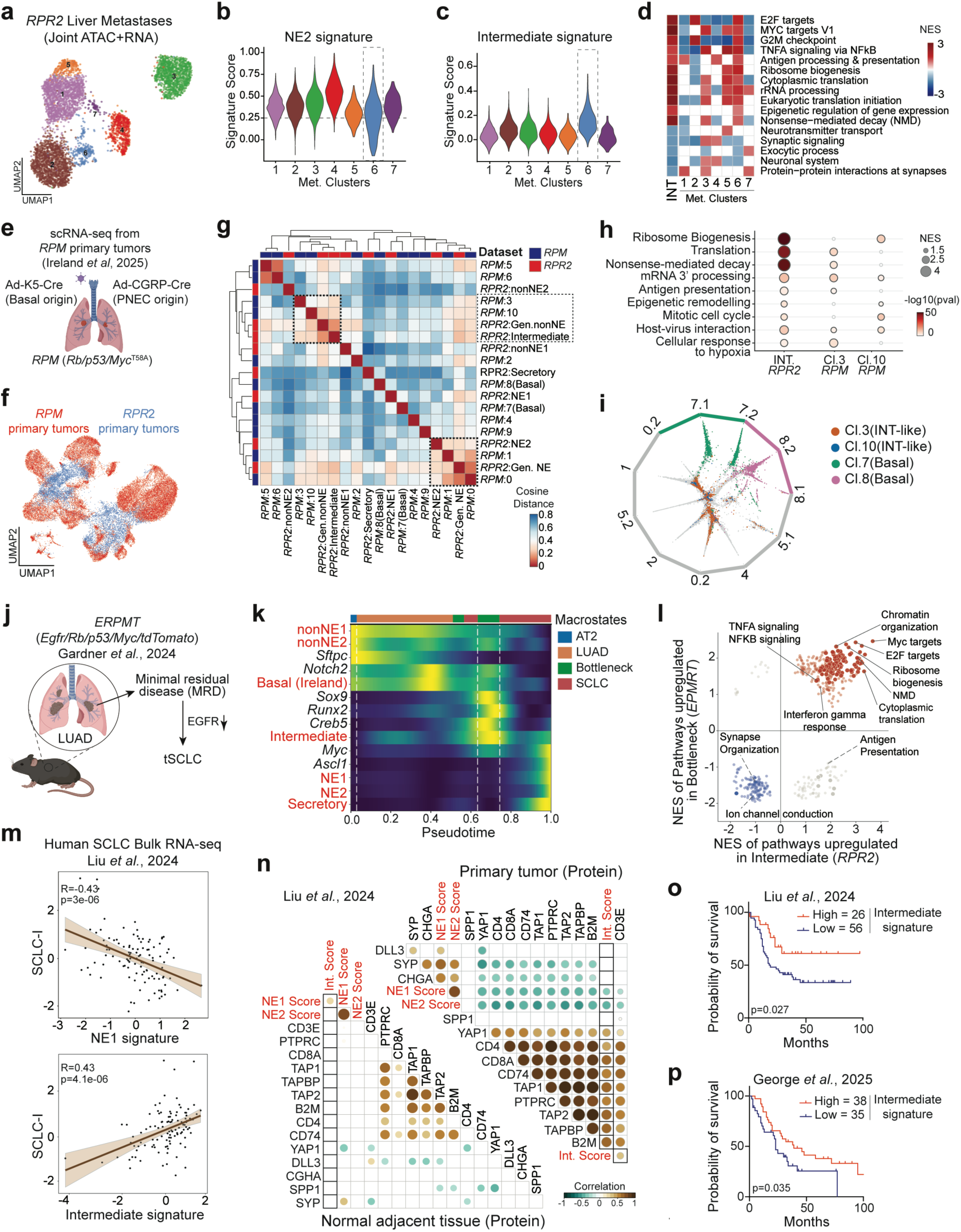
Conservation of the Intermediate archetype across mouse models and human tumours. **a.** Uniform manifold approximation and projection (UMAP) of single-cell ATAC-seq and RNA-seq data from cancer cells sorted from *RPR2* liver metastases, coloured by Leiden cluster. **b,c.** Violin plots of NE2 (b) and Intermediate (c) archetype signature activity across liver metastasis clusters. Box highlights cluster 6, the only cluster with high Intermediate signature activity and a broad spread of NE2 scores. **d.** Heatmap of differential enrichment of Intermediate-specific pathways across the seven liver metastasis clusters, with the Intermediate column from the primary tumours as reference. Cells are labelled with the normalised enrichment score. Cluster 6 is the only cluster in which all Intermediate-upregulated pathways are concordantly enriched. INT, Intermediate**. e.** Schematic of the experimental design from Ireland *et al*. for generating scRNA-seq data from primary tumours with different cells of origin. Adeno-K5-Cre initiates primary SCLC tumours from basal cells in the lung and Adeno-CGRP-Cre initiates tumours from pulmonary neuroendocrine cells (PNECs). **f.** Uniform manifold approximation and projection (UMAP) of canonical correlation analysis (CCA)-integrated scRNA-seq data from *RPM* and *RPR2* primary tumours, the latter restricted to 6-month cancer cells. **g.** Hierarchical clustering of pairwise cosine distances between every *RPM* cluster and *RPR2* archetype in the integrated CCA space, where smaller distances indicate greater similarity. The Intermediate archetype clusters with *RPM* clusters 3 and 10 (highlighted). **h.** Dot plot of enriched pathways shared between *RPR2* Intermediate cells and *RPM* clusters 3 and 10. **i.** Circular projection of *RPM* primary tumour single cells according to their fate probabilities. Fate-biased cells are placed next to their corresponding colours, while lineage-naïve cells are placed in the middle. *RPM* clusters 3 and 10 (orange and blue) sit predominantly at the centre of the plot, whereas the basal clusters 7 and 8 occupy the edges and contribute to distinct macrostates. **j.** Experimental design from Gardner *et al.* modelling the LUAD-to-SCLC lineage transition: switching off the oncogenic *Egfr* transgene after lung adenocarcinoma (LUAD) has developed in the *ERPMT* mouse model drives tumour regression to minimal residual disease (MRD), followed by transformation into SCLC (tSCLC). **k.** Heatmap of scaled activity score of *RPR2* archetype signatures and imputed expression of lineage markers along terminal cell-state probabilities computed with CellRank for all cells. Intermediate signature activity peaks at the “bottleneck” state, coinciding with expression of *Creb5* and *Runx2*, markers of the MRD state defined in Gardner *et al*. **l.** Scatter plot comparing pathways upregulated (red) and downregulated (blue) in *ERPMT* bottleneck cells with those in Intermediate cells of end-stage *RPR2* tumours. **m.** Plots showing the Pearson correlation between the human inflamed SCLC signature and the *RPR2* NE1 signature (top) and the Intermediate signature (bottom) in individual primary tumour samples from the Tongji University (TU) cohort (n = 112), described in Liu *et al*. **n.** Correlation matrices showing associations between Intermediate, NE1, and NE2 signatures and protein levels of antigen presentation genes, immune cell markers, and T cell markers in primary tumours (top) versus normal adjacent tissue samples (bottom) from the TU SCLC cohort. Significant pairwise Pearson correlations (p < 0.01) are highlighted. **o,p.** Kaplan-Meier curves for overall survival based on Intermediate state signature activity in limited-stage patients from the TU-SCLC cohort (o) and in extensive-stage patients from the George *et al.* cohort (p). P-values were determined through log-rank test.

To evaluate whether an Intermediate-like state exists in other SCLC mouse models, we re-analysed published scRNA-seq of autochthonous *RPM* (*Rb/p53/MycT58A*) primary tumours initiated from basal cells (Adeno-K5-Cre) or pulmonary neuroendocrine cells (PNECs, Adeno-CGRP-Cre) (**Fig. 3e**); this model displays broader transcriptional heterogeneity compared to the *RPR2* model ^12, 24^. We integrated the end-stage *RPR2* and *RPM* primary tumour datasets by canonical correlation analysis (CCA) and measured the cosine distance between each archetype and *RPM* cluster (**Fig. 3f**). *RPR2* NE cells co-localised with the NE-high *RPM* population, confirming that the approach detects genuine equivalence (**Supplementary Fig. 8d**). The Intermediate state was closest to *RPM* clusters 3, 10 and 0, and most distant from the basal-like clusters 7 and 8 (**Fig. 3g** and **Supplementary Fig. 8e**). Cluster 0, the largest, was also proximal to NE1 and NE2, so we focused on clusters 3 and 10, which were uniquely proximal to Intermediate. GSEA showed that these two clusters were enriched for all programs that define the Intermediate state, including translation and ribosome biogenesis, antigen presentation, chromatin remodelling, and cell cycle (**Fig. 3h**). When we implemented CellRank on the combined K5-Cre/CGRP-Cre *RPM* datasets, only clusters 3 and 10 failed to form or contribute significantly to terminal macrostates (representing stable differentiation endpoints), indicating that they comprise low-commitment, transitional cells (**Fig. 3i** and **Supplementary Fig. 8f**). By comparison, basal cells in clusters 7 and 8 reliably formed dedicated terminal macrostates, irrespective of the cell of origin. These analyses independently identify undifferentiated, low-lineage cell states in *RPM* tumours that recapitulate Intermediate cell state.

Therapy resistance in LUAD can occur through transformation to SCLC (tSCLC) ^42^. We re-analysed published scRNA-seq from a mouse model where tSCLC emerges from LUAD after minimal residual disease (MRD) upon EGFR downregulation ^36^ (**Fig. 3j)**. As expected, normal alveolar type 2 (AT2) and LUAD cells had high nonNE archetype activity, and tSCLC cells scored high for NE1/NE2 and Secretory signatures; strikingly, the Intermediate signature peaked at MRD (**Supplementary Fig. 8g**). Trajectory analysis uniquely placed peak Intermediate activity in the “bottleneck” cells, which emerge during MRD, while other archetype and SCLC state signatures peaked either prior to entering or after exiting the bottleneck (**Fig. 3k**). Pathway enrichment analysis of these bottleneck cells at MRD revealed near-complete concordance with the Intermediate state, except for antigen presentation (**Fig. 3l**). These data indicate that during the LUAD-SCLC lineage transition, cancer cells pass through an Intermediate-like bottleneck state characterised by the recurrent transcriptional program comprising translational regulation, cell cycle, epigenetic regulation and TNFα signalling.

Collectively, these analyses establish Intermediate cells as a conserved cancer cell state implicated in cell-state transitions across several mouse models.

### The Intermediate signature predicts immune infiltration in human SCLC

To assess the relevance of the Intermediate state identified in human SCLC, we applied *RPR2* archetype signatures to bulk RNA-seq from 112 treatment-naïve SCLC tumours with paired normal adjacent tissue (TU-SCLC cohort, Liu *et al.*, ^19^). The NE1/2, Secretory, and Intermediate signatures showed significantly higher scores in tumours versus normal lung; in contrast, the nonNE1/2 signatures were not cancer-specific (**Supplementary Fig. 9a**). A subset of human SCLC-I tumours (“SCLC-I-NE” subtype) shows improved response to combined chemotherapy and immunotherapy ^8, 37^. NE1/2 and Secretory signatures were strongly anti-correlated with SCLC-I-NE activity, but Intermediate and nonNE signatures showed positive correlation, with the Intermediate signature displaying the strongest association (**Fig. 3m** and **Supplementary Fig. 9b**). Intermediate signature activity was strongly positively correlated with the expression of antigen presentation and immune/T cell genes, negatively correlated with ASCL1 and NEUROD1, and weakly positively correlated with POU2F3 (**Supplementary Fig. 9c**). NE1 and NE2 signatures displayed reciprocal strong negative correlations with immune genes, consistent with NE tumour immunosuppression (**Supplementary Fig. 9c**). These associations were tumour specific and absent in normal lung tissue (**Supplementary Fig. 9c**). Proteomics data from tumour samples from the same cohort confirmed that elevated Intermediate signature activity predicted increased protein levels of antigen presentation machinery and T-cell infiltration, exclusively in tumour samples (**Fig. 3n**).

In an independent cohort of 65 patients with extensive-stage primary and metastatic tumours (George *et al.*, ^43^), Intermediate signature activity correlated positively with immune and antigen-presentation genes and negatively with NE markers, with NE1/NE2 signatures showing the reciprocal pattern (**Supplementary Fig. 9d**). Notably, in both cohorts, patients whose tumours had a high Intermediate signature had better overall survival (**Fig. 3o,p**), while the opposite was true for the NE2 signature (**Supplementary Fig. 9e**) and variable results were observed for the nonNE1 signature (**Supplementary Fig. 9f**). Thus, Intermediate signature activity is cancer-specific and predicts enhanced immune infiltration and antigen presentation across both limited- and extensive-stage SCLC.

### Intermediate cells exhibit a permissive chromatin state

To determine the epigenetic basis of SCLC cell plasticity, we used the scATAC-seq modality of the end-stage multi-omics dataset and identified differentially accessible regions across archetypes. This analysis revealed ∼10,000 variable *cis*-regulatory elements in two non-overlapping clusters by activity in NE versus nonNE lineages, underscoring the epigenetic barrier between these states ^23^ (**Fig. 4a**). Notably, Intermediate cells exhibited moderate chromatin accessibility at *cis*-regulatory elements of both the NE and nonNE lineages at the population level (**Fig. 4a**) and in individual cells (**Supplementary Fig. 10a**). This permissive chromatin architecture was evident at lineage-defining loci: the NE gene *Calca* and the nonNE gene *Notch2* each maintained lineage-restricted accessibility in their cognate populations, yet both genes were accessible in Intermediate cells (**Fig. 4b**). In contrast, Intermediate cells did not show accessibility at constitutively open scATAC-seq peaks (**Supplementary Fig. 10b**), indicating that this openness is restricted to the regulatory elements implementing NE and nonNE identity.

**Figure 4:**
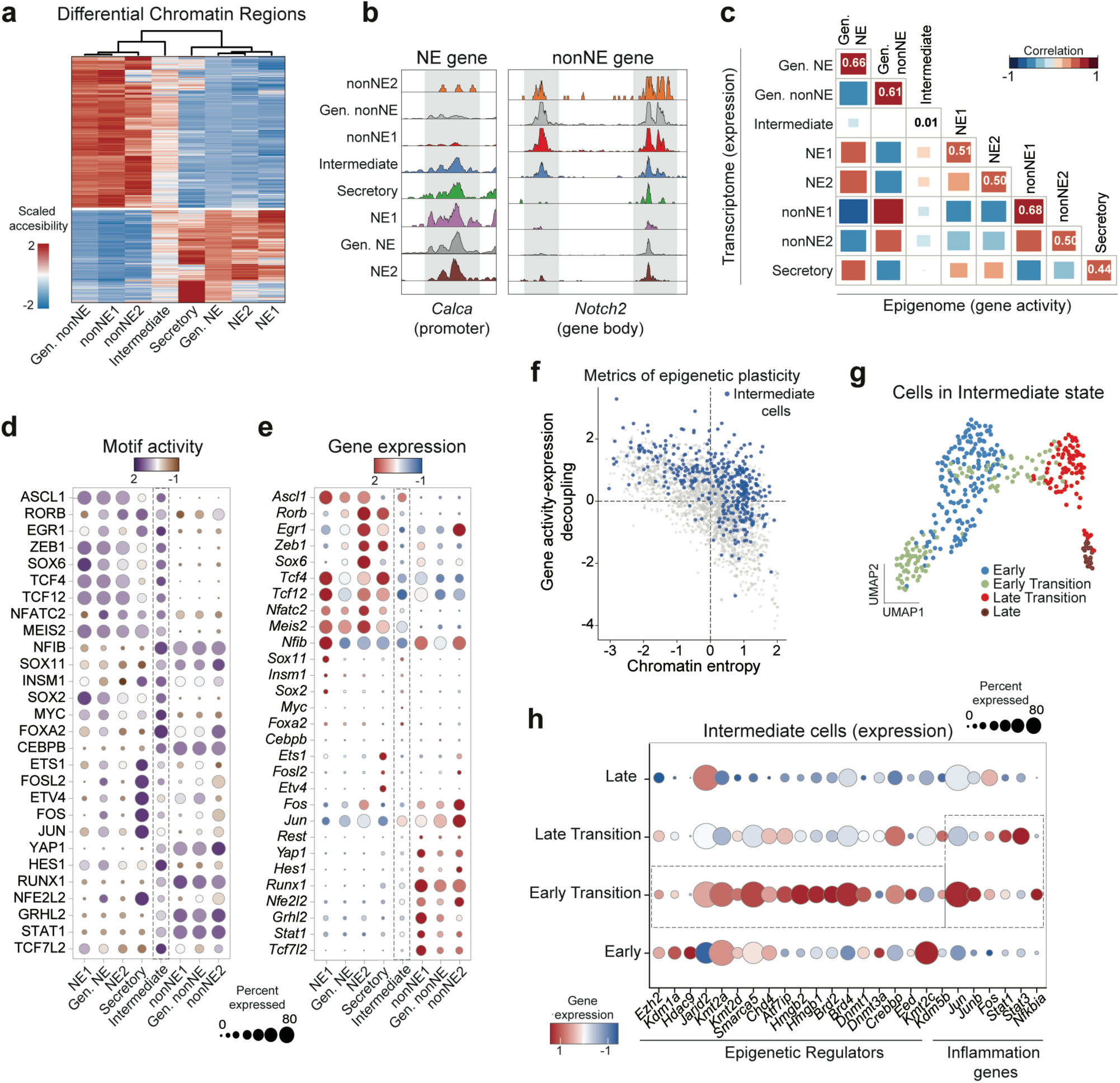
SCLC cells in the Intermediate state exhibit permissive chromatin accessibility and epigenetic plasticity. **a.** Heatmap of scaled accessibility for highly variable chromatin regions across *RPR2* cancer cell archetypes. **b.** Chromatin accessibility at the promoter of the NE gene *Calca* (left) and at the gene body of the nonNE gene *Notch2* (right) across archetypes. **c.** Heatmap showing Pearson correlations between gene expression and chromatin-based gene activity scores of all highly variable marker genes in each archetype. **d,e.** Dot plots comparing the motif activity (d) and RNA expression levels (e) of top variable transcription factors in *RPR2* archetypes. **f.** Scatter plot showing the gene activity-expression decoupling score and chromatin entropy in individual *RPR2* cells. Intermediate cells, shown in blue, predominantly occupy the upper-right quadrant and are high for both measures of chromatin plasticity. **g.** UMAP visualisation of the scRNA-seq modality of Intermediate cancer cells coloured by pseudotime-based cluster identity. **h.** Expression of selected differentially regulated genes coding for epigenetic regulators and inflammation-related transcription factors (TFs) across Intermediate cell pseudotime clusters as in (g).

Correlating gene activity scores (scATAC-seq) with mRNA expression (scRNA-seq) across all differential genes showed strong coupling in every archetype except Intermediate, where the two were uncorrelated (**Fig. 4c**). The same split held at the level of transcription factors (TFs): in every differentiated archetype, TF motif activity matched the expression of the top variable TFs, whereas Intermediate cells uniquely retained open motifs for both NE and nonNE TFs despite no detectable expression of these TFs in this state (**Fig. 4d,e**). To quantitatively assign an epigenetic plasticity score to cells, we combined (1) Shannon entropy of chromatin accessibility, and (2) discordance between accessibility and gene expression (see Methods). Intermediate cells displayed significantly elevated plasticity scores compared to all differentiated archetypes, both at the single-cell and population levels (**Fig. 4f** and **Supplementary Fig. 10c,d**), confirming that their chromatin landscape is more permissive than that of cells in the NE and nonNE lineages. In liver metastases, the Intermediate-equivalent cluster 6 showed the same pattern, with high motif activity for both NE and nonNE TFs but no corresponding RNA expression (**Supplementary Fig. 10e,f),** indicating that permissive chromatin at lineage-specific regions is an intrinsic feature of the Intermediate state.

To understand how chromatin resolves as cells traverse the bottleneck between the NE and nonNE lineages, we subclustered Intermediate cells by Palantir pseudotime into four clusters: “early,” “early-transitioning,” “late-transitioning,” and “late” (**Fig. 4g** and **Supplementary Fig. 11a**). Differential gene expression analysis revealed an upregulation of several epigenetic regulators in the “early-transitioning” Intermediate cells including *Kmt2a*, *Atf7ip*, *Jarid2*, *Brd2*, *Hmgb2, Smarca4*, and *Dnmt1* (**Fig. 4h**). Consistently, these cells also showed elevated expression of a broad epigenetic regulation and chromatin organisation signature (**Supplementary Fig. 11b**), suggesting that active epigenetic reprogramming initiates lineage conversion. Pseudotime-guided analysis with Monocle3 ^44^ further identified distinct gene modules across the four clusters (**Supplementary Fig. 11c,d**). Modules enriched in “early-transitioning” cells included TNFα-NFκB signalling genes, further implicating inflammation in SCLC lineage transition (**Supplementary Fig. 11e,f**). Accordingly, pseudotime progression within the early-transitioning cluster, but not within the other Intermediate clusters, showed a strong positive correlation with the inflammation score associated with SCLC-I-NE human tumours (**Supplementary Fig. 11g**).

Together, these findings support a model in which Intermediate cells maintain a more plastic chromatin landscape characterised by high entropy and low transcription-chromatin concordance, providing a molecular mechanism underlying the ability of SCLC cells to switch lineage.

### Multi-omics analysis defines gene regulatory networks in SCLC

Epigenetic regulation impinges upon TF networks to promote lineage plasticity ^45^. In SCLC, the regulators that read and reinforce chromatin states remain incompletely defined. Having characterised the epigenetic landscape of intratumoural heterogeneity in SCLC, we next sought to identify these regulators and map the gene regulatory networks governing SCLC cell states, using the FigR (Functional Inference of Gene Regulation) approach ^46^ (**Fig. 5a**). Linking scATAC-seq peaks to putative target genes revealed significant associations between 25,112 *cis*-regulatory elements and 9,747 genes (permutation p < 0.05) across all *RPR2* cancer cells. We then identified 1,269 domains of regulatory chromatin (DORCs), i.e. genes with ≥ 5 peak-gene links (**Fig. 5b**). These high-density regions are enriched for lineage-specific genes in multiple contexts ^47, 48^ and hence could be associated with specific cell states in SCLC. Indeed, the DORCs included known markers of the NE and nonNE cell states, as well as hundreds of genes from SCLC-relevant pathways, including axon guidance, endocrine secretion, synapse formation, and Hippo signalling (**Fig. 5b** and **Supplementary Fig. 12a**).

**Figure 5:**
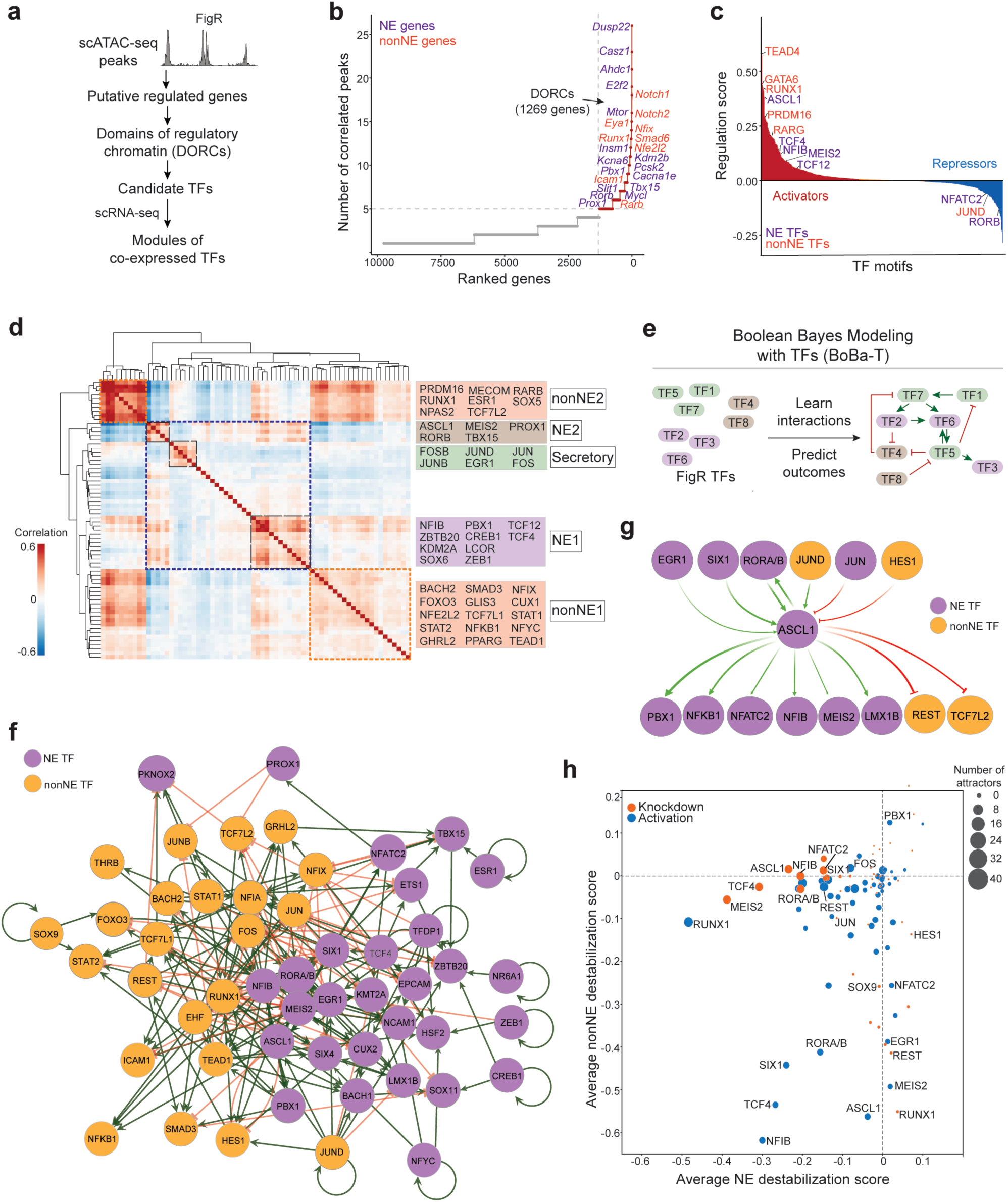
Network analysis reveals key transcription factors controlling SCLC cell states. **a.** Schematic of the FigR package workflow followed for scATAC-seq data analysis of SCLC cells in *RPR2* mutant tumours. **b.** Top domains of regulatory chromatin (DORCs) ranked by significant gene-peak correlations across all cancer cell archetypes in *RPR2* tumours; purple points represent NE lineage genes and orange points mark nonNE genes. **c.** Mean regulation score for each transcription factor (TF, n = 870) across DORCs (n = 1,269), distinguishing TF activators from repressors based on distribution. **d.** Heatmap showing correlation among the 64 TFs identified as DORC regulators (absolute regulation score ≥ 1.25); NE modules (blue rectangles) and nonNE modules (yellow rectangles) are marked, with corresponding cancer cell archetypes labelled. **e.** Schematic of the BoBa-T algorithm workflow. **f.** Regulatory network of the 64 TFs; colours indicate NE and nonNE lineages, with green lines for activating and red for inhibitory interactions. **g.** Top upstream regulators and downstream targets of ASCL1 as predicted by BoBa-T. **h.** Scatter plot of average destabilisation scores for NE and nonNE cell states following simulated TF knockdown (orange) or activation (blue), as predicted by BoBa-T. The size of the dots indicates the number of stable network states or attractors disrupted by *in silico* perturbation of each factor.

Motif enrichment then identified TFs whose binding sites were over-represented in DORC *cis*-regulatory elements and whose expression correlated with their putative targets. This revealed 64 “driver” TFs, with top activators being ASCL1 and TEAD1, which are established lineage drivers of NE and nonNE cell fates in *RPR2* tumours ^23^, as well as TFs not previously implicated in SCLC biology (**Fig. 5c**). For individual cell-state-specific genes such as Insm1 (NE) and Notch2 (nonNE), expression was predicted to be orchestrated by both known lineage determinants (ASCL1 and TEAD1, respectively) and TFs uncharacterised in SCLC (e.g., PRDM5, IRF6, and HES6) (**Supplementary Fig. 12b,c**).

To determine how these TFs may function together, we correlated their expression in individual cells, revealing five modules of co-expressed TFs, three comprising NE TFs and two nonNE TFs (**Fig. 5d**). The first NE module contained known promoters of NE tumours such as ASCL1 and PROX1 ^49^, as well as TFs not previously associated with the NE state (e.g., TBX15). The second NE module included NFIB and ZEB1, implicated in epithelial-mesenchymal transition ^50^, suggesting a role in metastatic progression. The third NE module comprised AP-1 TFs (e.g., c-JUN, c-FOS) that were negatively correlated with the ASCL1 module. Among the two nonNE modules, one included TFs such as TEAD1 and STAT1 that regulate nonNE identity and inflammation ^32, 51^, whereas the other comprised RUNX1-correlated TFs that broadly regulate angiogenesis and may be linked to vasculogenic mimicry in SCLC ^52^. Notably, these five TF modules mapped onto cancer cell archetypes identified by scRNA-seq (**Fig. 5d**, see **Fig. 1b**), even though the regulatory modules were compiled through an archetype-agnostic analysis. Only the Intermediate archetype lacked a dedicated TF module, likely because the epigenetic changes that accompany the establishment of this state require coordination among multiple modules.

### Transcription factor networks reveal deterministic regulators of SCLC heterogeneity

To determine how these modules interact to establish cellular lineage, we adapted a gene regulatory network-inference algorithm, BooleaBayes, to delineate interactions between the 64 FigR-identified TFs and key cell-type markers (NCAM1, ICAM1, EPCAM), and to infer regulatory rules governing archetype stability (**Fig. 5e**). Previously, BooleaBayes was applied to TF networks controlling SCLC subtypes using human bulk RNA-seq data ^38, 53^. The new Boolean Bayes model for transcription factor networks (BoBa-T) adds (1) expansion to a more comprehensive TF network that incorporates cell-type-specific TF interactions derived from paired ATAC-seq data, (2) training on high-resolution single-cell data, (3) robust quantification of regulatory rule metrics, and (4) expanded TF perturbation analyses. Briefly, BoBa-T infers a TF network from the multiomics data and simulates loss- and gain-of-function of individual TFs to predict drivers of each SCLC cell state (see Methods). Each TF-target relationship is returned as activating, inhibiting or non-interacting (**Fig. 5f**). We validated regulatory relationships between TFs using a train-test split on the single-cell data. BoBa-T predicted TF expression accurately across all network factors, with an average ROC-AUC of 0.97 and consistently high F1 scores (**Supplementary Fig. 12d-f**). For individual TFs, such as ASCL1, the predicted single-cell expression was highly concordant with actual normalised expression in the test scRNA-seq data (correlation > 0.96) (**Supplementary Fig. 12g,h**). At the level of TF interactions, BoBa-T also accurately captured known regulatory hierarchies. Analysis of the ASCL1 node, for instance, identified HES1 as an upstream repressor and NFIB among its downstream targets, which are experimentally validated regulators and targets of ASCL1 (**Fig. 5g**).

To identify TFs that may be critical for archetype stability, we conducted *in silico* perturbations where individual factors were either permanently activated (TF = 1) or silenced (TF = 0). TFs whose silencing or activation caused more than 20% destabilisation (score ≤ −0.25) of specific archetypes, were classified as master stabilisers and destabilisers, respectively. Consistent with prior experimental evidence ^23, 54^, ASCL1 was a predicted stabiliser of the NE state (and a destabiliser of the nonNE state), while HES1 and REST were stabilisers of the nonNE state (and destabilisers of the NE state) (**Fig. 5h**). Importantly, in addition to expected regulators of SCLC cell states, the network simulations identified several previously uncharacterised candidate regulators of SCLC archetypes such as SIX1, NFATC2, ROR factors and EGR1 (**Fig. 5h**).

### RORB is a novel regulator of the NE state

Since NE features are often associated with resistance to therapy, including immunotherapy ^5^, we prioritised identifying TFs whose downregulation could destabilise differentiated NE states. The retinoic acid-related orphan nuclear receptor paralogues RORA/RORB emerged as compelling candidates: BoBa-T simulations predicted that the knockdown of these factors could destabilise classical NE states (NE2) as effectively as ASCL1, while their upregulation would destabilise low/nonNE states (**Fig. 5h** and **Supplementary Fig. 12i,j**). Network analysis further indicated that RORA/B function as both activators and repressors and formed the only positive feed-forward loop with ASCL1 among NE factors (**Supplementary Fig. 13a**). These predictions indicated that loss of RORA/B would shift ASCL1-positive cells toward low-NE phenotypes, implicating these factors as candidate gatekeepers of the NE lineage.

To distinguish between paralogues, we analysed RORA/RORB expression across mouse lung cancer models. In *RPR2* primary tumours, *Rora* and *Rorb* had overlapping expression and were restricted to classical NE states (NE2) and absent from Intermediate and nonNE states; no other RAR/ROR TFs were significantly expressed in NE cancer cells, though *Rarb* was expressed in some nonNE states (**Supplementary Fig. 13b**). Similarly, in *RPM-RPMA* organoid-derived tumours, *Rorb* had higher expression than *Rora,* and its expression was restricted to NE and neuronal cancer cells, while *Rora*, alongside NE cells, also displayed strong expression in basal cells (**Supplementary Fig. 13c**). Across published transcriptomes of mouse models of lung cancer ^55^, *Rora* showed broad expression, whereas *Rorb* expression was restricted to SCLC tumours (**Supplementary Fig. 13d**). This specificity was confirmed in LUAD-to-SCLC data, where *Rora* was expressed in all cell types while *Rorb* expression only emerged in tSCLC cells (**Supplementary Fig. 13e**). In human tumours, *RORB* expression was confined to SCLC-A tumours, while *RORA* was expressed uniformly across all subtypes (**Supplementary Fig. 13f**). Consistent with this, in the TU-SCLC cohort, *RORB* expression was greatly elevated in immune-cold tumours with high NE scores compared to immune-hot tumours (**Supplementary Fig. 13g**).

To test whether elevated levels of RORB promote NE identity in a heterogeneous SCLC population, we overexpressed RORB in *RPR2* tumour organoids, which comprise both NE and nonNE cancer cells, characterised by the expression of cell surface proteins NCAM1 and ICAM1, respectively^23^ (**Fig. 6a**). Elevation of *Rorb* expression led to strong upregulation of NE markers *Ascl1*, *Chga, Calca,* and *Dll3* (**Supplementary Fig. 14a**), alongside an increase in the percentage of NCAM1^+^ NE cells (**Fig. 6b,c**). These observations support the *in-silico* prediction that RORB regulates NE identity.

**Figure 6:**
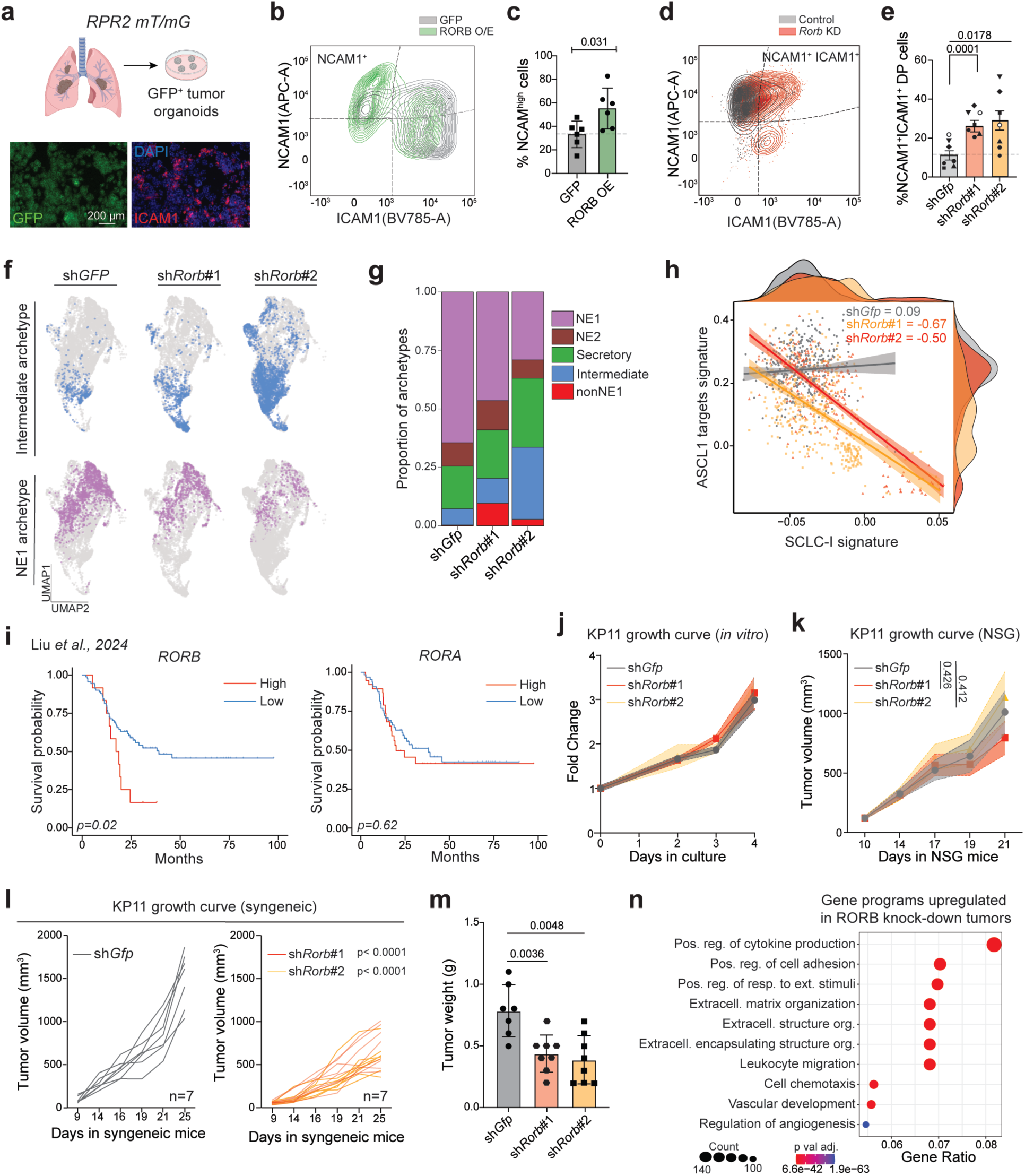
RORB loss destabilises neuroendocrine identity and enhances tumour immunogenicity. **a.** Experimental schematic for generating tumour organoids from *RPR2 mT/mG* lung tumours (normal cells are tdTomato^+^ while Cre-recombined cancer cells are GFP^+^). Representative immunofluorescence images show GFP^+^ tumour organoids (green) co-stained with ICAM1 (magenta) and DAPI (blue). Scale bar, 200 µm. **b.** Representative overlaid flow cytometry plots showing NCAM1 and ICAM1 surface expression in control (grey, GFP) and *Rorb* overexpressing (green) *RPR2* tumour organoids. **c.** Quantification of the percentage of NCAM1^+^ neuroendocrine (NE) cells in Control (grey) and *Rorb* overexpressing organoids, as determined by flow cytometry (n = 5 biological replicates). Error bars represent the s.e.m. P-values were determined by paired t-test. **d.** Representative overlaid flow cytometry plots showing NCAM1 and ICAM1 surface expression in control (grey, sh*Gfp*) and *Rorb* knockdown (red) *RPR2* tumour organoids. **e.** Quantification of NCAM1^+^ ICAM1^+^ double-positive (DP) Intermediate cells in organoid lines following *Rorb* knockdown (n = 7 biological replicates). Error bars represent the s.e.m. P-values were determined by paired t-tests comparing each *Rorb* shRNA to control sh*Gfp*. **f.** UMAP projections of control and *Rorb* knockdown organoid scRNA-seq dataset showing Intermediate (top, blue) and NE1 (bottom, mauve) archetype cells in each condition. **g.** Stacked bar plot showing the proportion of *RPR2* archetype assignments (NE1, NE2, Secretory, Intermediate, nonNE1) across control and *Rorb* knockdown organoid samples. **h.** Scatter plot with marginal density distributions showing the relationship between ASCL1 target gene activity (x-axis) and human SCLC-I inflamed signature activity (y-axis) in single cells from control and *Rorb* knockdown organoids. Note the anti-correlation and emergence of SCLC-I^high^ populations in knockdown samples. **i.** Kaplan-Meier survival curves stratified by *RORB* (left) and *RORA* (right) expression levels in human SCLC patients from the TU-SCLC cohort (Liu *et al.*). P-values were determined by log-rank test. **j.** Proliferation curves of control and *Rorb* knockdown murine KP11 SCLC cells. Fold change in cell number is plotted over the culture days (n = 3 technical replicates). Data represent mean ± s.e.m.; p-values were calculated by a 2-way ANOVA test. **k.** Tumour growth kinetics in immunodeficient NSG mice. KP11 cells with control or *Rorb* knockdown were implanted subcutaneously, and tumour volume was measured over time (n = 5-6 mice per group). Data represent mean ± s.e.m.; p-values were calculated by a 2-way ANOVA test. **l.** Tumour growth kinetics as in (k) in immunocompetent syngeneic mice (n = 7-8 mice per group). P-values were calculated by a 2-way ANOVA test. **m.** Tumour weight at endpoint as in (l). Data represent mean ± s.e.m.; individual data points shown; p-values were determined by using the Wilcox test to perform pairwise comparison of *Rorb* knockdown lines with s*hGfp* control. **n.** Dot plot showing significantly upregulated gene programs (y-axis) in *Rorb* knockdown versus control tumours from bulk RNA-seq of subcutaneous allografts grown in immunocompetent mice as in (l). Gene ratio (x-axis) indicates the fraction of genes in each pathway that are upregulated. Dot size represents gene count; colour intensity indicates statistical significance (−log10 adjusted p-value).

To examine the consequences of RORB loss-of-function, we treated *RPR2* tumour-derived organoids with AGN 193109, a pan-RAR antagonist with presumed activity against RORs ^56^. As RORA and RORB are the only family members expressed in NE SCLC cells in the *RPR2* model (**Supplementary Fig. 14b**), the effects of AGN 193109 likely reflect inhibition of these two TFs. Treatment with AGN 193109 for 72 h decreased NCAM1^+^ NE cells, increased ICAM1^+^ cells, and suppressed NE markers (**Supplementary Fig. 14b,c**). Next, we individually knocked down RORA and RORB using distinct shRNAs in the same *RPR2* tumour organoid system. RORB knockdown mirrored the phenotype of AGN treatment, including an increase in NCAM1^+^ ICAM1^+^ Double-positive (DP) cells (**Fig. 6d,e** and **Supplementary Fig. 14d,e**), together with strong downregulation of NE marker expression (**Supplementary Fig. 14f**). In contrast, RORA knockdown did not significantly impact cancer cell lineages or NE marker levels (**Supplementary Fig. 14g-j**). Knockdown of RORB in two established NE SCLC cell lines grown in 2D culture (KP1 and KP11) again promoted the emergence of NCAM1^+^ ICAM1^+^ DP cells and resulted in reduced protein levels of NE markers (**Supplementary Fig. 15a-f**), though the phenotype was less evident than in organoids, presumably reflecting the reduced plasticity of established cell lines.

We then performed single-cell RNA-seq on control (shRNA against GFP) and RORB knockdown (two shRNAs) samples from five tumour-derived organoid lines to determine how RORB loss affects cell states. UMAP-based visualisation and Leiden clustering identified eight cancer cell clusters with distinct enrichment of control and RORB knockdown cells (clusters 2, 3, 4, 6 for the controls, and clusters 1, 5, 7, 8 for RORB knockdown cells) (**Supplementary Fig. 15g**). Using the 6-month *RPR2* primary tumour scRNA-seq as reference, we mapped the six archetypes onto the organoid data. This analysis revealed a strong decrease in NE1 cells and a concomitant increase in Intermediate and nonNE1 populations in RORB knockdown samples compared to control cells (**Fig. 6f,g**). Differential abundance testing with the MILO package ^57^ confirmed that neighbourhoods containing Intermediate and nonNE cells were significantly increased upon RORB downregulation (**Supplementary Fig. 15h**). Gene programs associated with inflammation, interferon signalling, and antigen presentation were upregulated in clusters enriched for RORB knockdown cells (1,5,7,8), which also expressed NE genes at low levels (**Supplementary Fig. 15i**). At single-cell resolution, subpopulations of cancer cells exhibiting high activity for the SCLC-I signature uniquely emerged in RORB knockdown organoids, correlating with lower activity for an ASCL1 targets signature (**Fig. 6h**). These experiments validate the BoBa-T predictions and identify RORB as a regulator of the NE state in SCLC.

### Low RORB levels attenuate SCLC growth through immune-mediated mechanisms

In two independent human patient cohorts ^19, 58^, higher expression of *RORB*, but not *RORA*, was associated with worse overall survival (**Fig. 6i** and **Supplementary Fig. 16a**). These observations suggested that lowering RORB levels may impair SCLC tumour growth. However, *RORB* knockout in human SCLC cell lines does not inhibit the expansion of these cells in culture (**Supplementary Fig. 16b** from the DepMap dataset), suggesting that such tumour-inhibiting effects may depend on the tumour microenvironment. In support of this idea, knockout of *RORB* in human SCLC-A cell line NCI-H69 (**Supplementary Fig. 16c**) did not impact tumour growth in immunodeficient NSG mice, although knockout cells displayed reduced expression of NE marker genes *in vitro* (**Supplementary Fig. 16d,e**) and had a lower percentage of NCAM1^+^ cells in the xenograft tumours *in vivo* (**Supplementary Fig. 16f**). As loss of RORB promotes the transition of NE cells to the inflammation-associated Intermediate state, we focused subsequent analyses on cell models that can grow in immunocompetent conditions.

RORB knockdown mouse KP11 SCLC cells (derived from an *RPR2* mutant tumour) grew at rates comparable to controls in culture, consistent with the human data (**Fig. 6j**). RORB knockdown subcutaneous allografts also grew similarly to controls in NSG mice (**Fig. 6k**). In contrast, RORB knockdown tumours grew more slowly than control tumours in syngeneic immunocompetent hosts (**Fig. 6l,m**). As expected, these RORB knockdown tumours showed lower ASCL1 expression (**Supplementary Fig. 16g,h**). The growth of liver metastases upon tail vein injections in immunocompetent hosts was also inhibited by RORB knockdown (**Supplementary Fig. 16i,j**).

To understand this immune-mediated growth suppression, we performed bulk RNA sequencing on RORB-knockdown tumours grown in immunocompetent mice. This identified predominantly upregulated genes (**Supplementary Fig. 16k**), consistent with RORB functioning mostly as a transcriptional repressor in this context. The top upregulated genes included MHC-I and antigen presentation machinery, immune effector molecules, and markers of nonNE differentiation. Gene set enrichment analysis showed that leukocyte migration and cytokine secretion pathways were most significantly activated in RORB knockdown versus control tumours (**Fig. 6n**).

Thus, cancer cell-intrinsic plasticity changes driven by RORB loss, including enhanced antigen presentation and nonNE differentiation, remodel the microenvironment and render RORB-deficient SCLC tumours susceptible to immune-mediated control.

## DISCUSSION

Our analysis of mouse models and human tumours identified a transitory “Intermediate” state between the neuroendocrine and non-neuroendocrine lineages, in both *de novo* SCLC and the transformation from lung adenocarcinoma to SCLC. This state provides mechanistic insight into cancer cell plasticity and intratumoural heterogeneity, and points to new therapeutic strategies in one of the most aggressive human cancers.

Intermediate cancer cells maintain open chromatin at regulatory elements associated with both NE and nonNE lineages, even though the transcription factors whose motifs define these regions are expressed at low levels. Our data support a multi-layered model for how this permissive chromatin architecture is established and sustained. Low residual levels of lineage-specific TFs such as ASCL1 may help preserve accessibility at select NE regulatory elements. In parallel, non-sequence-specific chromatin remodellers such as BRD4 and MLL, acting in concert with broadly active, inflammation-associated TFs such as AP-1 and NFκB, may further sustain state-specific open chromatin in the absence of dominant lineage drivers. The cell-cycle activity characteristic of Intermediate cells may additionally contribute to epigenetic reprogramming, analogous to how loss of p53 can facilitate reprogramming towards a pluripotent stem cell state^59^. As Intermediate cells are present in *ex vivo* tumour organoids devoid of stromal cells, the mechanisms that establish and maintain this state are likely predominantly cancer cell intrinsic. These observations do not exclude a role for the tumour microenvironment, as shown in pancreatic cancer ^60, 61^, and cancer cell-stroma interplay in SCLC plasticity remains an important question.

Plastic transitional states have been identified across multiple cancer types, including prostate cancer ^35^, pancreatic cancer ^61, 62^, lung adenocarcinoma ^63, 64^, squamous cell lung cancer ^65^, colorectal cancer ^49^, and melanoma ^66, 67^. Although their molecular features differ, a recurrent theme is the association between inflammation and plasticity ^35, 61^, as observed in Intermediate SCLC cells, suggesting that inflammatory signalling may represent a broadly conserved mechanism for chromatin opening and lineage destabilisation. Our finding that the Intermediate signature peaks at the minimal residual disease stage during LUAD-to-SCLC histological transformation suggests that this state is not simply a feature of established SCLC heterogeneity, but a more general bottleneck through which cells pass during neuroendocrine lineage commitment.

BoBa-T identified RORB as a regulator of the NE state in SCLC, which we validated in loss- and gain-of-function studies: RORB knockdown reduced NE gene expression, while overexpression promoted an NE state. It remains to be determined if RORB has similar roles in other neuroendocrine cancer types. RORB is best known as a key component of the machinery that controls circadian rhythms ^68, 69^. In the circadian oscillator, RORB is thought to act as a transcriptional activator ^70^, which is distinct from the dual activator/repressor activity predicted by BoBa-T in SCLC (activator of the NE state, repressor of the nonNE state). Activating RORB may promote an NE state more amenable to immunotherapies targeting DLL3, an ASCL1 target; conversely, RORB inactivation may reprogram SCLC cells towards a low/nonNE state amenable to immune checkpoint blockade ^71, 72^. Agonists and antagonists of RARs and RORα/γ already exist ^73–75^, and some of these molecules may have similar activity against RORB or serve as scaffolds for RORB-specific compounds. Targeting molecules such as RORB may also link cancer cell plasticity to circadian rhythms, including how they influence immune cell activity and immunotherapy efficacy, given recent evidence that time-of-day scheduling of immunotherapy improves outcomes in lung cancer patients ^76^.

Collectively, our study reveals that SCLC tumours harbour a previously uncharacterised Intermediate state defined by permissive chromatin, bipotent differentiation capacity, and intrinsic immunogenicity. Rather than viewing SCLC heterogeneity as a fixed distribution of terminal NE and nonNE fates, our data support a dynamic model in which cancer cells traverse this plastic Intermediate state during lineage switching, with the transition gated by transcriptional regulators such as RORB. These findings reframe SCLC plasticity as a regulated, and therefore therapeutically targetable, biological process.

## METHODS

### Ethics statement

Mice were cared for in accordance with practices approved by the NIH, the Stanford Institutional Animal Care and Use Committee (IACUC), and the Association for Assessment and Accreditation of Laboratory Animal Care (AAALAC). The study protocol received approval from the Stanford Administrative Panel on Laboratory Animal Care (APLAC) (protocol 13565).

### Animal studies

For tumour induction in the autochthonous *RPR2* model (*Rb^f/f^p53^f/f^Rbl2^f/f^ crossed to Rosa26^LSL-Luc^* or *Rosa26^LSLmTmG^*), 9-12-week-old male and female mice were infected by intratracheal instillation with 2 × 10^10^ PFU/mL of Adeno-CMV-Cre (Baylor College of Medicine). Mice were euthanised at ∼24 weeks post-infection, or at the first sign of respiratory distress, in accordance with Stanford APLAC guidelines.

SCLC cell lines were derived from *Rb^f/f^p53^f/f^* (KP1) or *Rb^f/f^p53^f/f^Rbl2^f/f^* (KP11) mutant tumours, with associated Luciferase (luc) or the membrane-targeted tandem dimer Tomato-membrane targeted GFP (mTmG) reporters, as previously described ^77, 78^. NCI-H69 cells were obtained from ATCC (HTB-119). All cell lines tested negative for mycoplasma (Lonza), and their identity was verified by Short Tandem Repeat (STR) genotyping. For subcutaneous allograft experiments, B6129SF1/J mice (Jackson Laboratories, stock no. 101043) were used as immunocompetent recipients, and Nod.Cg-Prkdc^scid^IL2rg^tm1WjI/SzJ^ (NSG) mice (Jackson Laboratories, stock no. 005557) were used as immunodeficient recipients. Mice were engrafted with 0.1-0.25 × 10^6^ cancer cells in 100 µL of antibiotic-free, serum-free RPMI medium, mixed 1:1 with Matrigel (BD Matrigel, 356237), at 8-12 weeks of age. An equal number of male and female mice was used in all experiments. Tumours were measured every other day starting on Day 7 post-injection, and volumes were calculated as 0.5 × length × width^2^. Mice were euthanised when the largest tumour within the experimental cohort reached 1,500 mm^3^, in accordance with Stanford APLAC guidelines. All mice were housed at 22 °C, 40% humidity, and a 12 h light/dark cycle.

### Cell lines

All SCLC cells (KP1, KP11, NCI-H69, as above) were cultured in RPMI-1640 supplemented with 10% Bovine Growth Serum (Hyclone), 1× GlutaMax (Invitrogen), and 1× PenStrep. Cell lines were grown in suspension and dissociated into single cells by brief trituration in 1:4 diluted 0.25% Trypsin, followed by immediate quenching with serum-containing RPMI. Cell lines were cultured in humidified incubators at 37°C with 5% CO2.

### Single-cell dissociation of murine lung tumours

Large tumours from *RPR2* mice were micro-dissected and finely chopped with a razor blade. Pooled tumour pieces from individual mice were digested in 25 mL of DMEM-F12 media (11320033, Gibco) containing 1 mL of 5E mix (five-enzyme mix: DNase I (10104159001, Roche), Elastase (LS002279, Worthington Biochemical Company), Collagenase Type I (C0130, Millipore Sigma), Collagenase Type II (C6885, Millipore Sigma), Collagenase Type IV (C5138, Millipore Sigma)) for 45 min on a shaker at 37°C. The digested mixture was then passed through a 70 µm filter, spun down at 500 g for 5 min. The supernatant was discarded, and the pellet was resuspended in 5 mL of red blood cell lysis buffer (00-4333-57, eBioscience™) for 1-2 min at room temperature. RBC lysis was stopped by adding 35 mL of DPBS (Dulbecco’s Phosphate-Buffered Saline), after which cells were centrifuged and washed again with 30 mL of DPBS. The resulting single cells were then processed according to their intended downstream analysis.

### Generating tumour-derived organoid lines

Isolated cancer cells from tumours were resuspended in 1 mL DPBS, and live cells were counted using a Countess™ automated cell counter. If large cell clusters or debris were observed, the cell suspension was passed through a 40 µm filter, and the flow-through was spun down at 500 g for 5 min in a swing-bucket rotor to ensure the cell pellet settled at the bottom of the tube. The supernatant was discarded, and ∼2 × 10^6^ cells were gently resuspended in 100 µL of thawed Matrigel, and 8 µL Matrigel domes were dispensed onto culture plates, with 10 domes per well of a 6-well plate. The plate was incubated upside down at 37°C for 15-30 min until the domes solidified, after which 2 mL of appropriate organoid medium was added to each well. Organoid media consisted of Advanced DMEM-F12 (12634010, Gibco), 1× BEGM bronchial epithelial growth bullet kit (CC-3170, Lonza), 1× Glutamax (35050061, Gibco), and 1× PenStrep (15070063, Gibco). For the first 24 h of plating, the organoid media were supplemented with 10 µM ROCKi Thiazovivin (S1459, Selleckchem). To eliminate RB and p53-proficient non-cancer cells within the organoids, the lines were treated with 10 µM Palbociclib (PD-0332991, Selleckchem) and 3 µM Nutlin-3 (S1061, Selleckchem) for the first two weeks of culture. Successful selection was confirmed by the absence of TdTomato^+^ non-cancer cells in the organoid lines after 2 weeks. The organoid medium was changed every 2-3 days, and tumour organoids were split and expanded upon confluence using TrypLE Express (12605010, Gibco).

### Knockdown and overexpression studies

Short hairpin RNAs targeting eGFP (5’-TACAACAGCCACAACGTCTAT-3’), *Rorb* (#1: 5’-CCGGCGGGATAACAATGTCTGAGATCTCGAGATCTCAGACATTGTTATCCCG-3’; #2: 5’-CCGGCCATTGTACAAGGAGCTCTTTCTCGAGAAAGAGCTCCTTGTACAATGG-3’), and *Rora* (#1: 5’-CCGGCCGAAGACGAAATCGCGTTATCTCGAGATAACGCGATTTCGTCTTCGG-3’; #2: 5’-CCGGCACACACATCTCAAATTGAAACTCGAGTTTCAATTTGAGATGTGTGTG-3’) from the MISSION shRNA library (Sigma-Aldrich) were packaged in 293T cells via co-transfection with delta8.2 and VSV-G using Lipofectamine 3000 (L300015, Thermo). Lentiviral supernatant was collected 48 h post-transfection, filtered, and either directly added to suspension cultures of murine SCLC lines (KP1 and KP11) supplemented with 8 µg/mL polybrene (TR1003G, Sigma) or concentrated with a 1× PEG solution. For lentiviral transduction of organoids, established organoid lines were dissociated into single cells with TrypLE Express and Accutase (07922, Stem Cell Technologies), and 10^6^ cells were resuspended in 500 µL of organoid media with 8 µg/mL polybrene and 10 µM ROCKi in one well of a 24-well plate. An appropriate volume of concentrated virus, corresponding to the control or targeting shRNAs, was added to each well, and the plate was centrifuged at 300 g for 30 min at room temperature. The cells were then allowed to recover at 37°C for 3-4 h. Afterwards, they were spun down, resuspended in Matrigel, and plated as described. 24 h after transduction, organoids and cell lines were selected with 1 µg/mL puromycin (A111380, Thermo Fisher Scientific) for 48 h. Cells were harvested and processed for downstream analysis three days after selection.

The pCDH-3Flag-GFP-puro (167463, Addgene) and pLV-Puro-EF1A-mRorb (VB900095-9998atv, VectorBuilder) were packaged into lentiviral particles as described above, and organoid lines were transduced with concentrated virus via spinfection.

### Immunoassays and protein extraction

Cells were lysed in RIPA buffer (89900, Thermo Fisher Scientific) supplemented with protease and phosphatase inhibitors (05892970001, Roche cOmplete ULTRA tablets, Mini, EASYpack and 04906845001, PhosSTOP EASYpack). Protein concentration was measured using a Pierce™ BCA protein assay kit (23227, Thermo Fisher Scientific). Immunoassays were performed using the capillary-based Simple Western™ assay on the Wes™ system (ProteinSimple, 004-600) following the manufacturer’s protocol. Compass software v4.0.0 was used to analyse and visualise the results. The following antibodies were used: HSP90 (4877S, Cell Signaling, 1:5000), RORB (17635-1-AP, Proteintech, 1:100), UCHL1 (HPA005993, Sigma, 1:5000), ASCL1 (sc-374104, Santa Cruz, 1:200), INSM1 (A-8, sc-271408, Santa Cruz, 1:100) and Chromogranin A (PA5-35051).

### Quantitative Reverse transcription PCR

Cells from *in-vitro* experiments were counted (10-50,000 cells), washed once in PBS, and lysed in the lysis buffer from SYBR™ Green Fast Advanced Cells-to-CT™ Kit (A35380, Thermo Fisher). RNA extraction and cDNA preparation were performed following the recommended protocol. RT-PCR was conducted using the Maxima SYBR Green/ROX qPCR Master Mix (K0221, Thermo Fisher) on a Bio-Rad CFX384 machine. CT values for all genes were normalised to the reference gene *Gapdh* and presented as fold changes relative to the control sample (ddCT).

### Bulk RNA sequencing

Control and knockdown subcutaneous tumours were dissected, chopped with a razor, and incubated in 10 mL of DMEM-F12 medium containing 250 µL of 5E enzyme mix for 15 min at 37°C. The cancer cells were filtered and washed in DPBS, and red blood cell lysis was performed as described above. 1-2 × 10^6^ cells were used for RNA extraction using the AllPrep DNA/RNA kit (80204, Qiagen). Library preparation and sequencing were done by Novogene™. RNA counts were aligned to the mm10 genome using Hisat2 ^79^, and FeatureCounts ^80^ was used to generate raw counts. Differential RNA-seq analysis was conducted using DESeq2 ^81^.

### Flow Cytometry analysis and cell sorting for cell state analysis

For flow cytometry analysis of neuroendocrine and non-neuroendocrine cancer cell populations, single-cell suspensions from either *in vivo* lung tumours or *in vitro* organoid or cell lines were washed once in Annexin V binding buffer (BMS500BB, eBioscience™ Binding Buffer for Annexin V, Thermo Fisher Scientific) and resuspended in an antibody cocktail prepared in the same buffer. The antibodies used were as follows: mouse NCAM1/CD56-APC (FAB7820A, R&D Systems, 1:200), mouse ICAM1/CD54-BV785 (116145, BioLegend), and mouse H-2K^d^/H-2D^d^-PE (114708, BioLegend). Cells were stained for 30-45 min at 4°C, then washed in 1 mL of Annexin V binding buffer and resuspended in buffer containing a 1:10,000 dilution of DAPI (1 mg/mL). Flow cytometry was performed with a 100 µm nozzle, and gates were set based on FMO controls. For lineage analysis experiments, 5,000 cells each of NCAM1^+^, ICAM1^+^, and double-positive populations were sorted into 100 µL of organoid medium supplemented with 10 µM ROCKi. After sorting, cells were centrifuged at 300 g for 10 min, the supernatant was carefully removed, leaving approximately 10 µL, and cells were resuspended in a total of 80 µL of Matrigel. The cells were then plated as described above and allowed to grow for 3 weeks without additional passage before reanalysis.

### MIBI-TOF tissue staining

FFPE sections from 3 different *RPR2;mTmG* tumour-bearing lungs were baked at 70°C for 20 minutes and deparaffinised through a graded xylene and ethanol series (xylene ×3, 100%, 95%, 80%, 70% ethanol, ddH₂O) using a Leica ST4020 Linear Stainer (180 sec processing time, 3 dips per station). Heat-induced epitope retrieval (HIER) was performed using 1× Dako Target Retrieval Solution pH 9 (Agilent, S2375) in a Thermo Scientific PT Module (97°C for 10 min, cooled to 65°C). Slides were washed in TBS-Tween wash buffer containing 0.1% BSA and blocked for 1 hour at room temperature in blocking buffer consisting of 1× TBS-Tween, 5% normal donkey serum, 0.1% Triton X-100, and 0.05% sodium azide. Metal-conjugated primary antibodies were applied in antibody buffer (1× TBS-Tween, 5% normal donkey serum) at 100 µL per section and incubated overnight at 4°C in a humidified chamber. Slides were washed twice in wash buffer and post-fixed for 5 minutes in 2% glutaraldehyde / 4% PFA prepared in low-barium PBS. Slides were then dehydrated through a graded ethanol series via the Linear Stainer and dried under vacuum for at least 1 hour prior to imaging. All reagents were prepared in metal-free, low-barium PBS using plastic-only labware to minimise metal contamination. Images were acquired on a MIBIscope at super-fine preset (2048 × 2048 pixels, 800 µm field of view). MIBI image analysis is described below.

### Cancer cell sorting for single-cell multi-omics

Freshly isolated cancer cells from lung tumours were resuspended in PBS, counted, and divided into samples of approximately 4 × 10^6^ cells in preparation for flow cytometry staining. For staining, cells were resuspended in 200 µL of an antibody staining cocktail prepared in Cell Staining Buffer (420201, BioLegend), containing a mouse Fc receptor blocker (anti-mouse CD16/32 antibody, 101302, BioLegend), and stained for 45 min at 4°C. The following antibodies were used: CD45-Pacific Blue (BioLegend 103126, clone 30-F11, 1:100), CD31-Pacific Blue (BioLegend 102422, clone 390, 1:100), TER-119-Pacific Blue (BioLegend 116232, clone TER-119, 1:100), CD24-APC (eBioscience 17-0242-82, clone M1/69, 1:200). After staining, the cells were washed in 1 mL of cell staining buffer and then resuspended in buffer containing 1:10,000 dilution of 10 mg/mL DAPI solution. Flow cytometry was performed with a 100 µm nozzle on a BD FACSAria II using the FACSDiva software. Single cells were gated based on forward and side scatter values, and cancer cells were gated as CD45^neg^ Ter119^neg^ CD31^neg^ DAPI^neg^ CD24a^pos^ (see Supplementary Fig. 1a). ∼10^6^ cancer cells were sorted into 100 µL of PBS and immediately processed for downstream analysis.

### Nuclei isolation and single cell multi-omics library preparation

Cancer cells from pooled tumours of 2 female and 2 male mice were sorted separately as described above and processed for single-cell multi-omics using the 10x Genomics platform. Nuclei isolation was performed according to the kit’s protocol (CG000365) with the additional optimisation of the cell lysis time, which was standardised to 3 min on ice. Successful nuclear isolation was confirmed by trypan blue staining, and nuclei were immediately loaded onto the 10x Chromium X controller, targeting 10,000 cells per sample. Samples from male and female mice were processed separately. Library preparation was performed following the manufacturer’s protocols using the Chromium Next GEM Single Cell Multiome ATAC + Gene Expression Reagent Kits (10x Genomics; PN-1000280). Completed libraries were sequenced on an Illumina NovaSeq 6000 to target more than 20,000 reads per cell with the 10x Genomics-recommended paired-end sequencing mode for dual-indexed samples.

### Single-cell RNA-seq of cancer cells from tumors 4-and-5-month post initiation

GFP+ cancer cells from pooled tumors of *RPR2;mTmG* mice (2 mice/timepoint) were sorted separately and processed for scRNA-seq. Approximately 100k sorted cells/sample were fixed using the Evercode™ Low input fixation kit v4(Parse Biosciences, LCF500), according to kit protocol. These samples were barcoded and processed for sequencing using Evercode Mega v4 kit (ECWT4500). Completed libraries were on Illumina NovaSeq X Plus to target ∼20K reads/cell with paired end sequencing. Sequencing data were demultiplexed, aligned to the mm10 genome, and count matrices were generated Parse Biosciences™ Trailmaker pipeline.

### Single-cell RNA-seq of control and Rorb knockdown organoids

To evaluate cell state changes upon *Rorb loss*, five independently derived tumour organoid lines were transduced with either control (*shGfp*) or *Rorb*-targeting shRNAs (*shRorb#1, shRorb#2*) as described above. 3-4 days after puromycin selection, knockdown efficiency was evaluated by RT-qPCR for *Rorb,* and the cancer cell state-alteration phenotype was confirmed by flow cytometry and RT-qPCR for neuroendocrine genes. Approximately 10^6^ control and knockdown cells from each organoid line were collected within 7 days of selection and fixed according to the protocol detailed for the Evercode™ Fixation kits. After all samples were collected and fixed, single-cell library preparation was performed using the Evercode™ WT V3 kit (100K cells) according to the manufacturer’s protocol. Completed libraries were sequenced on an Illumina NovaSeq X Plus to target more than 50,000 reads per cell with recommended paired-end sequencing. Sequencing data were demultiplexed, aligned to the mm10 genome, and count matrices were generated using a custom pipeline provided by Parse Biosciences™.

### Single-cell multi-omics: Quality control and normalisation

For pre-processing and initial quality control of the single-cell multi-omics data, raw gene expression and chromatin accessibility matrices from aggregated male and female mice samples (M1 and M2) (Cell Ranger arc v2.0 pipeline, mm10 reference) were imported into R and analysed using Seurat and Signac ^82^. Gene expression counts were used to create a Seurat object ^83^ (RNA assay), and a corresponding ATAC assay was generated from peak counts and the 10x fragment file. Gene annotations were obtained from EnsDb.Mmusculus.v79, with chromosome names converted to UCSC style and the genome set to mm10 to ensure consistency with the multi-omics reference.

For ATAC-seq quality control, standard chromatin metrics were computed on a per-cell basis, including nucleosome signal and transcription start site (TSS) enrichment. Cells were retained if they passed joint RNA and ATAC thresholds (1,000–100,000 ATAC fragments and 750–25,000 RNA UMIs) and exhibited a nucleosome signal < 4, thereby removing low-complexity nuclei while preserving high-quality multi-omics profiles. Heterotypic doublets identified by DoubletFinder ^84^ were also removed from the dataset at this stage. Peaks were recalled directly from the aggregated ATAC fragments using MACS2 ^85^ via Signac, and a new chromatin assay (“peaks”) was constructed by quantifying fragment counts per called peak. This peak-by-cell matrix was then normalised using TF–IDF, and latent semantic indexing was applied to obtain a low-dimensional representation of chromatin accessibility. To correct for batch effects across samples, Harmony integration ^86^ was separately applied to the ATAC and RNA modalities. For the ATAC data, Harmony was run on the LSI components using sample identity as the covariate, and the resulting Harmony embedding (harmony_1) was used to compute a UMAP representation of chromatin state. For the RNA data, the percentage of mitochondrial reads/cell was first calculated, and normalisation was performed with SCTransformv2 ^87^, regressing out mitochondrial content, sample identity, and cell-cycle scores (S and G2/M). Principal component analysis was then performed on the SCTransform-normalised data, followed by Harmony integration across samples, and UMAP visualisation in the Harmony-corrected RNA space (harmony_2). Gene activity scores were calculated for all protein-coding genes using the GeneActivity() function of Signac with default parameters, and the resulting matrix was stored in the “gene_activity” assay of the Seurat object.

Finally, a joint multimodal representation was derived using a weighted nearest-neighbour (WNN) framework. Harmony-corrected RNA and ATAC embeddings were used as inputs to construct a multimodal neighbour graph, from which a joint UMAP embedding was generated. Graph-based clustering (Leiden, resolution 0.5) was performed on the WNN graph, which determined 15 cell clusters. To identify potential contamination of non-cancer cells in the dataset, the scMCA package ^88^ was used to map each cell’s identity to the publicly available mouse cell atlas (MCA v.2.0.0). Cell clusters that mapped with high confidence to normal lung cells were removed. The validity of this approach was further confirmed by the observations that these clusters were far away from the cancer cell clusters in the WNN UMAP space and did not express any cancer cell markers. Since low-quality cells usually cluster together, extra rounds of quality control and data filtering were performed on the dataset by assessing nCounts and mitochondrial percentage per cluster. Additionally, FindMarkers() was run on all clusters, and those whose top markers included multiple mitochondrial genes were further evaluated and removed. Following this stringent filtering pipeline, 9,000 cancer cells were retained and used for all downstream analyses.

### Archetype analysis

Archetype analysis was run on the combined (M1 and M2) scRNA-seq dataset using the principal convex hull analysis (PCHA) function from the Python package py_pcha (https://github.com/ulfaslak/py_pcha). The top 8 components of the Harmony dimensions were used. PCHA was run with 3 to 8 archetypes, and the number of archetypes was chosen using the knee of the explained variance vs number of archetypes plot. Six archetypes explained 95% of the variance from the Harmony components and were chosen for further analysis. Archetype scores are output from the PCHA function as the S matrix (archetypes × samples). Archetypal identity of individual cells was determined from these scores using a cutoff of S > 0.5. Marker genes for each archetype were identified using the FindAllMarkers() function in Seurat on the normalised SCT counts, with the Wilcoxon test (logFC threshold = 0.25, p_val_adj < 0.01, and min.pct = 0.25), which identified 3,550 differential genes. Pathway analysis across Gene Ontology, KEGG, REACTOME, and WIKIPATHWAYS was performed for all marker genes of each archetype using gProfiler ^89^ (Benjamini-Hochberg Correction, FDR < 0.05). The top 30 markers of each archetype were used as its “signature” and applied to other datasets (described below) to identify equivalent cancer populations. Differential chromatin regions were identified using FindAllMarkers() on the “peaks” assay using the Likelihood Ratio (LR) test (logFC threshold = 1, p_val_adj < 0.05, and min.pct = 0.1), which identified a total of 9,569 peaks.

### Transcriptomic gene signature scoring

Hallmark pathway and Gene Ontology biological process gene sets—including protein secretion (HALLMARK_PROTEIN_SECRETION), epithelial–mesenchymal transition (HALLMARK_EPITHELIAL_MESENCHYMAL_TRANSITION), inflammatory response (HALLMARK_INFLAMMATORY_RESPONSE), and antigen processing (GOBP_ANTIGEN_PROCESSING_AND_PRESENTATION_OF_ENDOGENOUS_ANTIGEN), were obtained from MSigDB ^90^ via the MSIGDBR package for mouse (H and C5 collections).

Published ASCL1 ^23^ and NFIB ^30^ gene targets were identified through combined ChIP-seq and RNA-seq analysis of these factors in *RPR2* lung tumours. NE and nonNE signatures were created from previously published bulk RNA-seq data^15^ of sorted HES1-positive (non-neuroendocrine cells) and HES1-negative (neuroendocrine) cancer cells from *RPR2* lung tumours. The genes that were significantly upregulated in HES1-positive cells, with logFC > 1 and p.adj < 0.001, formed the nonNE signature, while those significantly downregulated in these cells, with logFC less than −1 and p.adj < 0.001, formed the NE signature.

Human SCLC transcriptional subtype signatures were obtained from three different bulk SCLC patient studies: Nabet *et al.* ^37^, Gay *et al.* ^8^, and Liu *et al.* ^19^. Subtype-enriched gene sets corresponding to SCLC-NE-I (NMF3, Nabet *et al.*), SCLC-nonNE-I (NMF4, Nabet *et al.*), SCLC-A, SCLC-N, and SCLC-I-like states were obtained from supplemental differential expression tables, filtered on log-fold change and significance, and assigned to the subtype with the highest mean expression where multi-subtype data were available. These human subtype signatures were then converted to mouse orthologs to enable scoring in the mouse multi-omics dataset.

All curated gene sets, including Hallmark and GO pathways, inflamed SCLC programs, transcription factor targets, and SCLC subtype signatures, were applied to the SCTransform-normalised data using Seurat’s AddModuleScore() function. For each gene set, the corresponding mouse gene list, a gene set–specific number of control genes, and a fixed number of expression bins were provided, enabling the generation of normalised per-cell signature scores on the SCT assay and their use for downstream analyses.

### Gene Set Enrichment Analysis on RPR2 archetype

Identity signatures were derived from the 6-month dataset for Intermediate cells and for the pooled NE (Neuroendocrine1 and Neuroendocrine2) and nonNE (nonNE1 and nonNE2) groups, using identical settings throughout so that the three are directly comparable. For each group, one-versus-rest differential expression against all remaining archetypes was computed on the SCT assay with a Wilcoxon rank sum test (min.pct = 0.1, no log fold change threshold), and genes were ranked by average log2 fold change. Ranked lists were tested by GSEA against GO biological process terms (gseGO, org.Mm.eg.db, Benjamini-Hochberg correction, followed by semantic similarity simplification at a cutoff of 0.7) and against mouse Reactome gene sets from MSigDB, with terms retained at FDR below 0.05. For display, one representative pathway per distinct biological theme was selected manually from the full significant list, and plotted as normalised enrichment scores, such that positive values indicate enrichment in the group of interest and negative values enrichment in the remaining archetypes.

### CellRank analysis of the 6-month RPR2 cancer cells

To assess whether the Intermediate archetype behaves as a stable state within a single timepoint, we fitted CellRank’s GPCCA estimator to the 6-month dataset alone. A kNN graph was computed in scanpy on the harmony_2 integrated embedding, using the embedding that had not been cell-cycle regressed, since cycling is a feature of the Intermediate state rather than a confounder. This graph was combined with the Palantir pseudotime computed for the object to construct a CellRank PseudotimeKernel, and GPCCA was fitted to the resulting transition matrix with n_states = 10, at which each of the six archetypes forms its own macrostate and the dominant Generalist population resolves into four near-duplicate sub-macrostates. Terminal and initial states were predicted and fate probabilities computed using CellRank defaults, and fate probabilities were projected onto the six non-redundant vertices so that the Generalist sub-macrostates did not dominate the layout. GPCCA.compute_lineage_drivers function was used to determine the driver genes varying along the NE to nonNE pseudotime axis.

### RPR2 tumour evolution analysis with temporal scRNA-seq datasets

#### Preprocessing of the *RPR2 4-*month and 5-month cancer cell scRNA-seq

scRNA-seq dataset of *RPR2* cancer cells from tumours 4 and 5 months post initiation were processed as two samples per timepoint. Cells were retained at 500 to 6,000 detected genes, fewer than 20,000 UMIs and less than 20% mitochondrial reads, and counts were normalised with SCTransform (method = “glmGamPoi”, regressing percent mitochondrial reads). Archetype identity was assigned by anchor-based label transfer from the 6-month multiome dataset, in which Generalist cells were excluded and the remaining archetypes downsampled to 300 cells each to balance the reference. Anchors were identified with FindTransferAnchors against the reference harmony_2 reduction (dims = 1:30, k.anchor = 15, nn.method = “rann”, n.trees = 200) and labels transferred with TransferData over the same dimensions, using the full 4- and 5-month object prior to any downsampling.

#### Integration of the 4-, 5- and 6-month *RPR2* cancer cell datasets

The Specialist archetypes of the 6-month dataset were merged with the label-transferred 4- and 5-month cells, downsampled to 2,000 cells per sample, to ensure balance in cancer cell numbers between the 3 timepoints. The RNA assay was split by timepoint, SCT transformation was again performed on the combined object followed by PCA reduction, and the timepoints were integrated with CCAIntegration on the SCT assay. The resulting integrated reduction was used as the joint embedding for all downstream analyses, and a UMAP was computed on the same space.

#### Optimal transport analysis on combined timepoint dataset with moscot

A moscot TemporalProblem was prepared on the integrated reduction with a sequential policy over timepoints encoded as 4, 5 and 6 at equal spacing, and solved with uniform marginals (no proliferation prior) under default entropic regularisation with mean cost scaling (max_iterations = 100,000). The integrated reduction was used rather than a new PCA so that the Parse-versus-multiome difference was not read as temporal signal. Couplings were wrapped in a CellRank RealTimeKernel with diagonal self-transitions at the terminal timepoint and no connectivity blending, then aggregated by transferred archetype label to give the per-state transition rates reported throughout.

The analysis was repeated on SoupX ambient-corrected 4- and 5-month counts with the cell set fixed (Pearson r = 0.994 and 0.998) and on a larger object with the 4- and 5-month cap raised from 2,000 to 8,000 cells (r = 0.997 and 0.978), in both cases giving concordant per-state rates.

#### Fate-biased differential expression and pathway enrichment

From each coupling, cells at the earlier timepoint were assigned the probability of reaching one of two destination groups at the next timepoint, renormalised over those groups. Top and bottom terciles of this fate bias were compared, the middle tercile discarded, and genes detected in fewer than 20 cells excluded. Differential expression used a Wilcoxon rank sum test, and genes ranked by log fold change were tested by GSEA against GO biological process terms (gseGO, org.Mm.eg.db, Benjamini-Hochberg, simplification at 0.7) and mouse Reactome sets (MSigDB) at FDR < 0.05. The comparisons were 4-month Intermediate fated for NE versus nonNE, 5-month Intermediate fated for nonNE versus retained, and 5-month NE fated for Intermediate versus retained.

#### CellTag barcoding and analysis

Organoid lines F4 and M1 were transduced with the CellTag V1 lentiviral barcode library (Addgene #115643, 19,972 barcodes) and sampled at two timepoints one month apart. Libraries were prepared with Parse Biosciences Evercode WT chemistry across 16 sublibraries and processed with split-pipe (v1.8.2) against GRCm38 with a GFP-CellTag contig appended, so that CellTag transcripts carried the same cell barcode and UMI tags as the transcriptome. CellTag reads were identified by matching the V1 motif GGT[ACTG]_8_GAATTC within the sequence of reads aligned to that contig only, and the eight-nucleotide barcode was taken from between the flanks together with its cell barcode and UMI. Cell barcodes were disambiguated by sublibrary before pooling.

Per-cell barcode matrices were assembled with CellTagR and sequencing errors corrected with starcode (-s --print-clusters, default distance) applied to the barcode concatenated with the cell identifier, so collapsing acted within rather than across cells. The Addgene whitelist was applied after collapsing, leaving error-containing reads available to correct their parent barcode.

Barcode counts were binarised at one UMI and cells with 2 to 20 distinct barcodes retained, excluding single-barcode signatures and loads implausible for one founder. Clones were called from the Jaccard similarity of binarised profiles (CellTagR CloneCalling, cutoff 0.8), with the two timepoints of each line called jointly in a single barcode and clone-ID space. Joint calling returned 237 clones over 1,790 cells in F4 (64 shared) and 454 clones over 2,312 cells in M1 (204 shared).

#### Construction of the clone-bearing objects

Objects were built from the split-pipe DGE_filtered output, the same barcode set used for the CellTag matrices, so every clone-bearing cell was retained, then filtered on nFeature_RNA > 500 and percent.mt < 20. Depth ceilings were not applied: clone-bearing cells are two to three times deeper than untagged cells, so a fixed ceiling is confounded with clone detectability, removing 655 and 619 of F4’s 1,790 clone cells against 7 for the feature floor. Doublets were instead tested directly with scDblFinder within each first-round barcode well at the Parse-appropriate rate (3% per 10,000 cells) and removed. After QC, 232 clones (57 shared) remained in F4 and 441 (172 shared) in M1.

#### Cell state assignment and integration

Objects were normalised with SCTransform (percent.mt regressed out), then PCA (30 components), UMAP and Louvain clustering (resolution 0.5). Cell states were assigned by label transfer from the 6-month multiome reference, subset to the six archetypes and downsampled to 300 cells per state, with FindTransferAnchors against the reference Harmony space (dims 1:30, k.filter = 20, RANN, n.trees = 100; k.anchor = 25 for F4 and 15 for M1, each line’s swept optimum) and TransferData. Timepoints were then integrated by CCA (IntegrateLayers, SCT normalisation), with UMAP and clustering recomputed on the integrated reduction (dims 1:30, resolution 0.5), so integration affected the embedding and clustering only, not the state labels.

#### Optimal transport and the transition matrix

Cell-state flux between the two timepoints was estimated with Moscot TemporalProblem, prepared on the integrated reduction as the joint attribute with policy = “sequential”, and solved with uniform marginals, no proliferation reweighting and max_iterations = 100000, min_iterations = 100. The solution was passed to CellRank (RealTimeKernel.from_moscot, compute_transition_matrix (self_transitions = “diagonal”)) and state-level flows were computed with FlowPlotter.compute_flow. This is the same recipe, embedding and marginals used for the *in vivo* timepoint dataset, so that organoid and *in vivo* flux are comparable. CellTag barcodes were withheld from the transport problem entirely, a plain TemporalProblem being used rather than a LineageProblem. The clonal evidence and the transport solution are therefore statistically independent, so that the clones could be used to orient an edge and the transport size it. Every flux value reported is the percentage of the source state’s cells at the first timepoint that moved along that edge, equivalent to the flux quantity reported for the *in vivo* data.

#### Clonal attribution of transitions

A clone block is one clone at one timepoint; blocks of at least two cells were used. An edge from state x to y was credited only where y was absent from the parent block and present in the daughter block, so that only an appearance is scored and a pair of states held at both timepoints generates no directed credit. Within a block, daughter states were ordered so that no lineage could credit both x to y to z and x to z to y, contributions were scaled so that a clone could not count twice for the same pair, and candidate states were iterated in fixed archetype order for determinism.

For blocks whose parent set is not directly observed, the parent distribution was taken by composition matching, using as reference class those cross-timepoint clones whose second-timepoint state set is exactly the observed set, weighted by frequency. References are always second-timepoint blocks of cross-timepoint clones, and those never use the lookup, so no block is its own reference. Where no reference class existed the block fell back to the transport solution, which applied to 29 of 172 blocks in F4 and 6 of 195 in M1. A block of a cross-timepoint clone holding a single state was admitted, since its parent set is the clone’s own observed first-timepoint set and this is the least ambiguous case in the data. Where a parent produced two states, a stepwise rather than parallel origin was inferred when the conditional enrichment of the second state given the first exceeded 1.5 relative to the reference class, with a background comparison used only as a tie-break. Where clone and transport estimates were blended, the clone reference was weighted n/(n + 2). Clonal data was used to establish which edges exist and which way they point, and the transport supplied magnitude. Self-renewal is reported from the transport alone, since a lineage must be sampled at both timepoints to be counted and that selects for lineages that grew and diversified.

#### Clonal composition, patterns and pattern enrichment

Within-clone state co-occurrence was computed per timepoint as counts, Jaccard index and conditional composition. Clonal patterns were defined as the exact set of archetypes present in a clone at one timepoint, and all clones were included in the statistics. Patterns were tested against a curveball randomisation of the binary clone by archetype presence matrix, which holds fixed both the number of archetypes per clone and the number of clones per archetype, so that neither clone size nor archetype abundance can drive the result (5,000 samples, 200 burn-in sweeps, five swaps per clone per sweep). Timepoints were pooled per line by summing observed counts across timepoints while generating the null separately within each and summing that too, since the two timepoints differ sharply in composition and randomising the pooled matrix would invent enrichment for any pattern commoner at one timepoint. Enrichment is reported as the z of the observed count against this null with a two-sided permutation p and no multiple-testing correction, and a pattern is displayed only where the larger of its observed and expected count reaches three. Cross-timepoint clones were additionally drawn as a forest, grouped by the archetype set held at the first timepoint, which is the parent set the crediting rules key on, with each clone’s second-timepoint composition shown as a pie scaled to cell number. This is a time-ordered display and not a phylogeny, since a single consensus barcode signature is applied per clone and all cells within a clone are siblings.

### Re-analysis of RPM primary tumour single-cell RNA-seq data

#### Co-embedding of the *RPR2* and *RPM* tumour datasets

The published *RPM* dataset (RPM_K5vCGRP_Seurat.rds) was used with its defined clusters, Cre-driver annotation and per-cell signature scores. Cell cycle was scored separately in each object using the standard S and G2/M gene lists converted to mouse symbols. Objects were merged with dataset-unique cell names and the RNA assay split by dataset, then log-normalised, reduced to 3,000 variable features, scaled with S and G2/M scores regressed out, and taken to 30 principal components. Datasets were integrated with IntegrateLayers using CCA on log-normalised counts.

#### Cross-dataset neighbourhood signature enrichment

As the published *RPM* clusters are not labelled by lineage, each *RPR2* archetype was profiled against *RPM* signature scores rather than cluster identity. For every *RPR2* cell, the *RPM* signature scores of its k nearest *RPM* cells in the first 30 CCA dimensions were averaged, and per-cell means were averaged within each archetype. The null drew k random *RPM* cells per query cell over 1,000 permutations. Enrichment is the observed minus null mean, with a one-sided empirical p and Benjamini-Hochberg correction across all archetypes, signatures and k. Conclusions were unchanged at k = 10, 30 and 50.

#### Centroid distance in the integrated space

Centroids were computed after whitening: the pooled covariance across all cells in the first 30 CCA dimensions was inverted by generalised inverse and applied as a Cholesky transform, making Euclidean distance the Mahalanobis distance; pooled rather than per-group covariance avoided instability on the smaller *RPM* clusters. Centroid-to-centroid distances and per-cell distances to each cluster centroid were tabulated, and cosine distance (one minus cosine similarity) was computed on the same centroids as a scale-insensitive check.

#### Pathway comparison between *RPM* clusters 3 and 10 and the *RPR2* Intermediate archetype

Clusters nearest the Intermediate archetype were characterised within their own dataset. One-versus-rest differential expression used a Wilcoxon rank sum test on log-normalised counts (min.pct = 0.1, no logFC threshold), ranked by average log2 fold change; ranked lists were tested by GSEA against GO biological process terms (gseGO, org.Mm.eg.db, Benjamini-Hochberg, simplification at 0.7) and mouse Reactome sets (MSigDB) at FDR < 0.05. Intermediate terms came from the *RPR2* archetype GSEA under identical settings. One representative term per theme is shown across the three groups, dot size giving normalised enrichment score and colour the negative log10 adjusted p value.

#### Terminal-state analysis within the *RPM* dataset

Terminal-state behaviour was tested within the *RPM* data alone. Variable features, scaling and 30 principal components were recomputed from the log-normalised RNA assay of the full object (27,674 cells, 11 published clusters) and a neighbour graph built on it. A CellRank ConnectivityKernel matrix was computed and a GPCCA estimator fitted with 11 macrostates and cluster labels as the cluster key, followed by predict_terminal_states and compute_fate_probabilities. Each macrostate is defined by 30 core cells, whose cluster composition distinguishes a cluster anchoring its own macrostate from one absorbed into others; fate probabilities are shown as a circular projection.

#### Calculating RPR2 archetype signatures in human bulk RNA-seq data

*RPR2* archetype gene signatures were converted to their human orthologs using the Mouse Genome Informatics homology resource (HOM_MouseHumanSequence.rpt, Jackson Laboratory). For each mouse gene in an archetype-specific list, we retrieved the corresponding homology class and extracted all associated human symbols, yielding human NE1, NE2, Intermediate, Secretory, nonNE1, and nonNE2 signature gene sets.

#### Single-sample archetype scores in bulk RNA-seq

Bulk RNA-seq expression matrices from human tumours and matched adjacent normal tissues for 112 treatment-naïve patients from the TU-SCLC cohort were downloaded from Liu *et al.* ^19^. Additionally, a bulk RNA expression matrix for 160 tumour samples from 65 patients was downloaded from George *et al.* ^43^. Both datasets were preprocessed by standardising gene identifiers as row names and replacing missing or non-numeric entries with zeros. Single-sample gene set enrichment scores for each archetype were then computed across all samples using the GSVA R package ^91^ (method = “ssgsea”, Gaussian kernel, gene-set size between 5 and 500, and normalisation enabled), yielding a gene–set–by–sample matrix of archetype scores.

To relate archetype scores to canonical lineage and immune programs, we extracted bulk expression for key transcription factors and immune factors (e.g., ASCL1, NEUROD1, POU2F3, YAP1, MHC class I/II components, T-cell markers) and combined these with the ssGSEA-derived archetype scores into a single sample-level matrix. Pairwise correlations and corresponding P-values were estimated using Spearman correlation (‘*rcorr*’ function, *Hmisc* R package), and correlation structure was visualised with the corrplot R package using significance masking at p < 0.01 and ordered heatmaps for tumour and normal-adjacent tissue samples separately.

To assess concordance between transcript- and protein-level archetype associations, we performed an analogous analysis using matched proteomic measurements from Liu *et al*. ^19^ after harmonising protein identifiers and imputing missing values as zeros. Protein abundances for selected markers were combined with ssGSEA archetype scores, and sample-wise correlation matrices were computed again with *Hmisc* and visualised as correlation heatmaps with corrplot for the tumour and normal-adjacent cohorts, using the same significance thresholds and display settings.

#### Quantification of epigenetic plasticity score from scATAC-seq data

Epigenetic plasticity was quantified at the single-cell level by combining two complementary metrics derived from scATAC-scRNA multi-omics data: (i) a chromatin entropy score that quantifies the dispersion of accessible chromatin across peaks and (ii) a gene activity–expression decoupling score that reflects the extent to which chromatin accessibility and transcription are uncoupled in each cell.

#### Entropy score

Chromatin entropy was calculated on the ATAC assay (“peaks”) using a custom R function implemented with Signac and Seurat. The peak-by-cell accessibility matrix was extracted from the data layer, and variable peaks were identified either from pre-existing variable features or, when absent, by (i) ranking peaks by variance across cells if the counts layer was empty or (ii) applying FindTopFeatures with a 5% quantile cutoff when counts were available. The matrix was restricted to the top 5,000 variable peaks, and for each cell, we computed Shannon entropy over its normalised peak accessibility profile using the following formula:

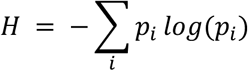

where *p_i_* is the fraction of total accessibility at peak *i*; cells with zero total accessibility were assigned an entropy of 0.

Entropy values were linearly rescaled to a 0–1 range and were added to the metadata as “chromatin entropy” (raw) and “chromatin entropy norm” (normalised).

#### Decoupling-based plasticity score

To quantify regulatory decoupling between chromatin accessibility and transcription, we compared per-cell gene-activity profiles from scATAC-seq with matched gene-expression profiles from scRNA-seq for all archetype marker genes with per-cell gene-activity data. For each cell, we calculated a Spearman correlation between gene activity and expression across these markers, interpreted this correlation as an “activity expression coupling” score, and then defined a decoupling-based plasticity score as 1-correlation, such that higher values reflect stronger epigenetic–transcriptional decoupling.

For each cell, a composite plasticity score was computed by z-scoring and summing the normalised chromatin entropy and decoupling-based plasticity scores, as shown in the archetype distribution in Fig. 3f.

#### Analysis of intermediate cancer cell states

Intermediate cancer cells were subset from the full multi-omics dataset and re-embedded using the WNN framework as described above (RNA harmony_2 dimensions 1–30, ATAC harmony_1 dimensions 2–30), with Leiden clustering performed at resolution 0.25. Palantir-derived pseudotime coordinates were transferred to the Seurat object metadata, and cells were stratified into four bins based on Palantir pseudotime scores: Early ( < 0.10), Transitioning Early (0.10– 0.50), Transitioning Late (0.50–0.75), and Late (0.75–1.0). Differential gene expression analysis was performed between the pseudotime bins using the FindAllMarkers() function in Seurat on the normalised SCT counts, with the Wilcoxon test (logFC threshold = 0.25, p_val_adj < 0.01, and min.pct = 0.1). Pearson correlations between pseudotime and the SCLC-I signature were computed for each bin and visualised as a scatter plot. Per-cell epigenetic regulation scores (GOBP_EPIGENETIC_REGULATION_OF_GENE_EXPRESSION, MSigDB C5) were computed using AddModuleScore on SCTransform-normalised counts (ctrl = 300, nbin = 12), and differences across pseudotime bins were assessed using pairwise Wilcoxon rank-sum tests with Benjamini-Hochberg correction (ref.group = “Mid_Early”).

To reconstruct the transcriptional trajectory of intermediate cells, the Seurat object was converted to a Monocle3 ^44^ cell_data_set comprising SCTransform assay and WNN UMAP coordinates. A principal graph was learned using learn_graph (use_partition = TRUE), and the trajectory root was defined programmatically by identifying the principal graph node closest to the centroid of “Early” cells. Pseudotime-variable genes were identified using graph_test (neighbour_graph = “principal_graph”, q < 1×10⁻⁵), and gene modules were defined using find_gene_modules across a resolution sweep (10^seq (−6, −1)).

#### Identification of domains of regulatory chromatin and gene regulatory network inference

To identify *cis*-regulatory elements and infer transcription factor (TF) activity at single-cell resolution, the FigR package ^46^ was implemented on the dataset. Peak ranges from the chromatin accessibility assay were first resized to 300 bp around their centres and trimmed to chromosome boundaries to standardise peak width, and a SummarizedExperiment object was then constructed from the resulting accessibility matrix, alongside SCTransform-normalised RNA expression data, with non-expressed genes excluded. *Cis*-regulatory peak–gene associations were computed with runGenePeakcorr, using 50 background peak pairs per gene–peak association for significance testing. Peak–gene pairs with a Z-test P-value ≤ 0.05 were retained for downstream analyses. Genes with five or more significantly correlated peaks were classified as domains of regulatory chromatin (DORCs), and DORC accessibility scores were computed per cell using getDORCScores. A TF–DORC gene regulatory network was inferred using runFigRGRN, which integrates TF motif enrichment among DORC-associated peaks with correlation between TF expression and DORC accessibility scores to nominate likely TF activators and repressors. TF– DORC regulatory scores were computed across all expressed TFs and DORC genes, and 131 TF regulators with an absolute regulatory score greater than 1.25 were initially shortlisted from this analysis. These were further filtered based on their expression in the single-cell dataset, and 68 transcription factors expressed in at least 10% of cells were retained for the BoBa-T network. The heatmap in Fig. 4d was generated by correlating the normalised SCT counts of the 68 transcription factors across all cells in the multi-omics data.

#### DIRECT-NET Analysis

To map *cis*-regulatory element (CRE)–gene relationships for transcription factors identified as FigR DORC regulators, DIRECT-NET ^92^ analysis was performed on the multi-omics Seurat object using the DIRECT-NET package. Genomic coordinate information was derived from mm10 gene transcription start site annotations, and DORC-associated TFs were used as focal markers. DIRECT-NET was run with binary ATAC accessibility, a cell neighbourhood size of 50, and a maximum peak overlap of 0.5, yielding per-TF regulatory link scores between accessible chromatin regions and target genes. Regulatory links were annotated as high-confidence (HC) or modulatory (MC) based on link function type and stored for downstream intersection analyses.

CRE–gene links from DIRECT-NET were then separately extracted for distal and promoter-associated regulatory regions using generate_CRE_Gene_links, with each TF assigned to its most enriched cell cluster based on prior DORC analysis. Distal and promoter CRE–gene links were subsequently intersected with the differentially accessible (DA) peaks (described above) to retain only regulatory regions showing significant differential accessibility across cell states.

TF motif enrichment within the DA-filtered CRE–gene linked peaks was assessed using motifmatchr against the JASPAR2020 CORE vertebrate motif database, with motif scanning performed on the mm10 genome (BSgenome.Mmusculus.UCSC.mm10). Motif score matrices were filtered to retain only peaks with non-zero motif scores, and columns were restricted to the subset of DORC TFs whose names were present in the JASPAR2020 motif database, supplemented by a curated set of additional regulators of interest (for example, RORA, SOX9, REST, ZBTB7A, and HES1). The resulting peak-by-TF motif score table, with each row representing a DA regulatory region linked to a target gene, was exported for downstream network visualisation and interpretation.

#### Network structure from DIRECT-NET

Using the DIRECT-NET output [SG1.1] with a significance threshold of 0.1 and the 2020 TF database as the reference, we built an initial network of all connections, including their motif scores. To reduce network complexity, for any nodes with more than 8 regulators, we used a Lasso regression approach as follows. For each target node in the network, the expression values of its candidate regulators were used as predictors, with the target expression as the response variable. Data were partitioned into training and test sets (80%/20% respectively). Optimal regularisation strength was identified via 5-fold cross-validated grid search across a range of alpha values (1e-5, 1e-4, 1e-3, 1e-2, 1e-1, 1). For each node, regulators with non-zero LASSO coefficients were retained as significant predictors for each target, and the lowest alpha value that yielded < 8 regulators was selected. The final pruned network was compiled into a combined edge list as a preliminary network for BoBa-T inference. Sinks were removed except ZBTB20, KMT2A, SIX4, STAT2, NFKB1, EPCAM, NCAM1, ICAM1, and SOX11 as readouts of cell state. Nodes without parents have a self-loop added for inference, but all other nodes have self-loops removed.

RORA and RORB were combined into a single node, as their binding motifs are indistinguishable [SG2.1]. This resulted in a preliminary network structure for rule inference.

#### Regulatory rule inference

Inference of regulatory rules was done using BooleaBayes 1.0.0 as described previously in Wooten, Groves, *et al.* ^53^ (https://pypi.org/project/booleabayes/). Briefly, data were binarised (0 and 1) using quantile normalisation, with thresholds q = 0.3 and q = 0.7 for 0 and 1, respectively. Rules were fit on the binarised data using the network structure’s vertex dictionary, with node_threshold = 0 (no parents removed). In-sample validation of the rules was done using the fit_validation method of BooleaBayes.

#### Destabilisation predictions

To quantify the effects of activating or inhibiting each node in the network, BooleaBayes was used to find attractors, with threshold = 0.5, within a basin of 2 from the single-cell data (see Wooten, Groves, *et al.* ^53^ for more details). Random walks from each identified attractor were performed until the walk exited the attractor basin of 2, or reached 500 steps, with and without each perturbation independently. Destabilisation score d was defined as

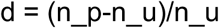

Where n_u is the average steps to leave the basin for an attractor in the unperturbed simulation and n_p is the equivalent in the perturbed simulation.

### Analysis of Control and Rorb Knockdown single cell data

#### Preprocessing

scRNA-seq data from SCLC organoids were generated using the Parse Biosciences platform and processed in R using Seurat. The filtered digital gene expression matrix was imported using ReadParseBio, and a Seurat object was constructed, retaining genes detected in at least 3 cells and cells with a minimum of 100 detected features. For quality control, mitochondrial read percentage was calculated, and cells were retained if they had between 500 and 6,000 detected genes, fewer than 20,000 UMI counts, and less than 15% mitochondrial reads, thereby removing low-quality cells and putative empty droplets. The RNA assay was split by sample to enable sample-aware integration, and count data were normalised using SCTransform (glmGamPoi backend), with mitochondrial content regressed out.

Following normalisation, principal-component analysis was performed, and sample integration was carried out using reciprocal PCA (RPCA) via Seurat’s IntegrateLayers() function (30 dimensions, k.anchor = 20) to correct for sample-level batch effects while preserving biological variation across experimental conditions. Shared nearest-neighbour graph construction and Leiden clustering (resolution 0.2) were performed on the RPCA-corrected embedding, and a cluster tree was built and reordered numerically for visualisation. Cells were embedded in a two-dimensional UMAP space via RPCA for downstream visualisation and annotation.

#### Cell type label transfer from multi-omics reference to organoid data

To assign archetype-based cell type labels to the organoid single-cell RNA-seq data, label transfer was performed from the *RPR2* multi-omics dataset as a reference using Seurat’s anchor-based transfer framework. Before transfer, non-lineage-committed cell populations (Generalist NE and Generalist nonNE) were excluded from the reference, and the remaining cells were downsampled to 300 cells per cell type to ensure balanced representation across archetypes. Transfer anchors between the reference and organoid query dataset were identified with FindTransferAnchors on the SCT-normalised data, using the Harmony-corrected PCA embedding of the multi-omics data as the reference reduction, archetype differential genes as the feature set, and non-default parameters of k.anchor = 10, n.trees = 100, and the RANN nearest-neighbour method (30 dimensions). Predicted cell type labels and related confidence scores were computed with TransferData and stored in the organoid Seurat object metadata for downstream analyses.

#### Differential abundance analysis using Milo

To identify cell neighbourhoods differentially abundant between knockdown and control conditions, Milo analysis was performed using the miloR ^57^ package (v2.5.1). Given the large dataset size, a random subset of 15,000 cells was used for initial analyses. The Seurat object was converted to a SingleCellExperiment and then to a Milo object using raw RNA counts, with the RPCA-corrected embedding used to construct the k-nearest-neighbour graph (k = 50, d = 30). Cellular neighbourhoods were defined using refined sampling at a proportion of 0.1, and cells were counted per sample within each neighbourhood to generate the neighbourhood count matrix. Each sample was assigned to one of five organoid lines, and a design matrix was constructed with knockdown condition (sh*Gfp* as control, sh*Rorb*#1, and sh*Rorb*#2 as KD) and line as covariates. To account for line-to-line variation, two complementary statistical models were tested: a fixed-effects model including line and condition as additive terms, and a mixed-effects model treating line as a random effect with condition as the fixed effect. Both models were tested using graph-overlap spatial FDR weighting, and neighbourhoods were considered significantly differentially abundant at a spatial FDR threshold of 0.1. Given the strong inter-line variation observed in initial analyses, Harmony integration was additionally performed on the PCA embedding using organoid line as the grouping variable (max 20 iterations), and Milo analysis was repeated on the Harmony-corrected embedding (k = 50, d = 30) with a fixed-effects model including line and KD condition as covariates. Differentially abundant neighbourhoods were annotated by their predominant RPCA cluster or transferred archetype identity, with neighbourhoods assigned a “mixed” label if no single cell type comprised more than 70% of the neighbourhood. Results were visualised as beeswarm plots stratified by cluster or predicted archetype identity, and neighbourhood DA scores were overlaid on the UMAP embedding for spatial context.

#### Gene set enrichment analysis and pathway activity scoring

To quantify pathway activity at single-cell resolution, enrichment scores for a curated panel of gene sets were computed using the irGSEA framework on log-normalised RNA counts. Gene sets were assembled from the MSigDB^90^ Hallmark collection and Gene Ontology Biological Process (GO: BP) collection for mouse, obtained via the msigdbr package (https://igordot.github.io/msigdbr/), encompassing pathways related to immune response, cellular stress, proliferation, metabolism, and signalling (for example, interferon response, inflammatory response, MYC targets, UPR, glycolysis, and antigen processing). Per-cell enrichment scores were calculated in parallel using three complementary methods, AUCell, UCell and ssGSEA (Gaussian kernel), to ensure robust estimation of pathway activity across cell states. Differential pathway activity across cell clusters was assessed using irGSEA.integrate, which consolidates results across scoring methods and applies robust rank aggregation (RRA) to identify gene sets consistently enriched or depleted across conditions. For each method, enrichment scores, normalised enrichment scores (NES), nominal P-values, and adjusted P-values were extracted per cluster. Gene sets were considered significantly enriched at an adjusted P-value threshold of 0.05. For visualisation, mean ssGSEA scores per cluster were computed and arranged in a pathway-by-cluster matrix. Columns were ordered to reflect biological groupings of cell clusters, and rows were hierarchically clustered. The resulting matrix was scaled by row and visualised as a heatmap using pheatmap, with row annotations indicating the significance tier of each pathway derived from the irGSEA differential analysis.

### MIBI analysis

#### Imaging and Preprocessing

Tissue sections were imaged using Multiplexed Ion Beam Imaging by Time-of-Flight (MIBI-TOF). Raw image data were extracted from bin files and processed using the toffy pipeline (https://github.com/angelolab/toffy). Background signal contamination and channel crosstalk were corrected using Rosetta, a flow cytometry-style compensation algorithm. Images were then normalised to account for variations in detector sensitivity across the run using median pulse-height calibration. Whole-cell segmentation was performed with DeepCell ^93^, and per-cell marker expression probabilities were generated using Nimbus Inference ^94^.

#### Spatial Compartment Assignment

Tumour and stromal regions were manually annotated in QuPath^29^ using overlays of ASCL1, Synaptophysin, ICAM1, Vimentin, and Histone H3 (HH3). These masks were used to partition cells into two spatial compartments: cancer and stroma. From these binary stroma masks, additional border and core spatial compartments were computed using Euclidean distance transforms. Cells within 100 pixels of the tumour-stroma boundary were classified as border regions (stroma border, cancer border); cells beyond this threshold were classified as core regions. This yielded four compartments: stroma core (> 100 pixels within the stroma), stroma border (≤ 100 pixels within the stroma), cancer border (≤ 100 pixels within the tumour), and cancer core (> 100 pixels within the tumour). Cells were assigned to compartments by majority vote of their constituent pixels.

#### Cell Phenotyping

FlowSOM ^95^ clustering was performed separately on cells from the stromal and cancer compartments, using Nimbus expression probabilities for ASCL1, Synaptophysin, ICAM1, Vimentin, and CD45. Stromal compartment cells were annotated as stroma or immune. Cancer compartment cells were classified as Neuroendocrine, Intermediate, Non-Neuroendocrine, Stroma, or Immune based on cluster marker profiles. Ambiguously clustered cells were re-clustered on a finer grid to improve assignment.

#### Distance-Based Expression Analysis

Euclidean distances from each cancer cell to the nearest stromal boundary were computed in microns. Cells were grouped into 25 µm bins out to 250 µm. Mean expression of 27 functional markers was calculated per bin. Values were row-normalised and hierarchically clustered to identify co-regulated marker modules.

#### Statistical Comparisons

Differences in marker expression between the cancer core and border were tested using t-tests with FDR correction. Significance was defined as adjusted p < 0.05 and |log₂FC| ≥ 0.5.

#### Statistics and reproducibility

Statistical significance was assayed with GraphPad Prism software for *in vivo* studies. Data are represented as mean ± s.e.m. Tests used are indicated in figure legends. No data were excluded from the analysis. To compare two groups, we used a t-test. The effect size between distributions was compared using Cliff’s delta. To compare growth curves, we used two-way ANOVA.

## Data and code availability

Sequencing data from single-cell experiments are available from the Gene Expression Omnibus (GEO) under accession number GSE325242. All code used for generating the figures, as well as the code for BoBa-T, has been deposited on GitHub (https://github.com/smgroves/Bhattacharya2026). All other data are available in the article and supplementary materials or from the corresponding author upon reasonable request.

## ACKNOWLEDGMENTS

The authors thank all members of the Sage lab for technical assistance and support throughout this work, as well as Dr. Dian Yang, Dr. Joanna Wysocka, and Dr. Or Gozani for helpful feedback. This work was supported by the Damon Runyon Cancer Research Foundation (DRG-2466-22, post-doctoral fellowship to D.B.), the NCI (F31 fellowship to G.G.H., and R35 award to J.S.), and the Ludwig Center at Stanford (J.S.).

## DECLARATION OF INTERESTS

J.S. has equity in, and is an advisor for, DISCO Pharmaceuticals. The other authors declare no competing interests.

---

**Supplementary Figure 1:**
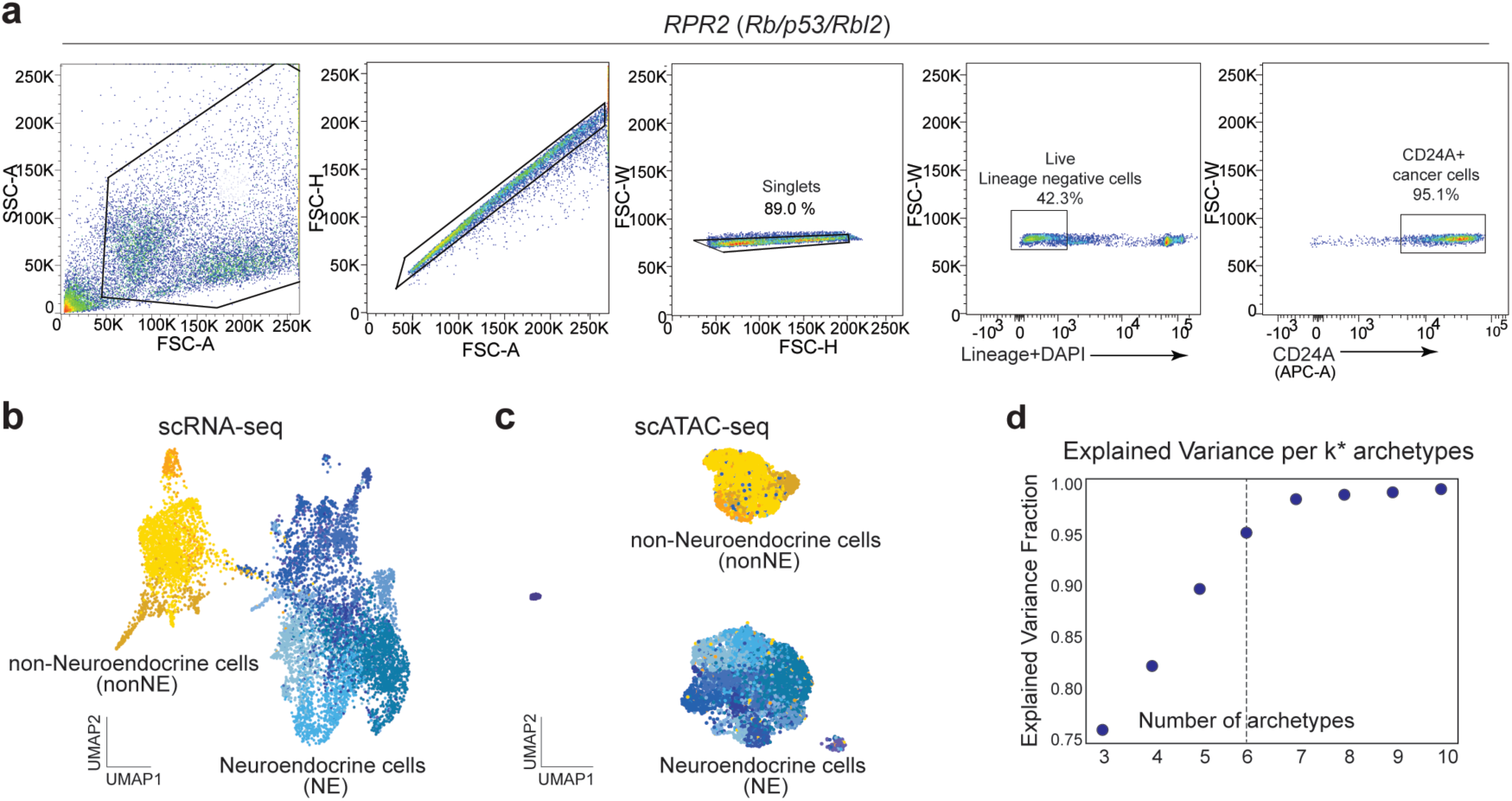
Single-cell multi-omics of SCLC cancer cells. **a.** Flow cytometry strategy to isolate cancer cells from primary lung tumours of *RPR2* mutant mice (as in Shue *et al.*, 2022). **b,c.** Uniform manifold approximation and projection (UMAP) depicting the scRNA-seq (b) and scATAC-seq (c) modalities of the multi-omics data. Neuroendocrine (NE) cell populations are shown in blue, while nonNE cells are shown in yellow. **d.** Knee plot showing the optimal number of archetypes required to explain the variance of the top 10 principal components of the scRNA-seq data.

**Supplementary Figure 2:**
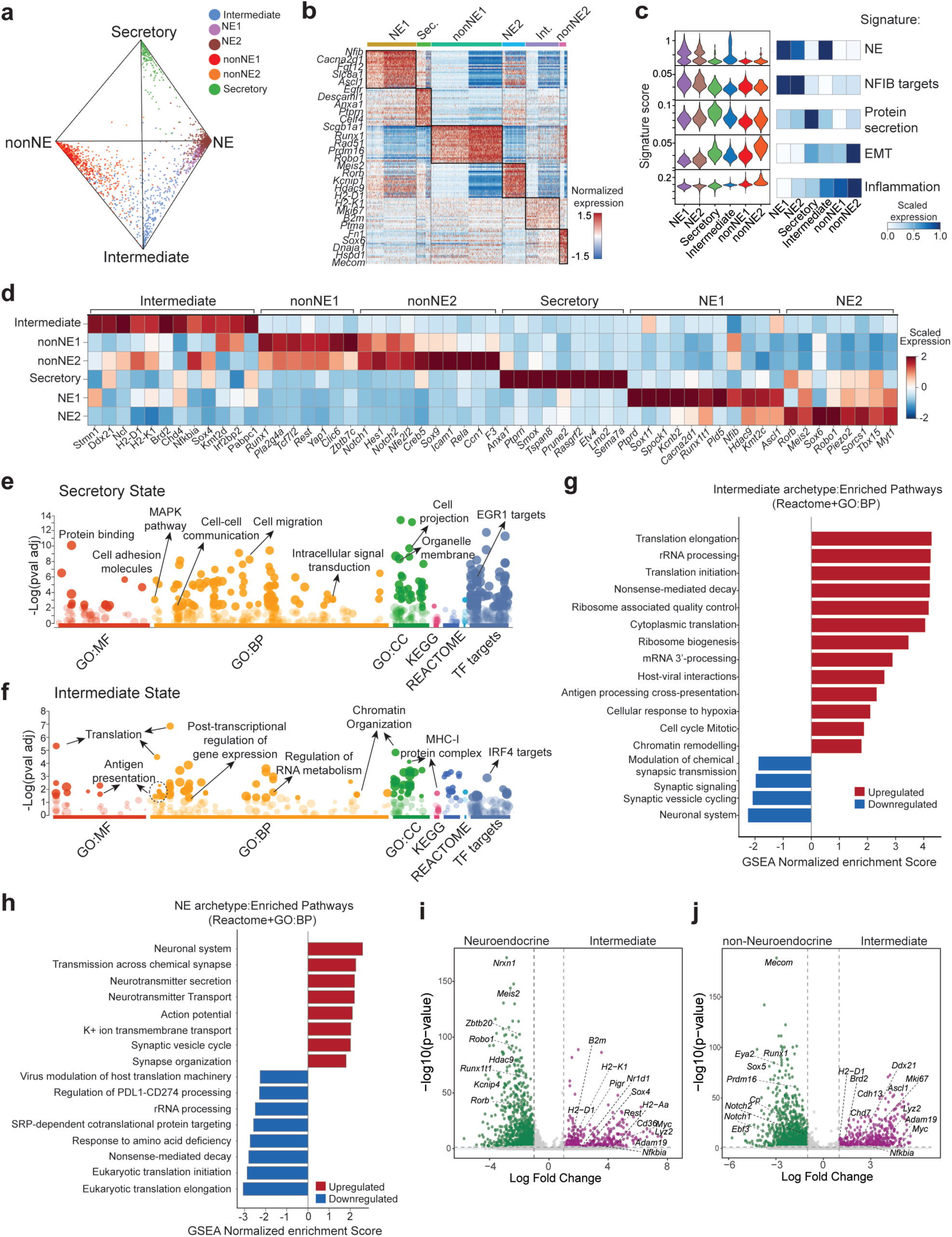
Molecular signatures defining SCLC archetypes. **a.** CytoSimplex quaternary plot of cancer cells in *RPR2* tumours, positioned by transcriptional similarity to the NE, nonNE, Secretory and Intermediate vertices. Cells near a vertex are Specialists of that state; cells at the centre express mixed programs. Intermediate cells occupy their own vertex rather than the centre. **b.** Heatmap showing expression of top 20 marker genes for each archetype across cancer cells, with selected representative markers labelled for each cluster. **c.** Activity of SCLC-relevant gene signatures across *RPR2* cancer cell archetypes. **d.** Heatmap showing expression (Z-score) of the top 10 marker genes of each archetype. **e,f.** Bubble plot showing significantly enriched (p < 0.01) gene programs associated with the low NE archetypes: Secretory (e) and Intermediate (f). Select gene programs of each cell state are highlighted. **g.** Diverging bar plots showing curated list of top upregulated and downregulated Gene Ontology: Biological Processes (GOBP) and Reactome pathways in the Intermediate archetype as determined by Gene set enrichment analysis (GSEA). **h.** Diverging bar plots showing curated list of top upregulated and downregulated Gene Ontology: Biological Processes (GOBP) and Reactome pathways in the NE archetypes (NE1+NE2) as determined by Gene set enrichment analysis (GSEA). **i.** Volcano plot identifying differentially expressed genes between classical NE cells (NE1 & NE2) and Intermediate cancer cells. **j.** Volcano plot identifying differentially expressed genes between classical nonNE cells (nonNE1 & nonNE2) and Intermediate cancer cells.

**Supplementary Figure 3:**
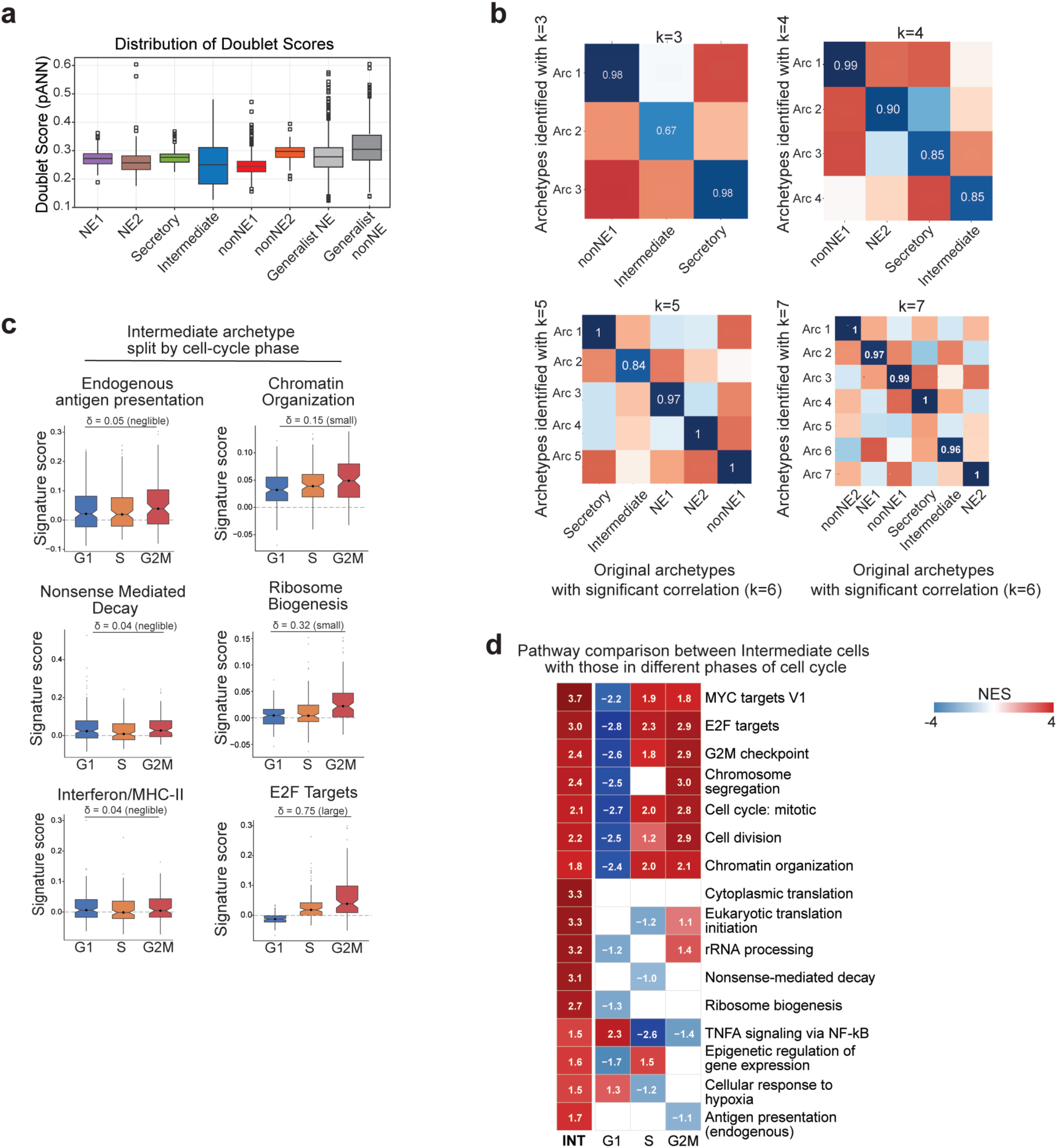
Validation and molecular characterisation of SCLC archetypes. **a.** Distribution of DoubletFinder pANN score of each archetype, calculated based on scRNA-seq data modality. **b.** Correlation between archetypes for k = 3, k = 4, k = 5, and k = 7 compared to k = 6. Only most correlated archetypes (of k = 6) are shown in each panel. An archetype strongly correlated to Intermediate was identified across all values of k. **c.** Activity of pathways characteristic of the Intermediate state, in Intermediate cells grouped by cell-cycle phase. Most pathways are comparable across phases, except for “E2F Targets”, included as a positive control. Cliff’s delta value (δ) was calculated to measure the effect size between non-parametric distributions. **d.** Heatmap of differential enrichment of Intermediate-specific pathways in *RPR2* cancer cells grouped by cell cycle phases. Pathways associated with cell cycle progression are concordantly upregulated in Intermediate and G2/M cells, while those associated with translational control, inflammation and hypoxia are cell cycle agnostic. Only significant enrichments (FDR < 0.05) are shown. INT, Intermediate.

**Supplementary Figure 4:**
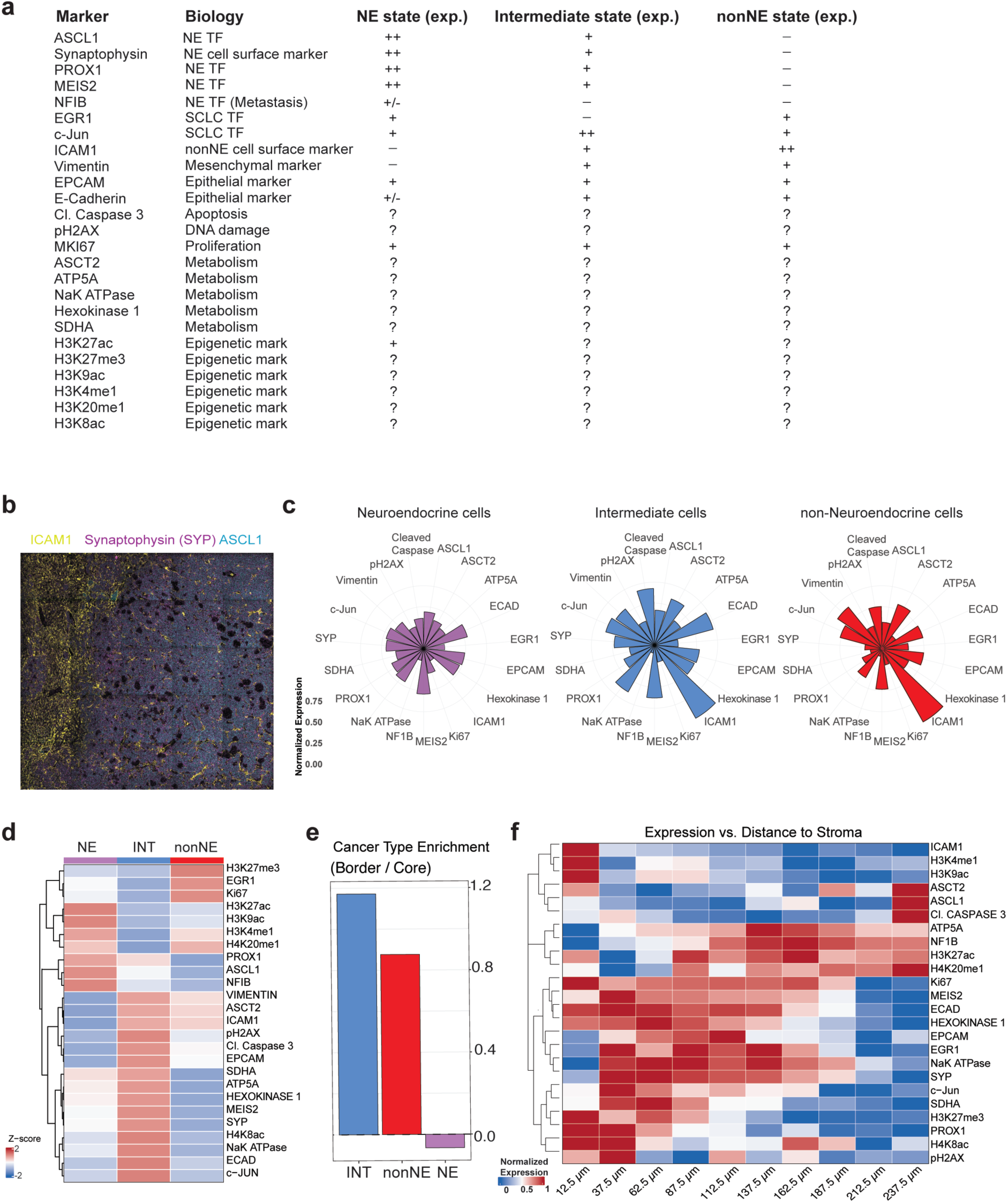
MIBI analysis of *RPR2* mutant tumours. **a.** Markers used in the MIBI analysis and their expected (exp.) signal in the three predominant SCLC cancer lineages: Neuroendocrine (NE), Intermediate (INT), and non-Neuroendocrine (nonNE). **b.** Representative image with pseudo colours for ICAM1, Synaptophysin (SYP), and ASCL1 in a primary lung tumour section from the *RPR2* model. **c.** Radial plot summarising the defining gene and feature signatures of each SCLC cancer cell lineage identified in MIBI. The length of each bar corresponds to the relative enrichment (Z-score across subsets) of the mean value for each feature. **d.** Heatmap showing the scaled signal of each protein marker in the three SCLC cell states. **e.** Plot depicting the ratio of cells at the tumour border vs the core for each cancer cell lineage. **f.** Similar analysis as in (e) with specific markers. The distance to the tumour stroma is shown at the bottom in µm.

**Supplementary Figure 5:**
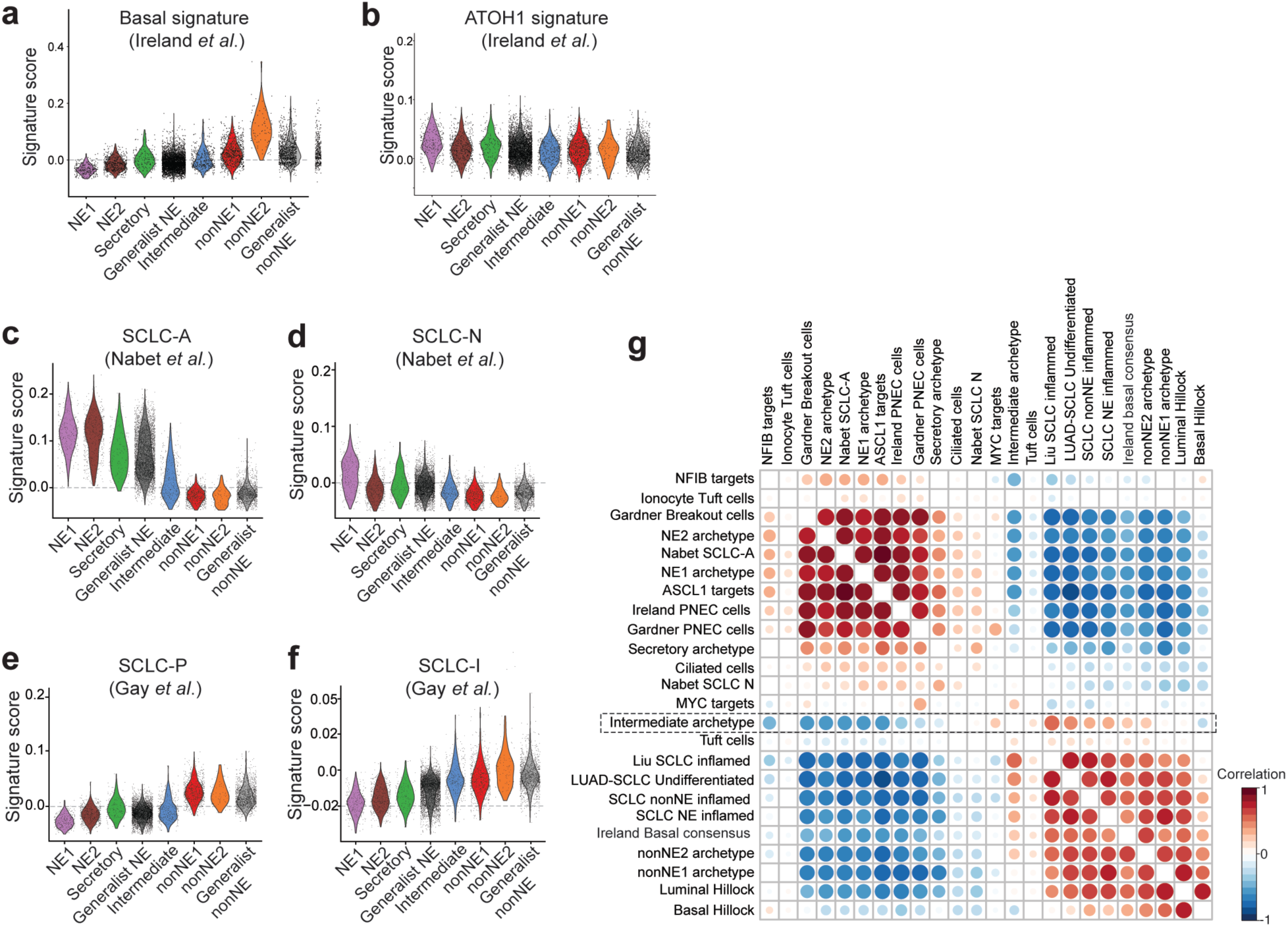
Comparative analysis of SCLC signatures across *RPR2* archetypes. **a,b.** Violin plot showing the activity of Basal signature (a) and Atoh1 (b) signature from Ireland *et al.*, across *RPR2* archetypes. **c-f.** Violin plot showing the activity of human gene signatures for SCLC-A (c), SCLC-N (d), SCLC-P (e) and SCLC-I (f) subtypes across *RPR2* cancer cell archetypes. The name of the first author on the corresponding publication from which the signature is obtained is shown. **g.** Unbiased hierarchical clustering of pairwise correlations between published SCLC gene signatures and the archetype signatures defined here, scored in the *RPR2* scRNA-seq data. The Intermediate signature (dashed box) clusters separately between the NE and nonNE signature groups. The name of the first author on the corresponding publication from which the gene signature is obtained is shown.

**Supplementary Figure 6:**
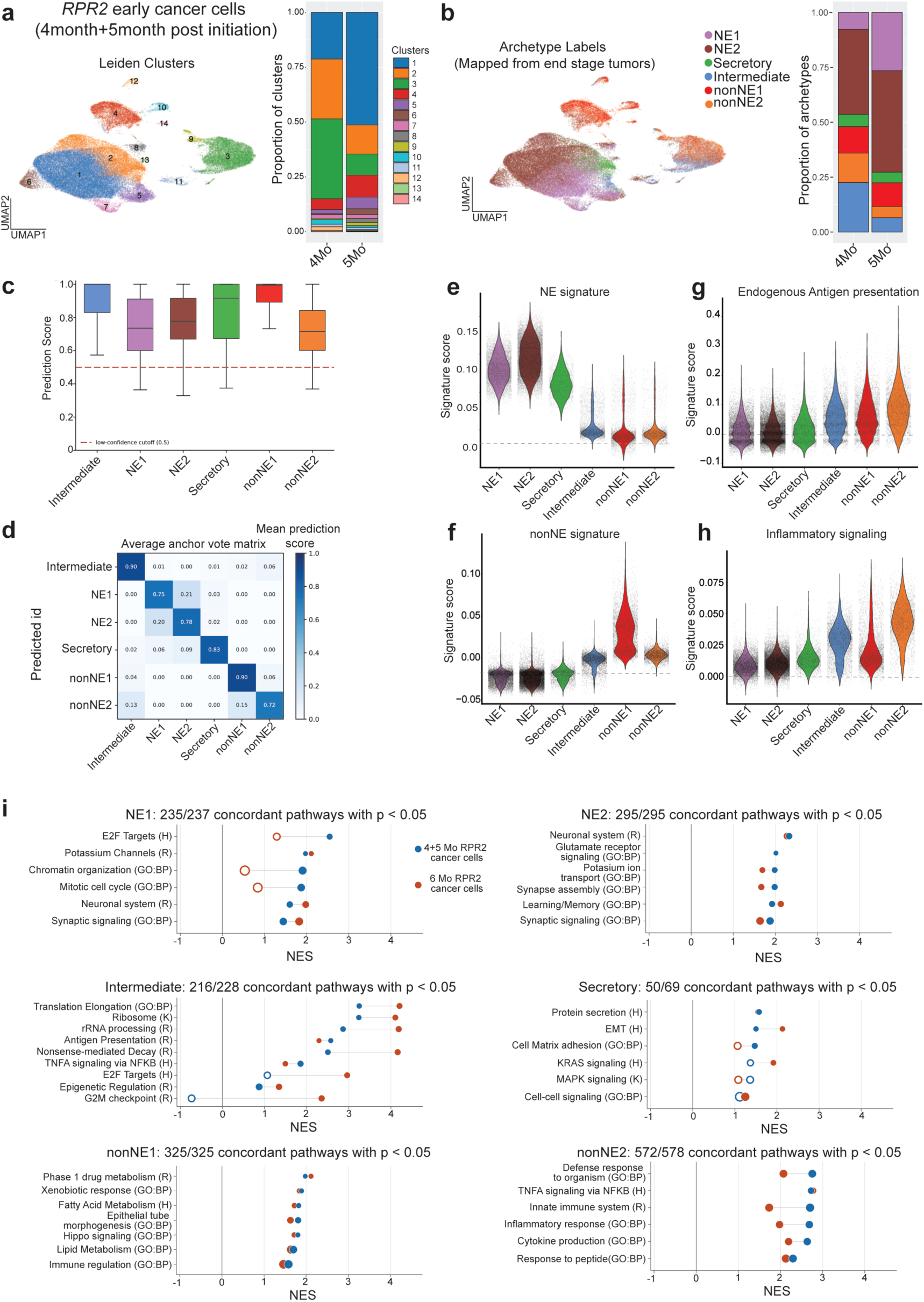
Archetypes and gene programs of *RPR2* cancer cells in tumours at 4 and 5 months post initiation. **a.** Uniform manifold approximation and projection (UMAP) of the combined scRNA-seq of cancer cells sorted from *RPR2* tumours 4-month and 5-month post initiation, coloured by Leiden clusters (left). Stacked bar plots comparing cluster composition of cancer cells from 4-month and 5-month tumours (right). **b.** Uniform manifold approximation and projection (UMAP) of the combined scRNA-seq of cancer cells sorted from *RPR2* tumours 4-month and 5-month post initiation, coloured by archetype labels mapped from end-stage tumour dataset (left). Stacked bar plots comparing archetype composition of cancer cells from 4-month and 5-month tumours (right). **c.** Prediction scores from Seurat anchor-based label transfer, using the end-stage (6 month) scRNA-seq dataset as the reference to assign archetype labels to cells in the 4- and 5-month datasets. Boxplots show the distribution of scores for cells assigned to each archetype; higher scores indicate more confident assignment. **d.** Mean anchor-vote score matrix for the transfer of 6-month archetype labels onto the 4- and 5-month cells. Cells are grouped by the label they were assigned (rows), and each column gives the mean confidence those cells received for one reference archetype; scores sum to 1 across a row, so a diagonal-heavy matrix indicates that each assigned population was supported by its own label rather than split between archetypes. **e-h**. Violin plot showing the activity of NE signature (e), nonNE signature (f), Endogenous Antigen presentation (GO: BP) (g), and Inflammatory signalling (Hallmark) (h), across cancer archetypes in the 4- and 5-month combined datasets. **i.** Pathway themes recovered by the transferred archetype labels, comparing the *RPR2* 4- and 5-month cells (query) against the 6-month reference, one panel per archetype. Each row is a curated theme, with the two dots giving its normalised enrichment score (NES) in the reference and the query and the line spanning the difference; dots are filled at nominal p < 0.05 and hollow otherwise, and dot area scales with gene set size. Panel headings give the number of shared gene sets and how many jointly significant sets agree in direction.

**Supplementary Figure 7:**
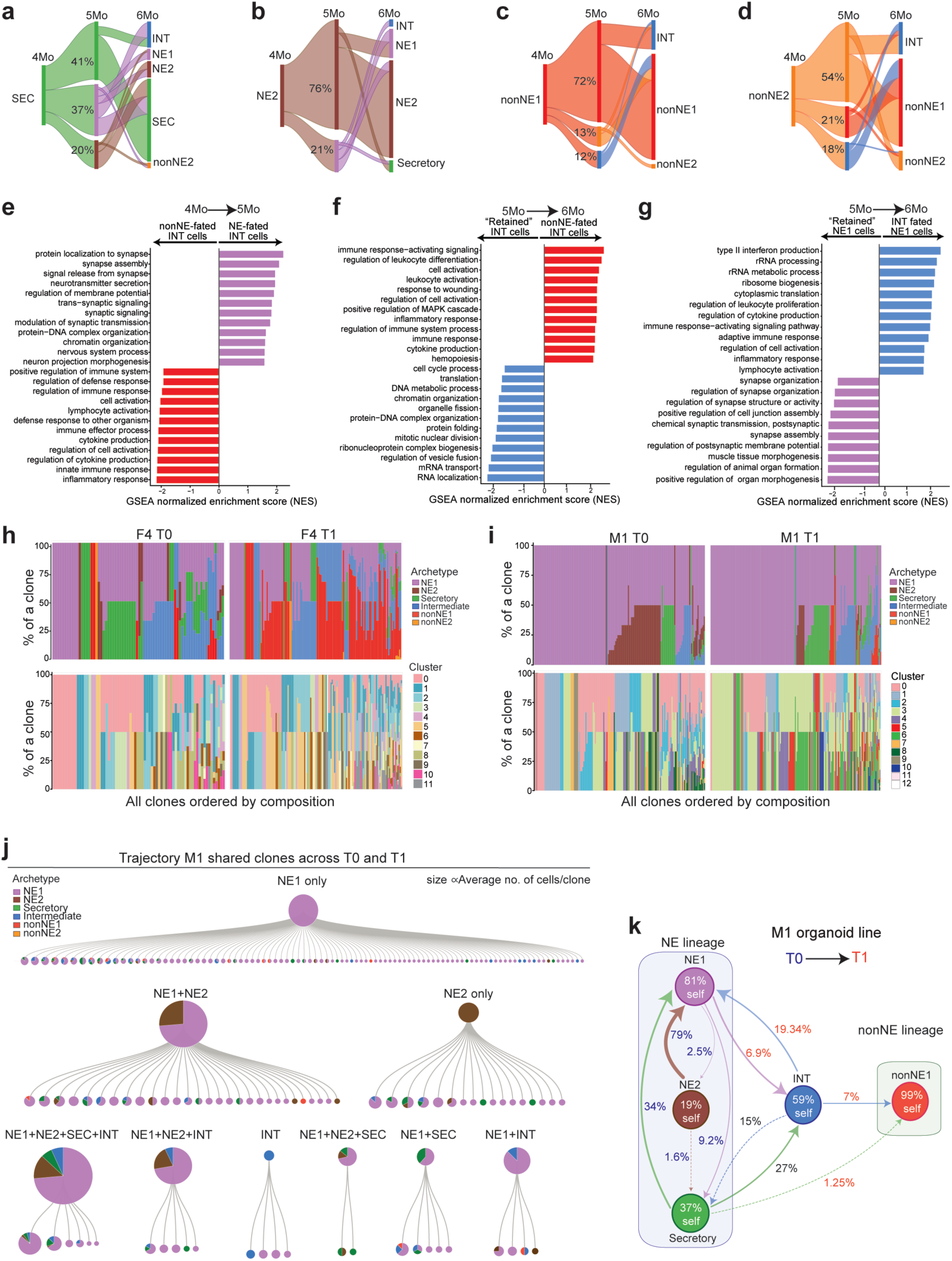
Temporal and clonal dynamics of SCLC cell-state transitions in the *RPR2* model. **a-d.** Alluvial flow plots from 4-month Secretory (a), NE2 (b), nonNE1 (c), nonNE2 (d). Ribbon width is proportional to cell mass, and percentages at 5-month show the fraction of the source’s 4-month cells taking each route; routes below 5% are omitted at both hops. **e.** Pathways enriched in 4-month Intermediate cells by predicted fate at 5 months towards either NE or nonNE. Diverging bar plots show normalised enrichment scores for each pathway. Bars to the right are enriched in NE-fated Intermediate cells, while those on the left are enriched in nonNE-fated Intermediate cells. **f.** Pathways enriched in 5-month Intermediate cells by predicted fate at 6 months, either towards nonNE or self-renewal (retained). Bars to the right are enriched in Intermediate cells that self-renew, while those on the left are enriched in nonNE-fated Intermediate cells. **g.** Pathways enriched in 5-month NE cells by predicted fate at 6 months, either towards Intermediate or self-renewal (retained). Bars to the right are enriched in NE cells that self-renew, while those on the left are enriched in Intermediate-fated NE cells. **h.** Stacked barplots showing the archetype composition (top) and cluster composition (bottom) of individual clones in the F4 organoid line at time T0 and T1. **i.** Stacked barplots showing the archetype composition (top) and cluster composition (bottom) of individual clones in the M1 organoid line at time T0 and T1. **j.** M1 clones shared across timepoints T0 and T1 (172 clones), grouped by the set of archetypes they contained at T0 and ordered left to right by the proportion of nonNE cells they reached at T1; groups represented by a single clone are omitted. The upper pie of each group gives the pooled composition of its founding clones, and each lower pie one individual clone at T1. **k.** Cell-state flux in M1 organoids, combining moscot-inferred transitions with the CellTag lineage evidence. Every percentage is moscot flux, i.e. the fraction of the source state’s T0 cells reaching that destination. Label colour indicates whether that clonal evidence came mainly from T0 (blue), mainly from T1 (red), or both (black).

**Supplementary Figure 8:**
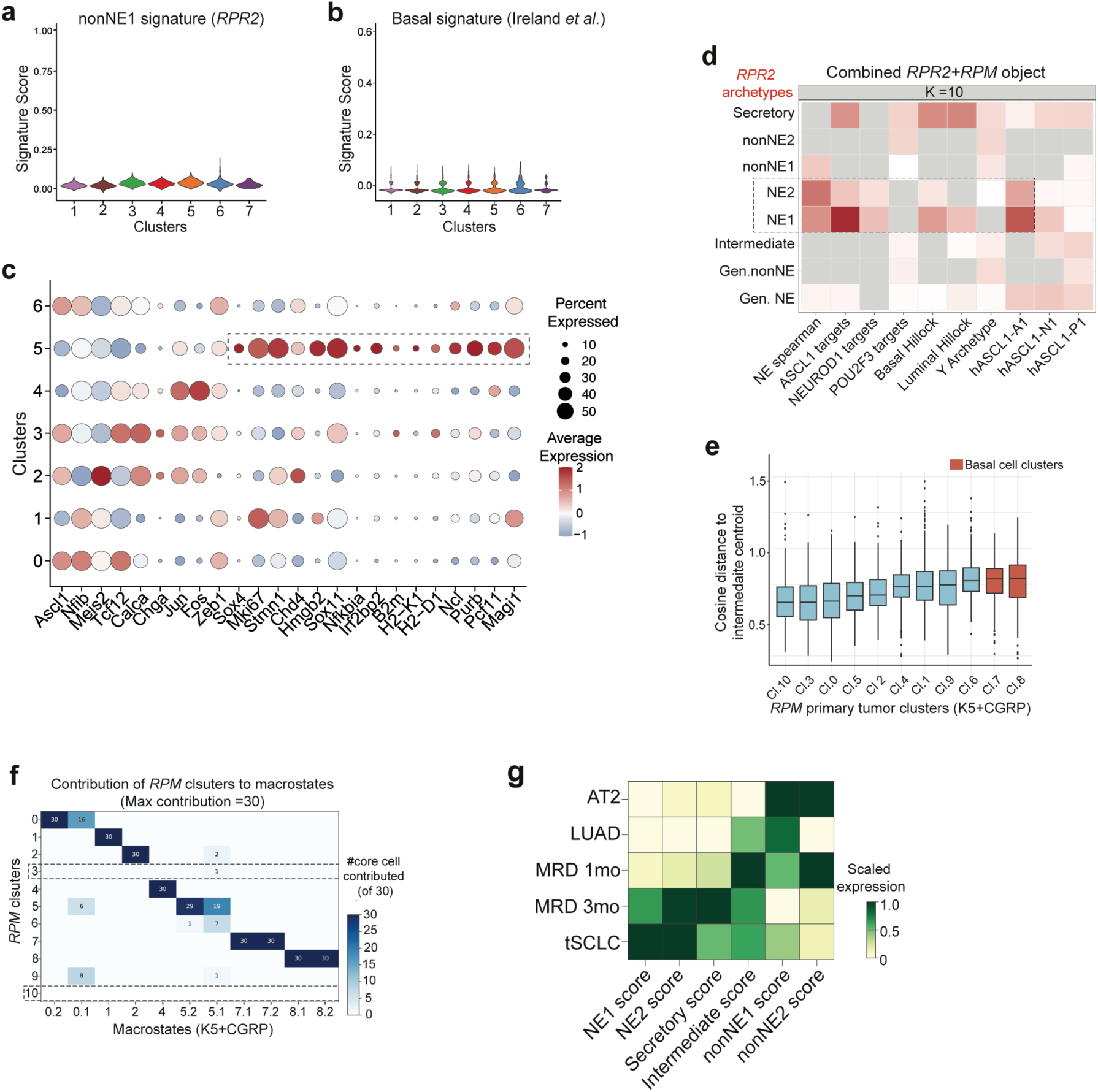
Presence of the Intermediate cell state across mouse models of SCLC. **a,b**. Violin plots of nonNE1 archetype (a) and Inflamed Basal (b) signature activity across liver metastasis clusters. **c.** DotPlot showing the expression of neuroendocrine and Intermediate marker genes in liver metastases clusters. Genes highlighted within the box are top markers of cluster 6 that are shared with Intermediate. **d.** Heatmap showing the enrichment of SCLC lineage signatures in the k = 10 nearest *RPM* cells to every *RPR2* archetype cell, in the integrated canonical correlation analysis (CCA) space of the combined *RPR2-RPM* primary tumour dataset. Colour is the enrichment over chance, and grey marks signatures not significantly enriched. The nearest *RPM* cells to the *RPR2* neuroendocrine archetypes (NE1 and NE2) are enriched for neuroendocrine signatures, serving as a positive control for successful dataset integration. **e.** Boxplots showing the spread of cosine distance of every *RPM* cluster from *the RPR2* Intermediate archetype in the CCA integrated space of the combined *RPM-RPR2* object. Clusters are arranged based on nearest to furthest distance from Intermediate cells, with clusters 3 and 10 being the closest and basal-like clusters 7 and 8 being the furthest. **f.** Heatmap showing which *RPM* cluster contributes the core cells of each terminal state in the combined K5-CGRP object, with terminal states named for the cluster they derive from. Each macrostate is defined by 30 core cells, and each tile indicates how many of those come from that cluster; empty rows are clusters that anchor no macrostate. Basal clusters 7 and 8 each anchor two terminal states, whereas the lineage-low clusters 3 and 10 contribute almost no terminal states. **g.** Scaled activity score of *RPR2* archetype signatures across *ERPMT* tumour sampling times. AT2, alveolar type 2 cells.

**Supplementary Figure 9:**
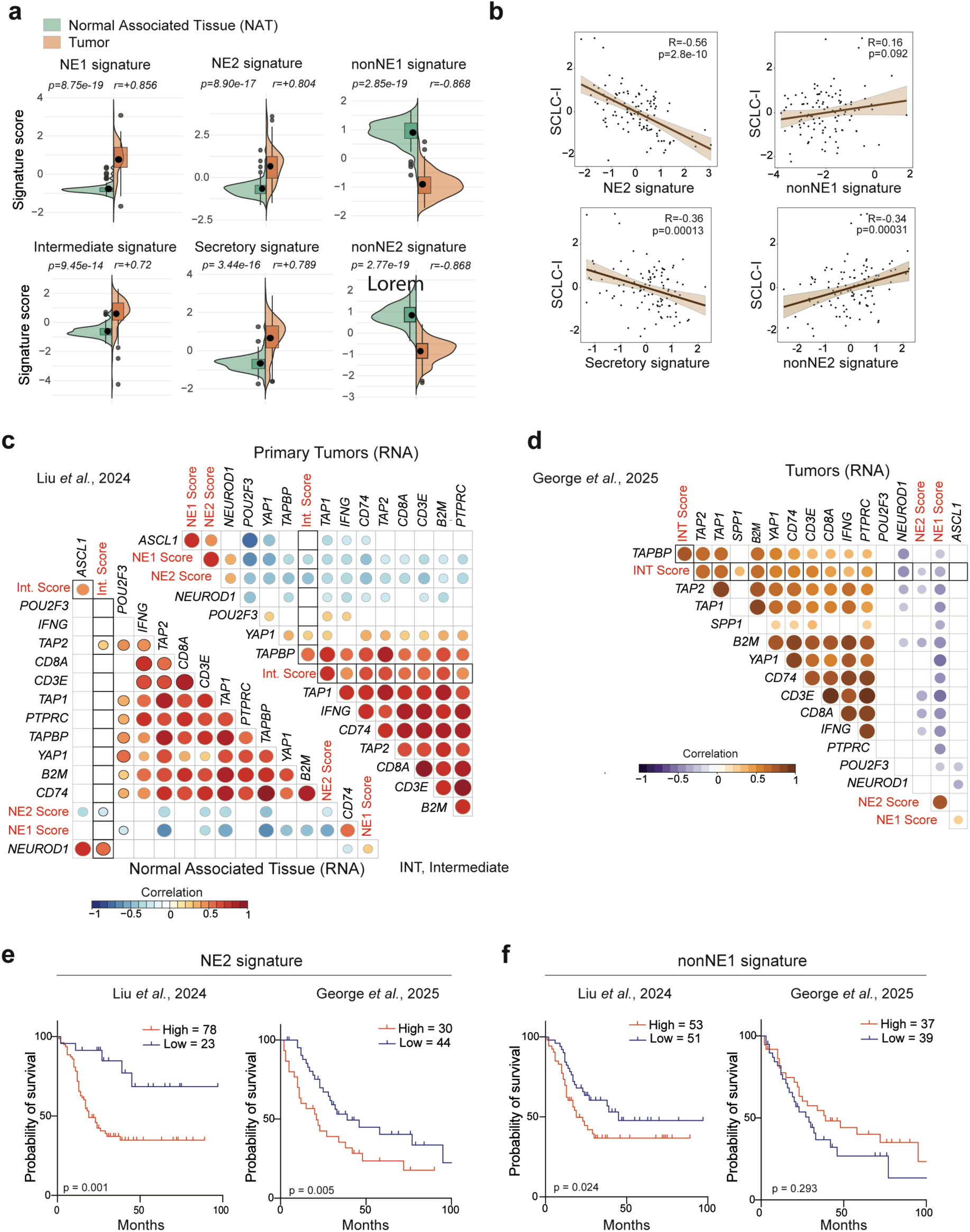
The Intermediate signature correlates with immune features in human SCLC. **a.** Plots comparing the gene signature activity of *RPR2* archetypes between tumour and normal adjacent tissue in the TU SCLC cohort from Liu *et al*. **b.** Plots showing the Pearson correlation between the human inflamed SCLC signature and the *RPR2* archetype signatures: NE2 signature (top left), nonNE1 (top right), Secretory (bottom left), and nonNE2 (bottom right) in individual primary tumour samples from the TU SCLC cohort. **c.** Correlation matrices showing associations between Intermediate, NE1, and NE2 signatures and RNA levels of antigen presentation genes, immune cell markers, and T cell markers in primary tumours (top) versus NAT samples (bottom) from the TU-SCLC cohort. Significant pairwise Pearson correlations (p < 0.01) are highlighted. **d.** Correlation matrices showing associations between Intermediate, NE1, and NE2 signatures and RNA levels of antigen presentation genes, immune cell markers, and T cell markers in primary and metastatic tumours of extensive-stage SCLC patients from the George et *al.* cohort (2025). **e.** Kaplan-Meier curves for overall survival based on *RPR2* archetype NE2 signature activity in limited-stage patients from the TU SCLC cohort (Liu *et al.*, left) and in extensive-stage patients from the George *et al.* cohort (right). P-value was determined through log-rank testing. **f.** Kaplan-Meier curves for overall survival based on *RPR2* archetype nonNE1 signature activity in limited-stage patients from the TU SCLC cohort (Liu *et al.*, left) and in extensive-stage patients from the George *et al.* cohort (right). P-value was determined through log-rank testing.

**Supplementary Figure 10:**
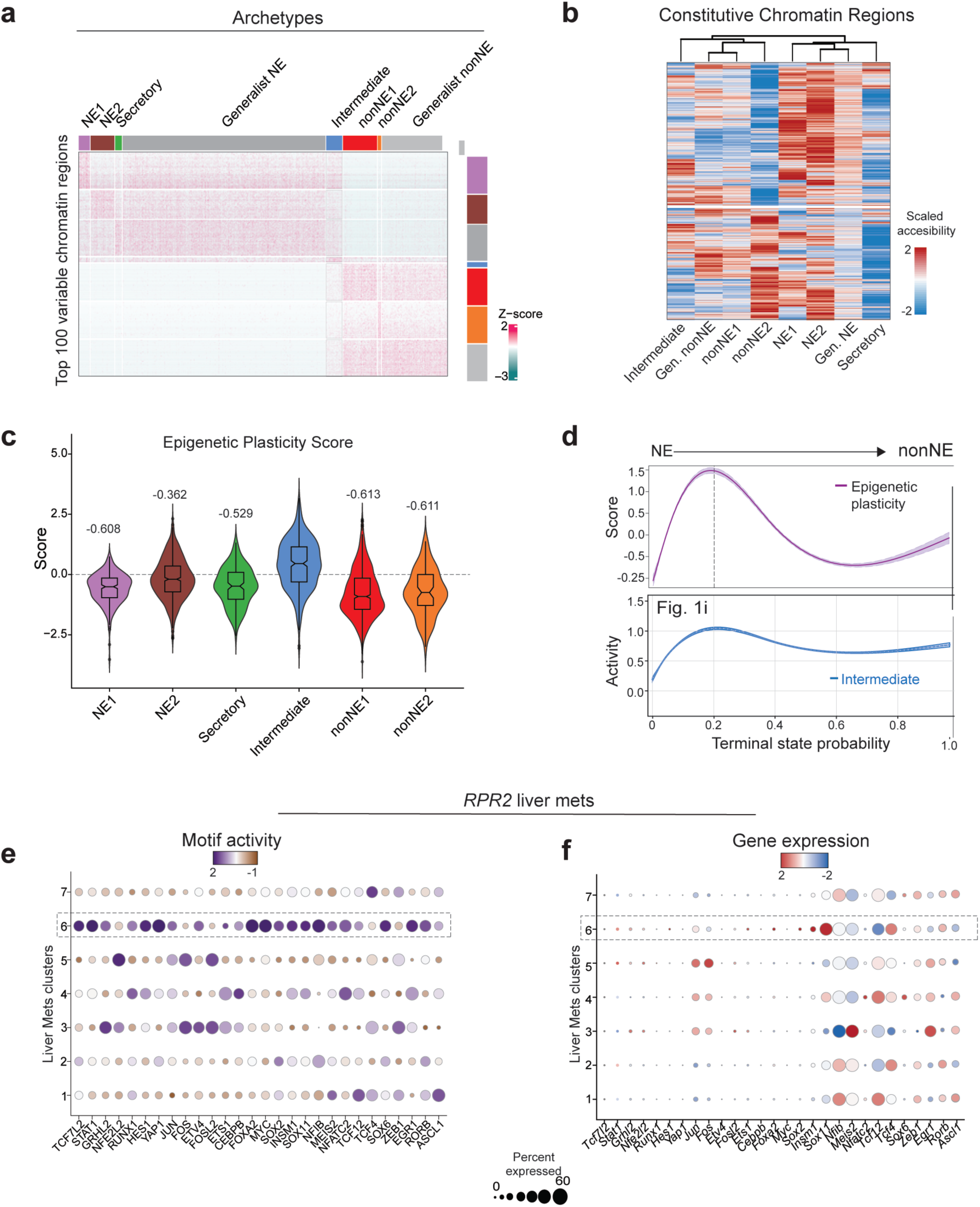
Epigenetic plasticity in Intermediate SCLC cells. **a.** Heatmap showing accessibility of top 100 variable chromatin regions of each archetype (y-axis) across cancer cells, individual cancer cells clustered based on their archetype identity (x-axis). **b.** Heatmap of scaled accessibility for constitutively open chromatin regions across *RPR2* cancer cell archetypes. **c.** Quantification of epigenetic plasticity score of cells in *RPR2* archetypes. The Cliff’s delta (δ) was used to assess the effect size between different distributions. Values of δ from 0 to 0.2 indicate a small effect, 0.2 to 0.5 denote a medium effect, and scores above 0.5 reflect a large difference. Each archetype was tested separately against the Intermediate state. A negative δ suggests lower values in the test archetype cells, whereas a positive score indicates higher values compared to the Intermediate state. **d.** Spline fit for epigenetic plasticity score (top) of *RPR2* cancer cells ordered by increasing terminal state probability. A similar plot for Intermediate signature activity (from Fig. 1j) is shown (bottom) to highlight that the two scores peak in the same cells. **e,f.** Dot plots comparing the motif activity (e) and RNA expression levels (f) of top variable transcription factors in liver metastases clusters.

**Supplementary Figure 11:**
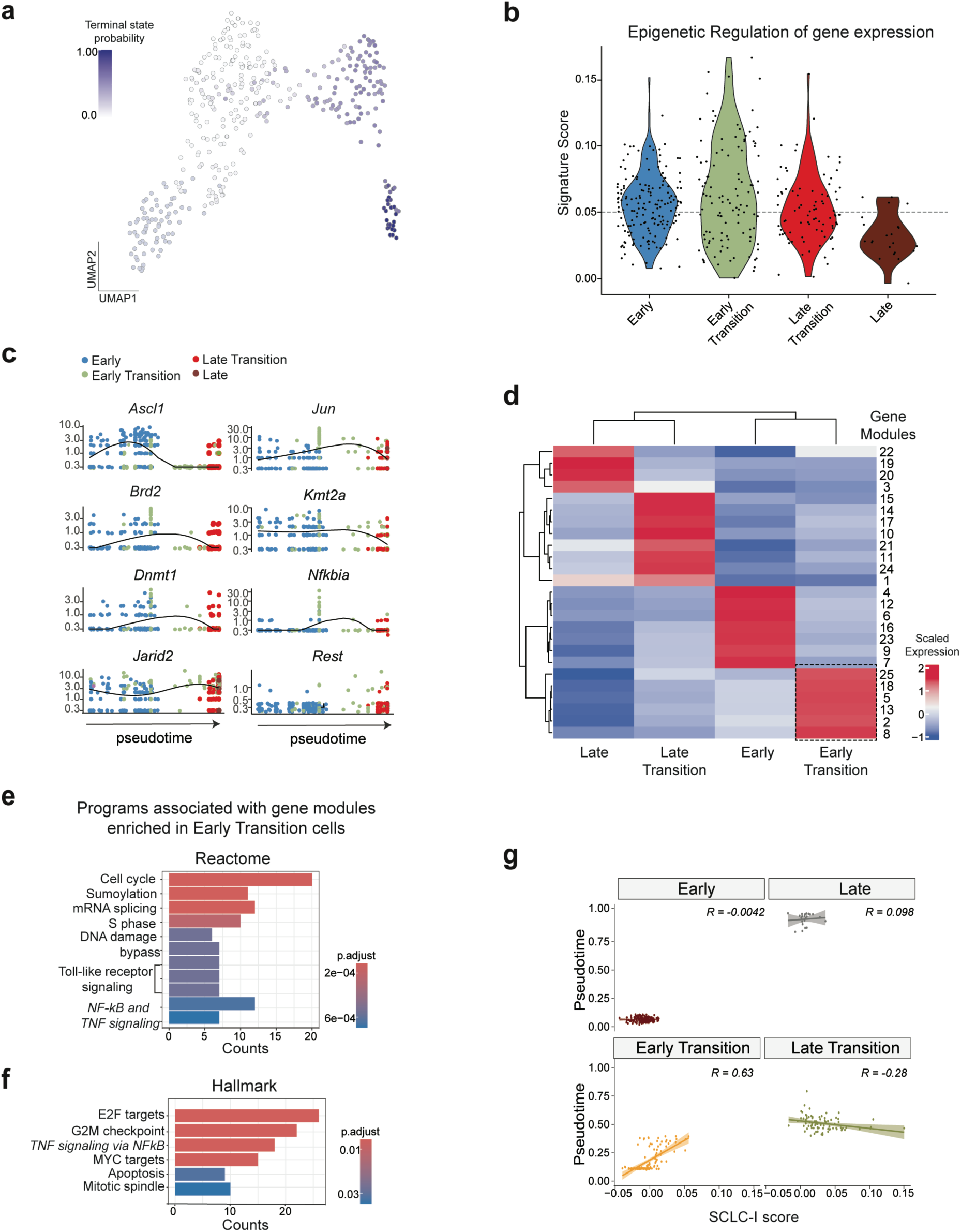
Epigenetic reprogramming and inflammation during Intermediate cell transitions. **a.** UMAP of Intermediate cancer cells showing terminal state probability scores of individual cells. **b.** Plots showing the expression of lineage marker genes (*Ascl1, Rest*), epigenetic modifiers (*Brd2, Dnmt1, Jarid2, Kmt2a*), and inflammation regulators (*Jun, Nfkbia*) in individual Intermediate cells ordered by their pseudotime score. **c.** Heatmap of gene modules that are differentially expressed between the four pseudotime clusters (early, early transitioning, late transitioning, and late) of cells in the Intermediate state, as determined by Monocle. The six gene modules upregulated in “early transitioning” cells are highlighted. **d,e.** Bar plots showing the Reactome (d) and Hallmark (e) pathways overrepresented in the six gene modules upregulated in early transitioning cancer cells. **f.** Violin plots comparing the activity of the epigenetic regulation signature in Intermediate cell clusters. **g.** Correlation plots between pseudotime progression and inflammation signature scores in Intermediate cells, stratified by pseudotime cluster identity.

**Supplementary Figure 12:**
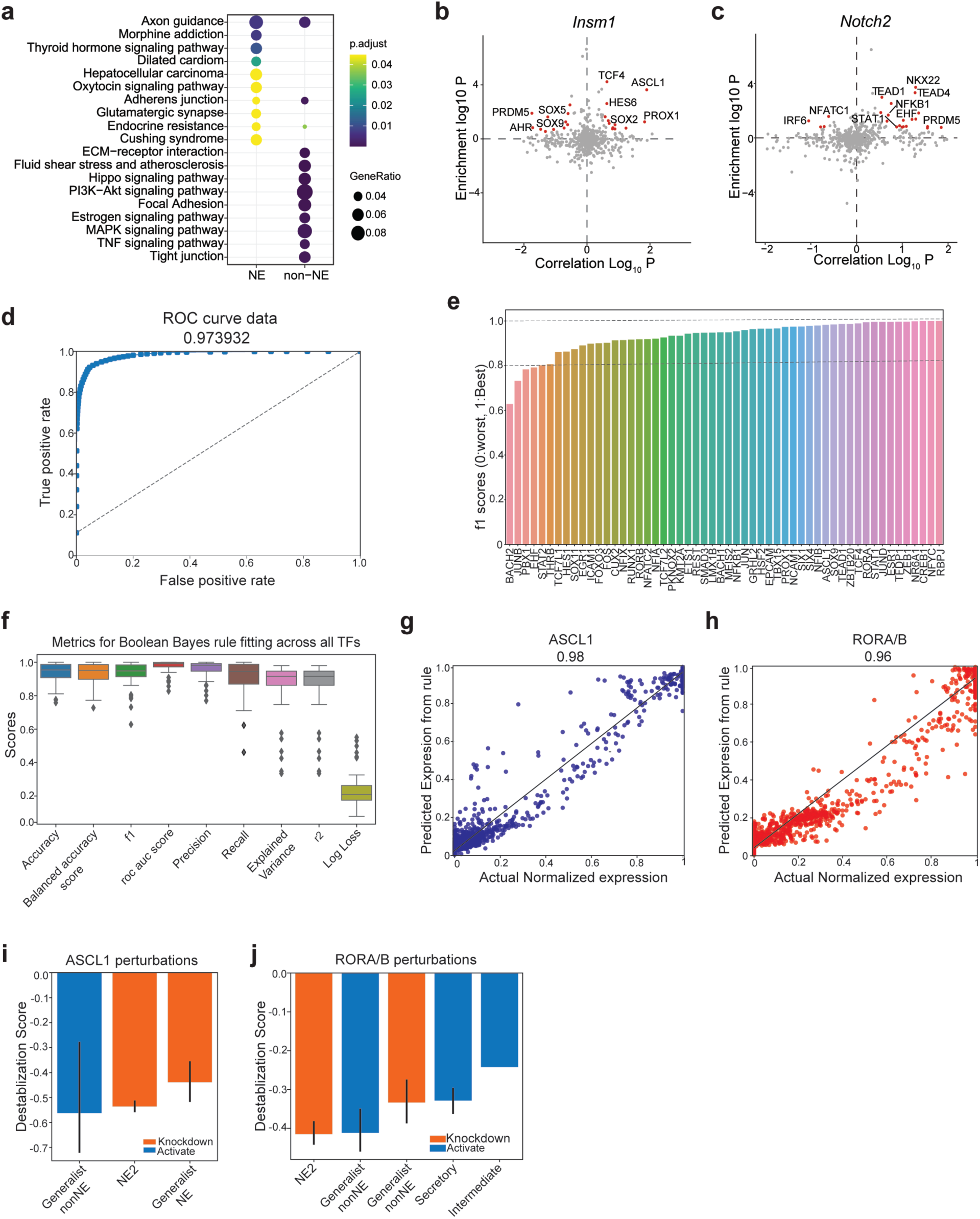
Identification of transcription factors controlling SCLC cell states. **a.** Plot comparing the significantly enriched KEGG (p < 0.05) pathways between NE and nonNE DORC genes shown in Fig. 5b. **b,c.** Candidate TF regulators for the NE lineage gene *Insm1* (b) and nonNE lineage gene *Notch2* (c); TFs with an absolute regulation score ≥ 1 are highlighted, while other TFs are shown in grey. **d.** A Receiver Operating Characteristic (ROC) plot showing classifier performance for BoBa-T by plotting its true positive rate versus false positive rate at different threshold settings. The area under the curve, a measure of overall performance, is shown at the top of the plot. **e.** f1 score, a measure of the model’s classification performance, for individual TFs in the network. A score between 0.7-0.9 indicates good performance, while a score above 0.9 indicates excellent performance of the model in predicting TF expression using Boolean-Bayes rules. **f.** Boxplot showing the distribution of model performance metric scores for all TFs. For all metrics, scores closer to 1 are ideal, except for log loss, which should be small for a well-performing model. **g,h.** Plots depicting the strong correlation between predicted expression (BoBa-T) and actual expression (scRNA-seq) of two transcription factors in the network, ASCL1 (g) and RORA/B (h). **i,j.** Barplots showing the extent of destabilisation of different *RPR2* archetypes upon simulated knockdown (orange) and overexpression (blue) of transcription factors ASCL1 (i) and RORA/B (j).

**Supplementary Figure 13:**
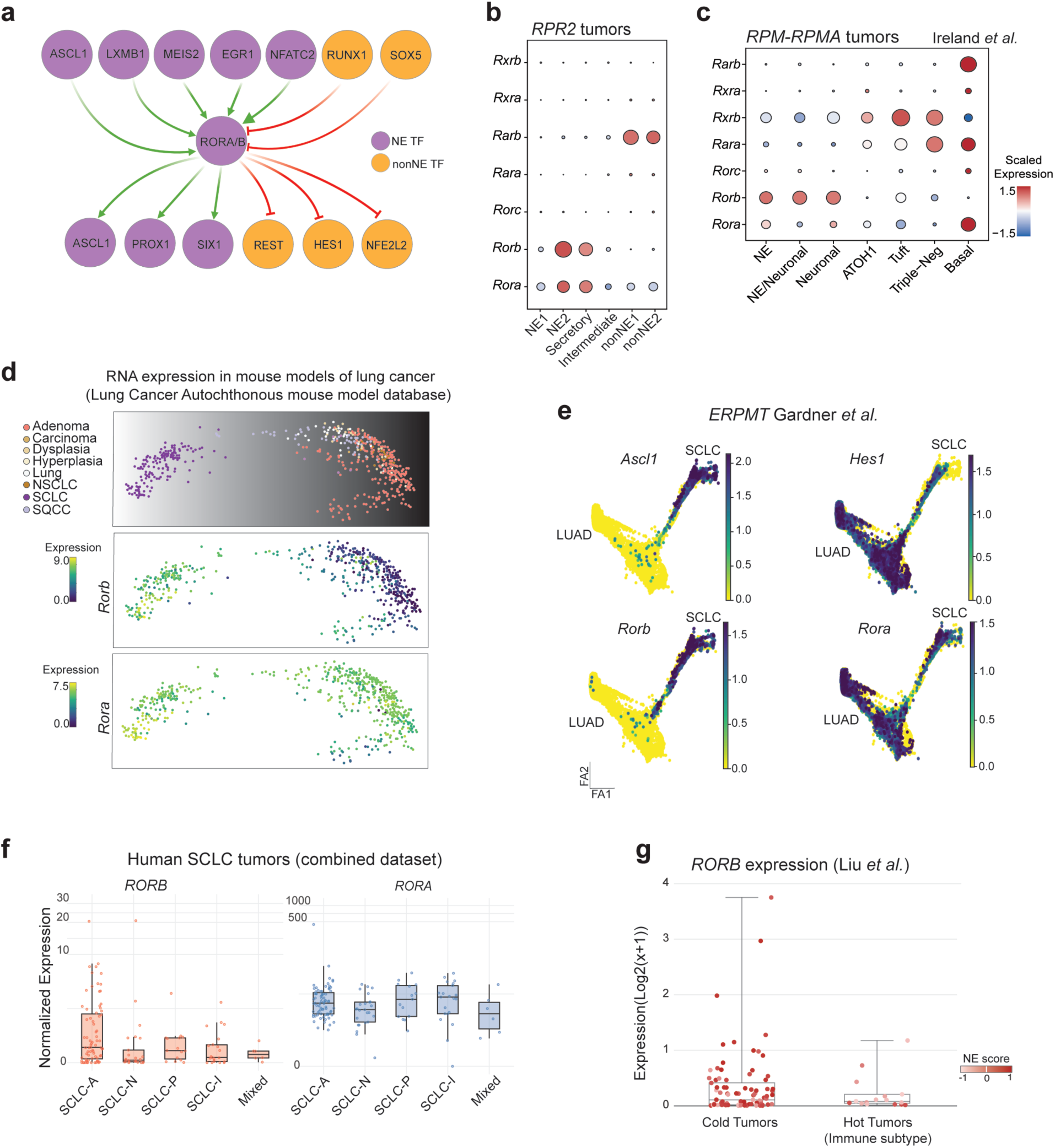
Network analysis identifies RORB as a transcription factor controlling SCLC cell states. **a.** Top upstream regulators and downstream targets of RORA/B transcription factors as predicted from scATAC-seq data and the BoBa-T algorithm. **b,c.** Dotplots showing the RNA expression of members of the retinoic acid-related nuclear receptor family across archetypes in *RPR2* scRNA-seq data (b) and cancer cell types in *RPM-RPMA* scRNA-seq data (c). **d.** PCA Plot of lung tumour transcriptomes from the Lung Cancer Autochthonous mouse model database (top) showing the expression of *Rorb* (middle) and *Rora* (bottom) RNA in published murine lung cancer models. **e.** Force-directed layout of tSCLC (*ERPMT* model) single-cell data depicting the transition from AT2 cells to LUAD and subsequently to neuroendocrine SCLC. NE markers *Ascl1* and *Rorb* are specifically expressed in SCLC cells, while nonNE markers *Hes1* and *Rora* are ubiquitously expressed through the LUAD to SCLC transition. FA, ForceAtlas components. **f.** *RORB* (and *RORA*) expression across transcriptional subtypes of SCLC in a combined patient bulk RNA-seq dataset from TU-SCLC, George *et al.,* and the Gemini cohort from Heeke *et al*. **g.** *RORB* expression in immune cold (low immune infiltrated) and immune hot (high immune infiltrated) tumours from TU-SCLC patients, as classified by Liu *et al.* Each dot represents an individual patient sample, coloured by the tumour’s NE score. P-value was determined by an unpaired non-parametric Wilcoxon test.

**Supplementary Figure 14:**
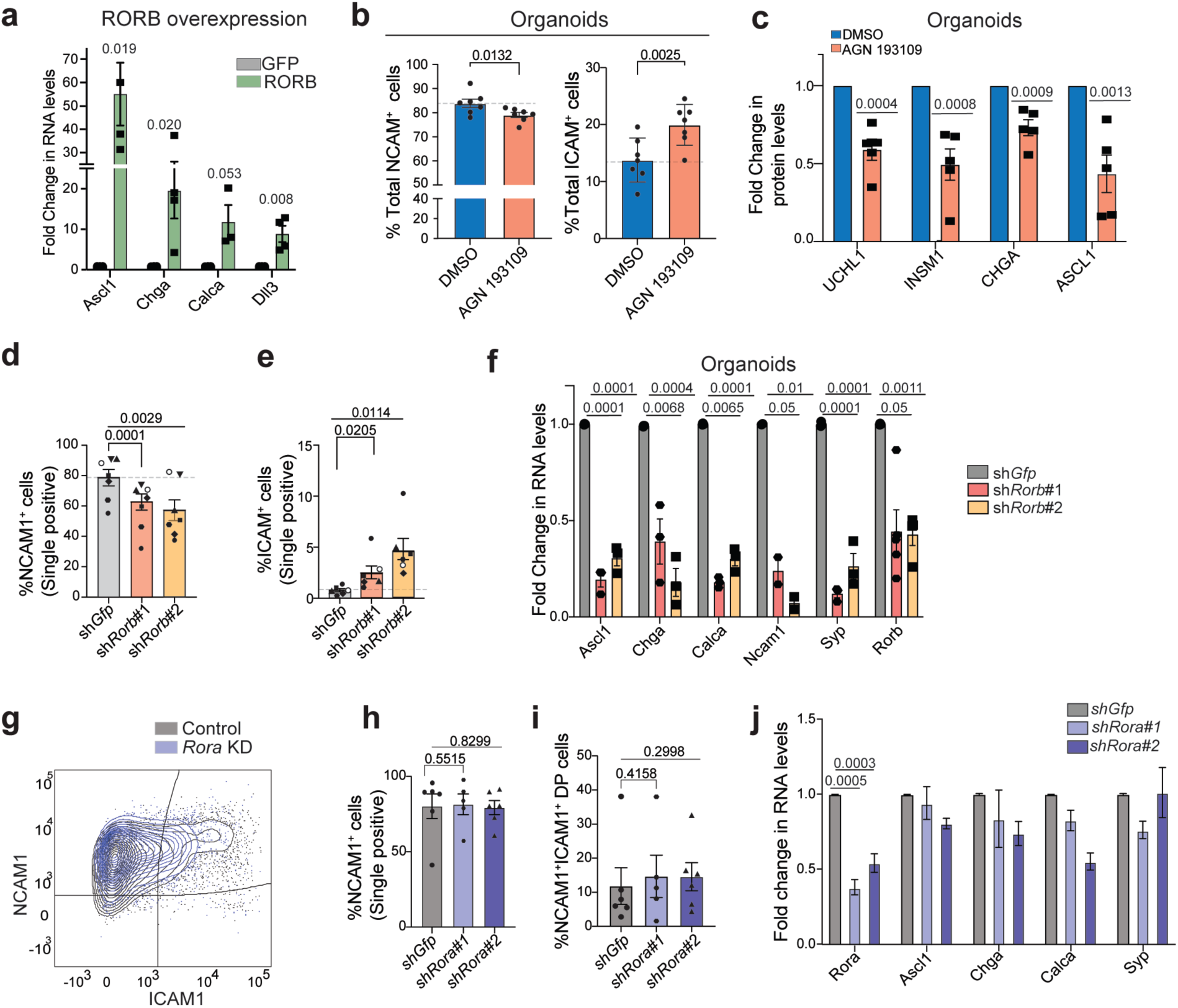
Pharmacological and genetic validation of RORB as a regulator of neuroendocrine identity. **a.** RT-qPCR analysis of neuroendocrine marker gene expression (*Ascl1, Chga, Calca, Dll3*) in Control and *Rorb* overexpressing organoids. n = 4 biological replicates. Error bars represent the s.e.m. P-values are determined by Wilcox test comparing control and overexpression samples. **b.** Flow cytometry-based quantitation of NCAM1 (left) and ICAM1 (right) surface expression in organoid lines treated with vehicle control (DMSO) or AGN 193109 (1 µM, 72 h). Representative plots from n = 7 biological replicates. Error bars represent the s.e.m. P-values are determined by paired t-tests comparing control and treated organoids. **c.** Quantitation of immunoassays for NE marker proteins (ASCL1, CHGA, SYP) in organoid lines treated with vehicle (DMSO) or AGN 193109 (1 µM, 72 h) (n = 7 biological replicates). Error bars represent the s.e.m. P-values were determined using the Wilcox test via pairwise comparisons between control and treated organoids. **d,e.** Quantification of NCAM1+ NE cells (d) and ICAM1^+^ nonNE cells (e) in organoid lines following *Rorb* knockdown (n = 7 biological replicates). Error bars represent the s.e.m. P-values were determined by paired t-tests comparing each *Rorb* shRNA to control *shGfp*. **f.** RT-qPCR analysis of NE marker gene expression (*Ascl1, Chga, Calca, Syp*) in *Rorb* knockdown (*shRorb#1, shRorb#2*) versus control (sh*Gfp*) organoid lines. Data are normalised to housekeeping genes and expressed as fold change relative to control (n = 6 biological replicates). Error bars represent the s.e.m. P-values were determined by the Wilcox test through pairwise comparison of individual shRNA lines to control sh*Gfp*. **g.** Representative overlaid flow cytometry plots showing NCAM1 and ICAM1 surface expression in control (grey, shGfp) and *Rora* knockdown (blue) organoid lines. **h,i.** Flow cytometry quantification of NCAM1^+^ NE cells (h), and NCAM1^+^ ICAM1^+^ Intermediate cells (i) in *Rora* knockdown (*shRora#1, shRora#2*) versus control (sh*Gfp*) organoid lines (n = 5 biological replicates). Error bars represent the s.e.m. P-values were determined by paired t-tests comparing each *Rora* shRNA to control sh*Gfp*. **j.** RT-qPCR analysis for *Rora* and NE marker genes (*Ascl1, Chga, Syp*) in *Rora* knockdown versus control organoid lines. Data are normalised to housekeeping genes and expressed as fold change. n = 3 biological replicates. Error bars represent s.e.m. P-values were determined by the Wilcox test through pairwise comparison of individual shRNA lines to control sh*Gfp*.

**Supplementary Figure 15:**
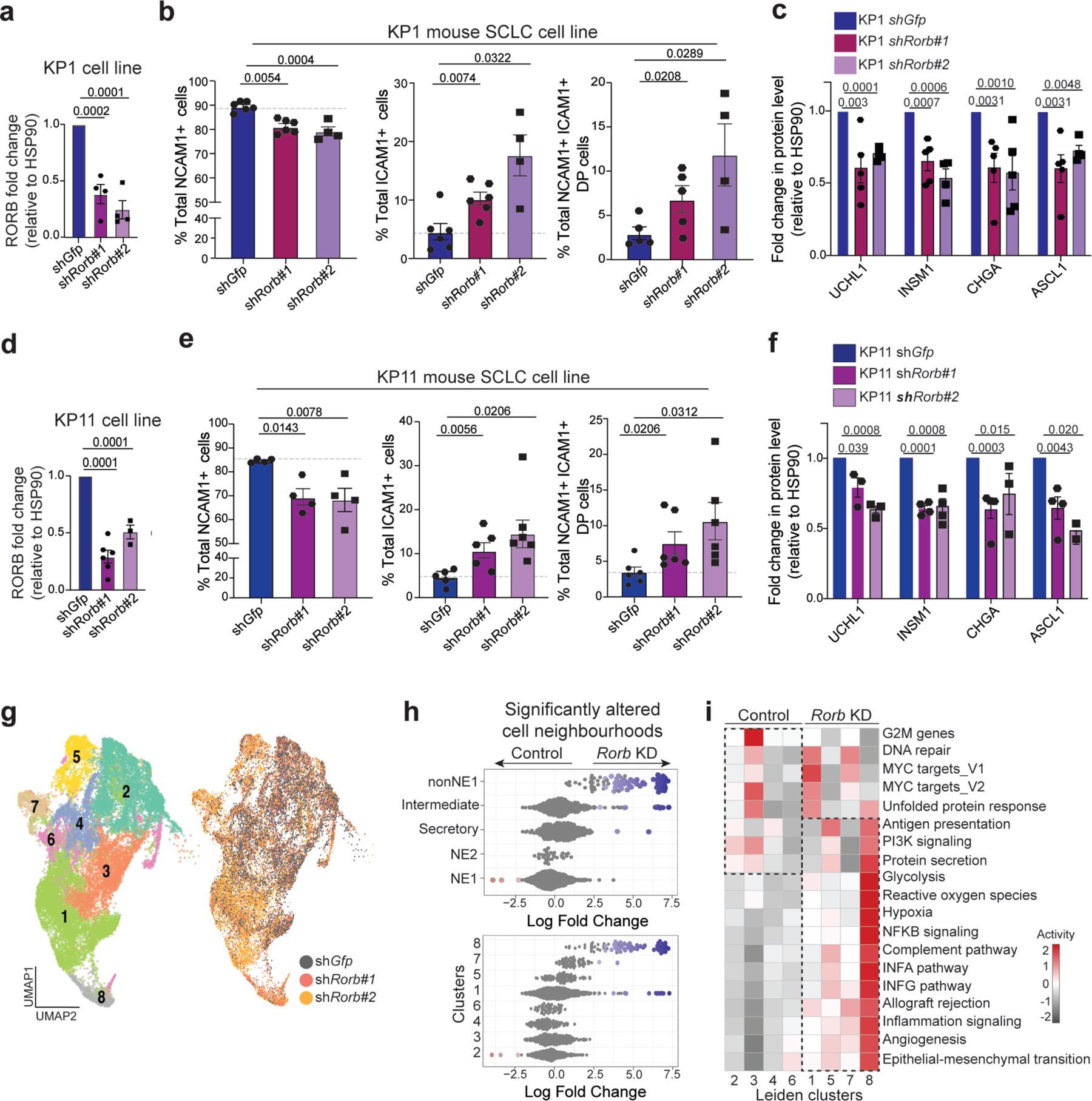
Validation of RORB as a regulator of neuroendocrine identity in murine SCLC cell lines. **a.** Quantification of immunoassays confirming *Rorb* knockdown in murine KP1 cells. Error bars represent the s.e.m. P-values were determined by the Wilcox test through pairwise comparison of individual shRNA lines to control *shGfp*. **b.** Plots showing the percentage of NCAM1^+^ NE cells, ICAM1^+^ nonNE cells, and NCAM1^+^ ICAM1^+^ double-positive (DP) Intermediate cells in control and *Rorb* knockdown KP1 cells. n = 4-5 technical replicates. Error bars represent the s.e.m. P-values were determined by the Wilcox test through pairwise comparison of individual shRNA lines to control *shGfp*. **c.** Quantification of immunoassays for NE marker proteins levels in KP1 control and *Rorb* knockdown cells. n = 4-5 technical replicates. Error bars represent the s.e.m. P-values were determined by the Wilcox test through pairwise comparison of individual shRNA lines to control *shGfp*. **d.** Immunoassay quantification confirming *Rorb* knockdown in murine KP11 cells. n = 5 technical replicates. Error bars represent the s.e.m. P-values were determined by the Wilcox test through pairwise comparison of individual shRNA lines to control *shGfp*. **e.** Bar plots showing the percentage of NCAM1^+^ NE cells, ICAM1^+^ nonNE cells, and NCAM1^+^ ICAM1^+^ double-positive (DP) Intermediate cells in control and *Rorb* knockdown KP11 cells. n = 4 technical replicates. Error bars represent the s.e.m. P-values were determined by the Wilcox test through pairwise comparison of individual shRNA lines to control *shGfp*. **f.** Quantification of immunoassays for NE marker proteins levels in KP11 control and *Rorb* knockdown cells. n = 4-5 technical replicates. Error bars represent the s.e.m. P-values were determined by the Wilcox test through pairwise comparison of individual shRNA lines to control *shGfp*. **g.** UMAP visualisation of scRNA-seq data from control (sh*Gfp*) and *Rorb* knockdown (*shRorb#1, shRorb#2*) tumour organoid lines (n = 5 biological replicates). The Leiden clustering identified eight cancer cell clusters, with cells coloured by their cluster assignments. **h.** Beeswarm plot showing the significantly altered cell neighbourhoods (FDR < 0.05) between Control and *Rorb* knockdown cells, based on their archetype identity (top) and cluster identity (bottom). **i.** Heatmap showing scaled expression of gene program signatures across the eight clusters. *Rorb* knockdown-enriched clusters (1, 5, 7, 8) exhibit upregulation of inflammation, interferon signalling, antigen presentation, and MYC target genes, with concurrent downregulation of neuroendocrine (NE) markers.

**Supplementary Figure 16:**
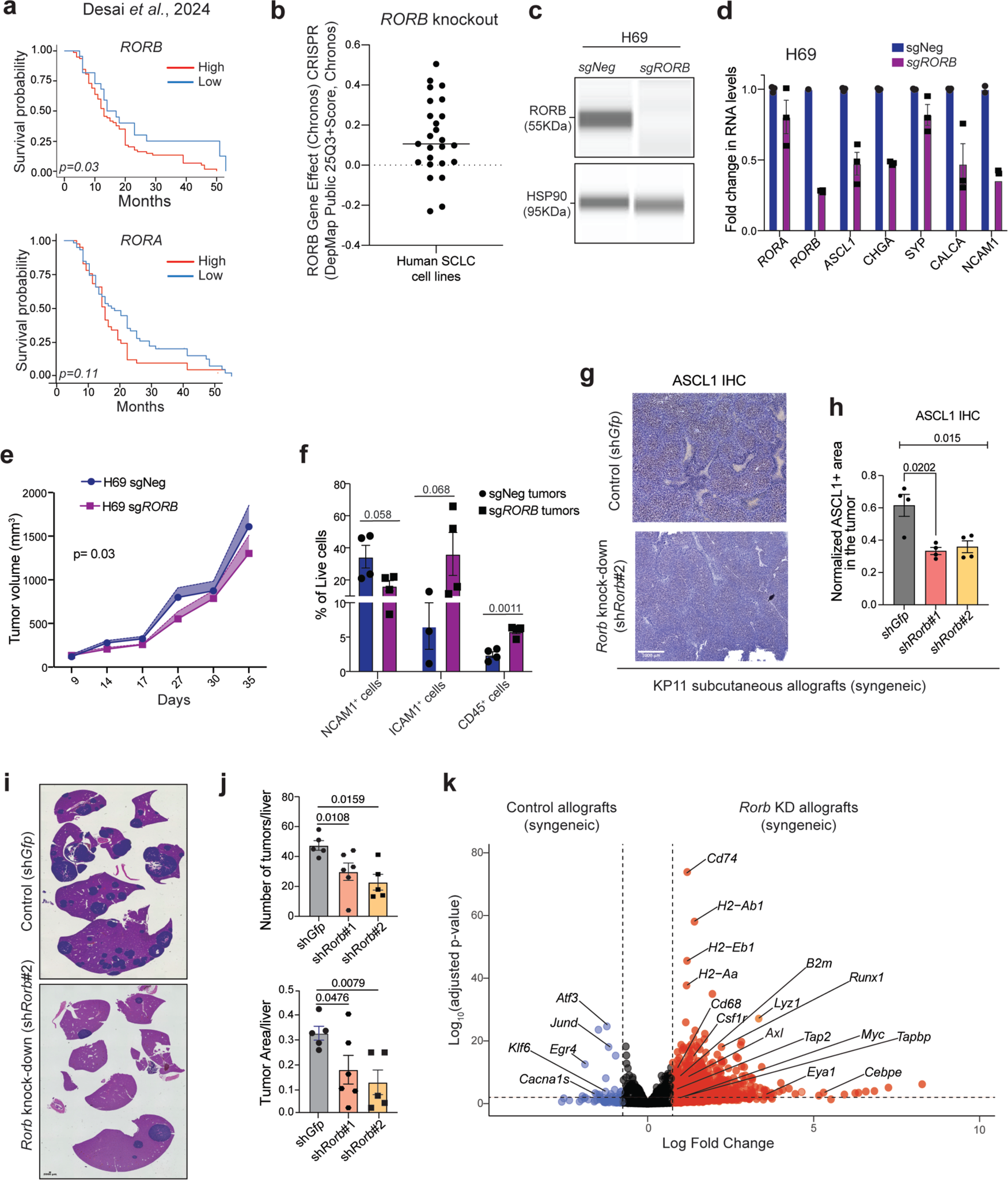
Single-cell transcriptomic characterisation of RORB loss and immune-mediated tumour suppression. **a.** Kaplan-Meier survival curves stratified by *RORB* (top) and *RORA* (bottom) expression levels in human SCLC patients from the Desai *et al.*, 2024 cohort. P-values were determined by log-rank test. **b.** Dot plot showing CRISPR knockout effect scores for *RORB* across human SCLC cell lines from the DepMap database. **c.** Representative immunoassays showing the expression of RORB and HSP90 in control (sg*Neg*) and knockout (sg*RORB*) human SCLC H69 cells. **d.** RT-qPCR analysis of RORB and other NE markers (*ASCL1, CHGA, SYP, CALCA*) in Control (sg*Neg*) and *RORB* knockout (sg*RORB*) H69 cells. Data are normalised to housekeeping genes and expressed as fold change. Mean ± s.e.m.; n = 3 technical replicates. **e.** Tumour growth kinetics of Control (sg*Neg*) and *RORB* knockout (sg*RORB*) human H69 cells in immunodeficient NSG mice. Data represent mean ± s.e.m.; n = 5-6 mice per group. P-values were calculated by a 2-way ANOVA test. **f.** Flow cytometry-based quantitation of NCAM1^+^ cancer cells, ICAM1^+^ cancer cells, and CD45^+^ immune cells as a percentage of total live cells in Control (sg*Neg*) and *RORB* knockout (sg*RORB*) human H69 xenograft tumours from (e). Data represent mean ± s.e.m.; n = 4 mice per group. P-values were calculated by Wilcox test. **g,h.** Representative immunohistochemistry(IHC) images (g) and quantification (h) of ASCL1 protein levels in control versus *Rorb* knockdown subcutaneous allografts grown in syngeneic mice. Data represent mean ± s.e.m.; n = 4 tumours per group. Scale bar, 1000 µm. **i.** Representative images of liver metastases (indicated by arrows) following tail vein injection of control (*shGfp*) or RORB knockdown (*shRorb#1, shRorb#2*) KP11 cells in syngeneic mice. **j.** Quantification of liver metastatic burden assessed by liver weight or metastatic nodule count in mice injected with control versus RORB knockdown KP11 cells. Data represent mean ± s.e.m.; n = 5-6 mice per group. P-values were determined by using the Wilcox test to perform pairwise comparison of *Rorb* knockdown lines with *shGfp* control. **k.** Volcano plot showing differentially expressed genes from bulk RNA-seq of control versus *Rorb* knockdown subcutaneous allografts grown in immunocompetent mice. The x-axis shows log2 fold change; y-axis shows −log10 adjusted p-value. Top upregulated genes (red) include MHC class I components (*H2-K1, H2-D1, B2m*), antigen presentation machinery (*Tap1, Tap2, Psmb9*), immune effector molecules (*Cxcl9, Cxcl10, Ccl5*), and nonNE differentiation markers (*Icam1, Cd274*).

## Notes

https://github.com/smgroves/Bhattacharya2026

## BIBLIOGRAPHY

1. Perez-Gonzalez A, Bevant K, Blanpain C. Cancer cell plasticity during tumor progression, metastasis and response to therapy. Nat Cancer. 2023;4(8):1063–82. Epub 20230803. doi: 10.1038/s43018-023-00595-y. PubMed PMID: 37537300; PMCID: PMC7615147.

2. Gargiulo G, Serresi M, Marine JC. Cell States in Cancer: Drivers, Passengers, and Trailers. Cancer Discov. 2024;14(4):610–4. doi: 10.1158/2159-8290.CD-23-1510. PubMed PMID: 38571419.

3. Mehta A, Stanger BZ. Lineage Plasticity: The New Cancer Hallmark on the Block. Cancer Res. 2024;84(2):184–91. doi: 10.1158/0008-5472.CAN-23-1067. PubMed PMID: 37963209.

4. Megyesfalvi Z, Gay CM, Popper H, Pirker R, Ostoros G, Heeke S, Lang C, Hoetzenecker K, Schwendenwein A, Boettiger K, Bunn PA, Jr., Renyi-Vamos F, Schelch K, Prosch H, Byers LA, Hirsch FR, Dome B. Clinical insights into small cell lung cancer: Tumor heterogeneity, diagnosis, therapy, and future directions. CA Cancer J Clin. 2023. Epub 20230617. doi: 10.3322/caac.21785. PubMed PMID: 37329269.

5. Simpson KL, Rothwell DG, Blackhall F, Dive C. Challenges of small cell lung cancer heterogeneity and phenotypic plasticity. Nat Rev Cancer. 2025. Epub 20250410. doi: 10.1038/s41568-025-00803-0. PubMed PMID: 40211072.

6. Sutherland KD, Ireland AS, Oliver TG. Killing SCLC: insights into how to target a shapeshifting tumor. Genes Dev. 2022;36(5-6):241–58. doi: 10.1101/gad.349359.122. PubMed PMID: 35318269; PMCID: PMC8973850.

7. Hanahan D. Hallmarks of Cancer: New Dimensions. Cancer Discov. 2022;12(1):31–46. doi: 10.1158/2159-8290.CD-21-1059. PubMed PMID: 35022204.

8. Gay CM, Stewart CA, Park EM, Diao L, Groves SM, Heeke S, Nabet BY, Fujimoto J, Solis LM, Lu W, Xi Y, Cardnell RJ, Wang Q, Fabbri G, Cargill KR, Vokes NI, Ramkumar K, Zhang B, Della Corte CM, Robson P, Swisher SG, Roth JA, Glisson BS, Shames DS, Wistuba, II, Wang J, Quaranta V, Minna J, Heymach JV, Byers LA. Patterns of transcription factor programs and immune pathway activation define four major subtypes of SCLC with distinct therapeutic vulnerabilities. Cancer Cell. 2021. Epub 2021/01/23. doi: 10.1016/j.ccell.2020.12.014. PubMed PMID: 33482121.

9. Pearsall SM, Humphrey S, Revill M, Morgan D, Frese KK, Galvin M, Kerr A, Carter M, Priest L, Blackhall F, Simpson KL, Dive C. The Rare YAP1 Subtype of SCLC Revisited in a Biobank of 39 Circulating Tumor Cell Patient Derived Explant Models: A Brief Report. J Thorac Oncol. 2020;15(12):1836–43. Epub 2020/07/30. doi: 10.1016/j.jtho.2020.07.008. PubMed PMID: 32721553; PMCID: PMC7718082.

10. Ito T, Matsubara D, Tanaka I, Makiya K, Tanei ZI, Kumagai Y, Shiu SJ, Nakaoka HJ, Ishikawa S, Isagawa T, Morikawa T, Shinozaki-Ushiku A, Goto Y, Nakano T, Tsuchiya T, Tsubochi H, Komura D, Aburatani H, Dobashi Y, Nakajima J, Endo S, Fukayama M, Sekido Y, Niki T, Murakami Y. Loss of YAP1 defines neuroendocrine differentiation of lung tumors. Cancer Sci. 2016;107(10):1527–38. Epub 2016/10/30. doi: 10.1111/cas.13013. PubMed PMID: 27418196; PMCID: PMC5084673.

11. Baine MK, Hsieh MS, Lai WV, Egger JV, Jungbluth AA, Daneshbod Y, Beras A, Spencer R, Lopardo J, Bodd F, Montecalvo J, Sauter JL, Chang JC, Buonocore DJ, Travis WD, Sen T, Poirier JT, Rudin CM, Rekhtman N. SCLC Subtypes Defined by ASCL1, NEUROD1, POU2F3, and YAP1: A Comprehensive Immunohistochemical and Histopathologic Characterization. J Thorac Oncol. 2020;15(12):1823–35. Epub 2020/10/05. doi: 10.1016/j.jtho.2020.09.009. PubMed PMID: 33011388.

12. Ireland AS, Xie DA, Hawgood SB, Barbier MW, Zuo LY, Hanna BE, Lucas-Randolph S, Tyson DR, Witt BL, Govindan R, Dowlati A, Moser JC, Thomas A, Puri S, Rudin CM, Chan JM, Elliott A, Oliver TG. Basal cell of origin resolves neuroendocrine-tuft lineage plasticity in cancer. Nature. 2025. Epub 20250917. doi: 10.1038/s41586-025-09503-z. PubMed PMID: 40963028.

13. Chen H, Deng C, Gao J, Wang J, Fu F, Wang Y, Wang Q, Zhang M, Zhang S, Fan F, Liu K, Yang B, He Q, Zheng Q, Shen X, Wang J, Hu T, Zhu C, Yang F, He Y, Hu H, Wang J, Li Y, Zhang Y, Cao Z. Integrative spatial analysis reveals tumor heterogeneity and immune colony niche related to clinical outcomes in small cell lung cancer. Cancer Cell. 2025. Epub 20250214. doi: 10.1016/j.ccell.2025.01.012. PubMed PMID: 39983726.

14. Jin Y, Wu Y, Reuben A, Zhu L, Gay CM, Wu Q, Zhou X, Mo H, Zheng Q, Ren J, Fang Z, Peng T, Wang N, Ma L, Lungevity PIC, Fan Y, Song H, Zhang J, Chen M. Single-cell and spatial proteo-transcriptomic profiling reveals immune infiltration heterogeneity associated with neuroendocrine features in small cell lung cancer. Cell Discov. 2024;10(1):93. Epub 20240904. doi: 10.1038/s41421-024-00703-x. PubMed PMID: 39231924; PMCID: PMC11375181.

15. Zhang X, Wang H, Liu W, Xiao Z, Ma Z, Zhang Z, Gong W, Chen J, Liu Z. Molecular features and evolutionary trajectory of ASCL1(+) and NEUROD1(+) SCLC cells. Br J Cancer. 2023;128(5):748–59. Epub 20221214. doi: 10.1038/s41416-022-02103-y. PubMed PMID: 36517551; PMCID: PMC9977910.

16. Zhang Z, Sun X, Liu Y, Zhang Y, Yang Z, Dong J, Wang N, Ying J, Zhou M, Yang L. Spatial Transcriptome-Wide Profiling of Small Cell Lung Cancer Reveals Intra-Tumoral Molecular and Subtype Heterogeneity. Adv Sci (Weinh). 2024;11(31):e2402716. Epub 20240619. doi: 10.1002/adversus202402716. PubMed PMID: 38896789; PMCID: PMC11336901.

17. Tian Y, Li Q, Yang Z, Zhang S, Xu J, Wang Z, Bai H, Duan J, Zheng B, Li W, Cui Y, Wang X, Wan R, Fei K, Zhong J, Gao S, He J, Gay CM, Zhang J, Wang J, Tang F. Single-cell transcriptomic profiling reveals the tumor heterogeneity of small-cell lung cancer. Signal Transduct Target Ther. 2022;7(1):346. Epub 20221005. doi: 10.1038/s41392-022-01150-4. PubMed PMID: 36195615; PMCID: PMC9532437.

18. Chan JM, Quintanal-Villalonga A, Gao VR, Xie Y, Allaj V, Chaudhary O, Masilionis I, Egger J, Chow A, Walle T, Mattar M, Yarlagadda DVK, Wang JL, Uddin F, Offin M, Ciampricotti M, Qeriqi B, Bahr A, de Stanchina E, Bhanot UK, Lai WV, Bott MJ, Jones DR, Ruiz A, Baine MK, Li Y, Rekhtman N, Poirier JT, Nawy T, Sen T, Mazutis L, Hollmann TJ, Pe’er D, Rudin CM. Signatures of plasticity, metastasis, and immunosuppression in an atlas of human small cell lung cancer. Cancer Cell. 2021;39(11):1479–96 e18. Epub 2021/10/16. doi: 10.1016/j.ccell.2021.09.008. PubMed PMID: 34653364; PMCID: PMC8628860.

19. Liu Q, Zhang J, Guo C, Wang M, Wang C, Yan Y, Sun L, Wang D, Zhang L, Yu H, Hou L, Wu C, Zhu Y, Jiang G, Zhu H, Zhou Y, Fang S, Zhang T, Hu L, Li J, Liu Y, Zhang H, Zhang B, Ding L, Robles AI, Rodriguez H, Gao D, Ji H, Zhou H, Zhang P. Proteogenomic characterization of small cell lung cancer identifies biological insights and subtype-specific therapeutic strategies. Cell. 2024;187(1):184–203 e28. doi: 10.1016/j.cell.2023.12.004. PubMed PMID: 38181741.

20. Sriuranpong V, Borges MW, Ravi RK, Arnold DR, Nelkin BD, Baylin SB, Ball DW. Notch signaling induces cell cycle arrest in small cell lung cancer cells. Cancer Res. 2001;61(7):3200–5. Epub 2001/04/18. PubMed PMID: 11306509.

21. Coulson JM, Edgson JL, Woll PJ, Quinn JP. A splice variant of the neuron-restrictive silencer factor repressor is expressed in small cell lung cancer: a potential role in derepression of neuroendocrine genes and a useful clinical marker. Cancer Res. 2000;60(7):1840–4. Epub 2000/04/15. PubMed PMID: 10766169.

22. Lim JS, Ibaseta A, Fischer MM, Cancilla B, O’Young G, Cristea S, Luca VC, Yang D, Jahchan NS, Hamard C, Antoine M, Wislez M, Kong C, Cain J, Liu YW, Kapoun AM, Garcia KC, Hoey T, Murriel CL, Sage J. Intratumoural heterogeneity generated by Notch signalling promotes small-cell lung cancer. Nature. 2017;545(7654):360–4. doi: 10.1038/nature22323. PubMed PMID: 28489825.

23. Shue YT, Drainas AP, Li NY, Pearsall SM, Morgan D, Sinnott-Armstrong N, Hipkins SQ, Coles GL, Lim JS, Oro AE, Simpson KL, Dive C, Sage J. A conserved YAP/Notch/REST network controls the neuroendocrine cell fate in the lungs. Nat Commun. 2022;13(1):2690. Epub 2022/05/17. doi: 10.1038/s41467-022-30416-2. PubMed PMID: 35577801; PMCID: PMC9110333.

24. Ireland AS, Micinski AM, Kastner DW, Guo B, Wait SJ, Spainhower KB, Conley CC, Chen OS, Guthrie MR, Soltero D, Qiao Y, Huang X, Tarapcsak S, Devarakonda S, Chalishazar MD, Gertz J, Moser JC, Marth G, Puri S, Witt BL, Spike BT, Oliver TG. MYC Drives Temporal Evolution of Small Cell Lung Cancer Subtypes by Reprogramming Neuroendocrine Fate. Cancer Cell. 2020;38(1):60–78 e12. doi: 10.1016/j.ccell.2020.05.001. PubMed PMID: 32473656.

25. Sun NY, Kumar S, Kim YS, Varghese D, Mendoza A, Nguyen R, Okada R, Reilly K, Widemann B, Pommier Y, Elloumi F, Dhall A, Taniyama D, Patel M, Aber E, Contreras CF, Kaplan RN, Kiseljak-Vassiliades K, Wierman ME, Martinez D, Pogoriler J, Hamilton AK, Diskin SJ, Maris JM, Robey RW, Gottesman MM, Del Rivero J, Roper N. Identification of the Notch ligand DLK1 as an immunotherapeutic target and regulator of tumor cell plasticity and chemoresistance in adrenocortical carcinoma. Nat Commun. 2025;16(1):5511. Epub 20250701. doi: 10.1038/s41467-025-60649-w. PubMed PMID: 40595495; PMCID: PMC12216638.

26. Augert A, Eastwood E, Ibrahim AH, Wu N, Grunblatt E, Basom R, Liggitt D, Eaton KD, Martins R, Poirier JT, Rudin CM, Milletti F, Cheng WY, Mack F, MacPherson D. Targeting NOTCH activation in small cell lung cancer through LSD1 inhibition. Sci Signal. 2019;12(567). doi: 10.1126/scisignal.aau2922. PubMed PMID: 30723171.

27. Mahadevan NR, Knelson EH, Wolff JO, Vajdi A, Saigi M, Campisi M, Hong D, Thai TC, Piel B, Han S, Reinhold BB, Duke-Cohan JS, Poitras MJ, Taus LJ, Lizotte PH, Portell A, Quadros V, Santucci AD, Murayama T, Canadas I, Kitajima S, Akitsu A, Fridrikh M, Watanabe H, Reardon B, Gokhale PC, Paweletz CP, Awad MM, Van Allen EM, Lako A, Wang XT, Chen B, Hong F, Sholl LM, Tolstorukov MY, Pfaff K, Janne PA, Gjini E, Edwards R, Rodig S, Reinherz EL, Oser MG, Barbie DA. Intrinsic Immunogenicity of Small Cell Lung Carcinoma Revealed by Its Cellular Plasticity. Cancer Discov. 2021;11(8):1952–69. Epub 2021/03/13. doi: 10.1158/2159-8290.CD-20-0913. PubMed PMID: 33707236; PMCID: PMC8338750.

28. Mohammad HP, Smitheman KN, Kamat CD, Soong D, Federowicz KE, Van Aller GS, Schneck JL, Carson JD, Liu Y, Butticello M, Bonnette WG, Gorman SA, Degenhardt Y, Bai Y, McCabe MT, Pappalardi MB, Kasparec J, Tian X, McNulty KC, Rouse M, McDevitt P, Ho T, Crouthamel M, Hart TK, Concha NO, McHugh CF, Miller WH, Dhanak D, Tummino PJ, Carpenter CL, Johnson NW, Hann CL, Kruger RG. A DNA Hypomethylation Signature Predicts Antitumor Activity of LSD1 Inhibitors in SCLC. Cancer Cell. 2015;28(1):57–69. Epub 2015/07/16. doi: 10.1016/j.ccell.2015.06.002. PubMed PMID: 26175415.

29. George J, Lim JS, Jang SJ, Cun Y, Ozretic L, Kong G, Leenders F, Lu X, Fernandez-Cuesta L, Bosco G, Muller C, Dahmen I, Jahchan NS, Park KS, Yang D, Karnezis AN, Vaka D, Torres A, Wang MS, Korbel JO, Menon R, Chun SM, Kim D, Wilkerson M, Hayes N, Engelmann D, Putzer B, Bos M, Michels S, Vlasic I, Seidel D, Pinther B, Schaub P, Becker C, Altmuller J, Yokota J, Kohno T, Iwakawa R, Tsuta K, Noguchi M, Muley T, Hoffmann H, Schnabel PA, Petersen I, Chen Y, Soltermann A, Tischler V, Choi CM, Kim YH, Massion PP, Zou Y, Jovanovic D, Kontic M, Wright GM, Russell PA, Solomon B, Koch I, Lindner M, Muscarella LA, la Torre A, Field JK, Jakopovic M, Knezevic J, Castanos-Velez E, Roz L, Pastorino U, Brustugun OT, Lund-Iversen M, Thunnissen E, Kohler J, Schuler M, Botling J, Sandelin M, Sanchez-Cespedes M, Salvesen HB, Achter V, Lang U, Bogus M, Schneider PM, Zander T, Ansen S, Hallek M, Wolf J, Vingron M, Yatabe Y, Travis WD, Nurnberg P, Reinhardt C, Perner S, Heukamp L, Buttner R, Haas SA, Brambilla E, Peifer M, Sage J, Thomas RK. Comprehensive genomic profiles of small cell lung cancer. Nature. 2015;524(7563):47–53. doi: 10.1038/nature14664. PubMed PMID: 26168399.

30. Denny SK, Yang D, Chuang CH, Brady JJ, Lim JS, Gruner BM, Chiou SH, Schep AN, Baral J, Hamard C, Antoine M, Wislez M, Kong CS, Connolly AJ, Park KS, Sage J, Greenleaf WJ, Winslow MM. Nfib Promotes Metastasis through a Widespread Increase in Chromatin Accessibility. Cell. 2016;166(2):328–42. doi: 10.1016/j.cell.2016.05.052. PubMed PMID: 27374332.

31. Ko JH, Lambert KE, Bhattacharya D, Lee MC, Colon CI, Hauser H, Sage J. Small Cell Lung Cancer Plasticity Enables NFIB-Independent Metastasis. Cancer Res. 2024;84(2):226–40. doi: 10.1158/0008-5472.CAN-23-1079. PubMed PMID: 37963187; PMCID: PMC10842891.

32. Shue YT, Lim JS, Sage J. Tumor heterogeneity in small cell lung cancer defined and investigated in pre-clinical mouse models. Transl Lung Cancer Res. 2018;7(1):21–31. doi: 10.21037/tlcr.2018.01.15. PubMed PMID: 29535910; PMCID: PMC5835592.

33. Shoval O, Sheftel H, Shinar G, Hart Y, Ramote O, Mayo A, Dekel E, Kavanagh K, Alon U. Evolutionary trade-offs, Pareto optimality, and the geometry of phenotype space. Science. 2012;336(6085):1157–60. Epub 20120426. doi: 10.1126/science.1217405. PubMed PMID: 22539553.

34. Hausser J, Szekely P, Bar N, Zimmer A, Sheftel H, Caldas C, Alon U. Tumor diversity and the trade-off between universal cancer tasks. Nat Commun. 2019;10(1):5423. Epub 20191128. doi: 10.1038/s41467-019-13195-1. PubMed PMID: 31780652; PMCID: PMC6882839.

35. Chan JM, Zaidi S, Love JR, Zhao JL, Setty M, Wadosky KM, Gopalan A, Choo ZN, Persad S, Choi J, LaClair J, Lawrence KE, Chaudhary O, Xu T, Masilionis I, Linkov I, Wang S, Lee C, Barlas A, Morris MJ, Mazutis L, Chaligne R, Chen Y, Goodrich DW, Karthaus WR, Pe’er D, Sawyers CL. Lineage plasticity in prostate cancer depends on JAK/STAT inflammatory signaling. Science. 2022;377(6611):1180–91. Epub 20220818. doi: 10.1126/science.abn0478. PubMed PMID: 35981096; PMCID: PMC9653178.

36. Gardner EE, Earlie EM, Li K, Thomas J, Hubisz MJ, Stein BD, Zhang C, Cantley LC, Laughney AM, Varmus H. Lineage-specific intolerance to oncogenic drivers restricts histological transformation. Science. 2024;383(6683):eadj1415. Epub 20240209. doi: 10.1126/science.adj1415. PubMed PMID: 38330136.

37. Nabet BY, Hamidi H, Lee MC, Banchereau R, Morris S, Adler L, Gayevskiy V, Elhossiny AM, Srivastava MK, Patil NS, Smith KA, Jesudason R, Chan C, Chang PS, Fernandez M, Rost S, McGinnis LM, Koeppen H, Gay CM, Minna JD, Heymach JV, Chan JM, Rudin CM, Byers LA, Liu SV, Reck M, Shames DS. Immune heterogeneity in small-cell lung cancer and vulnerability to immune checkpoint blockade. Cancer Cell. 2024;42(3):429–43 e4. Epub 20240215. doi: 10.1016/j.ccell.2024.01.010. PubMed PMID: 38366589.

38. Groves SM, Ildefonso GV, McAtee CO, Ozawa PMM, Ireland AS, Stauffer PE, Wasdin PT, Huang X, Qiao Y, Lim JS, Bader J, Liu Q, Simmons AJ, Lau KS, Iams WT, Hardin DP, Saff EB, Holmes WR, Tyson DR, Lovly CM, Rathmell JC, Marth G, Sage J, Oliver TG, Weaver AM, Quaranta V. Archetype tasks link intratumoral heterogeneity to plasticity and cancer hallmarks in small cell lung cancer. Cell Syst. 2022;13(9):690–710 e17. Epub 20220817. doi: 10.1016/j.cels.2022.07.006. PubMed PMID: 35981544; PMCID: PMC9615940.

39. Lange M, Bergen V, Klein M, Setty M, Reuter B, Bakhti M, Lickert H, Ansari M, Schniering J, Schiller HB, Pe’er D, Theis FJ. CellRank for directed single-cell fate mapping. Nat Methods. 2022;19(2):159–70. Epub 20220113. doi: 10.1038/s41592-021-01346-6. PubMed PMID: 35027767; PMCID: PMC8828480.

40. Klein D, Palla G, Lange M, Klein M, Piran Z, Gander M, Meng-Papaxanthos L, Sterr M, Saber L, Jing C, Bastidas-Ponce A, Cota P, Tarquis-Medina M, Parikh S, Gold I, Lickert H, Bakhti M, Nitzan M, Cuturi M, Theis FJ. Mapping cells through time and space with moscot. Nature. 2025;638(8052):1065–75. Epub 20250122. doi: 10.1038/s41586-024-08453-2. PubMed PMID: 39843746; PMCID: PMC11864987.

41. Guo C, Kong W, Kamimoto K, Rivera-Gonzalez GC, Yang X, Kirita Y, Morris SA. CellTag Indexing: genetic barcode-based sample multiplexing for single-cell genomics. Genome Biol. 2019;20(1):90. Epub 20190509. doi: 10.1186/s13059-019-1699-y. PubMed PMID: 31072405; PMCID: PMC6509836.

42. Sen T, Takahashi N, Chakraborty S, Takebe N, Nassar AH, Karim NA, Puri S, Naqash AR. Emerging advances in defining the molecular and therapeutic landscape of small-cell lung cancer. Nature reviews Clinical oncology. 2024;21(8):610–27. Epub 20240704. doi: 10.1038/s41571-024-00914-x. PubMed PMID: 38965396; PMCID: PMC11875021.

43. George J, Maas L, Abedpour N, Cartolano M, Kaiser L, Fischer RN, Scheel AH, Weber JP, Hellmich M, Bosco G, Volz C, Mueller C, Dahmen I, John F, Alves CP, Werr L, Panse JP, Kirschner M, Engel-Riedel W, Jurgens J, Stoelben E, Brockmann M, Grau S, Sebastian M, Stratmann JA, Kern J, Hummel HD, Hegedus B, Schuler M, Plones T, Aigner C, Elter T, Toepelt K, Ko YD, Kurz S, Grohe C, Serke M, Hopker K, Hagmeyer L, Doerr F, Hekmath K, Strapatsas J, Kambartel KO, Chakupurakal G, Busch A, Bauernfeind FG, Griesinger F, Luers A, Dirks W, Wiewrodt R, Luecke A, Rodermann E, Diel A, Hagen V, Severin K, Ullrich RT, Reinhardt HC, Quaas A, Bogus M, Courts C, Nurnberg P, Becker K, Achter V, Buttner R, Wolf J, Peifer M, Thomas RK. Evolutionary trajectories of small cell lung cancer under therapy. Nature. 2024;627(8005):880–9. Epub 20240313. doi: 10.1038/s41586-024-07177-7. PubMed PMID: 38480884; PMCID: PMC10972747.

44. Cao J, Spielmann M, Qiu X, Huang X, Ibrahim DM, Hill AJ, Zhang F, Mundlos S, Christiansen L, Steemers FJ, Trapnell C, Shendure J. The single-cell transcriptional landscape of mammalian organogenesis. Nature. 2019;566(7745):496–502. Epub 20190220. doi: 10.1038/s41586-019-0969-x. PubMed PMID: 30787437; PMCID: PMC6434952.

45. Davies A, Zoubeidi A, Beltran H, Selth LA. The Transcriptional and Epigenetic Landscape of Cancer Cell Lineage Plasticity. Cancer Discov. 2023;13(8):1771–88. doi: 10.1158/2159-8290.CD-23-0225. PubMed PMID: 37470668; PMCID: PMC10527883.

46. Kartha VK, Duarte FM, Hu Y, Ma S, Chew JG, Lareau CA, Earl A, Burkett ZD, Kohlway AS, Lebofsky R, Buenrostro JD. Functional inference of gene regulation using single-cell multiomics. Cell Genom. 2022;2(9). Epub 20220804. doi: 10.1016/j.xgen.2022.100166. PubMed PMID: 36204155; PMCID: PMC9534481.

47. Ma S, Zhang B, LaFave LM, Earl AS, Chiang Z, Hu Y, Ding J, Brack A, Kartha VK, Tay T, Law T, Lareau C, Hsu YC, Regev A, Buenrostro JD. Chromatin Potential Identified by Shared Single-Cell Profiling of RNA and Chromatin. Cell. 2020;183(4):1103–16 e20. Epub 20201023. doi: 10.1016/j.cell.2020.09.056. PubMed PMID: 33098772; PMCID: PMC7669735.

48. Minnie SA, Waltner OG, Zhang P, Takahashi S, Nemychenkov NS, Ensbey KS, Schmidt CR, Legg SRW, Comstock M, Boiko JR, Nelson E, Bhise SS, Wilkens AB, Koyama M, Dhodapkar MV, Chesi M, Riddell SR, Green DJ, Spencer A, Furlan SN, Hill GR. TIM-3(+) CD8 T cells with a terminally exhausted phenotype retain functional capacity in hematological malignancies. Sci Immunol. 2024;9(94):eadg1094. Epub 20240419. doi: 10.1126/sciimmunol.adg1094. PubMed PMID: 38640253; PMCID: PMC11093588.

49. Moorman A, Benitez EK, Cambulli F, Jiang Q, Mahmoud A, Lumish M, Hartner S, Balkaran S, Bermeo J, Asawa S, Firat C, Saxena A, Wu F, Luthra A, Burdziak C, Xie Y, Sgambati V, Luckett K, Li Y, Yi Z, Masilionis I, Soares K, Pappou E, Yaeger R, Kingham TP, Jarnagin W, Paty PB, Weiser MR, Mazutis L, D’Angelica M, Shia J, Garcia-Aguilar J, Nawy T, Hollmann TJ, Chaligne R, Sanchez-Vega F, Sharma R, Pe’er D, Ganesh K. Progressive plasticity during colorectal cancer metastasis. Nature. 2025;637(8047):947–54. Epub 20241030. doi: 10.1038/s41586-024-08150-0. PubMed PMID: 39478232; PMCID: PMC11754107.

50. Krebs AM, Mitschke J, Lasierra Losada M, Schmalhofer O, Boerries M, Busch H, Boettcher M, Mougiakakos D, Reichardt W, Bronsert P, Brunton VG, Pilarsky C, Winkler TH, Brabletz S, Stemmler MP, Brabletz T. The EMT-activator Zeb1 is a key factor for cell plasticity and promotes metastasis in pancreatic cancer. Nat Cell Biol. 2017;19(5):518–29. Epub 20170417. doi: 10.1038/ncb3513. PubMed PMID: 28414315.

51. Tolomeo M, Cavalli A, Cascio A. STAT1 and Its Crucial Role in the Control of Viral Infections. Int J Mol Sci. 2022;23(8). Epub 20220407. doi: 10.3390/ijms23084095. PubMed PMID: 35456913; PMCID: PMC9028532.

52. Pearsall SM, Williamson SC, Humphrey S, Hughes E, Morgan D, Garcia Marques FJ, Awanis G, Carroll R, Burks L, Shue YT, Bermudez A, Frese KK, Galvin M, Carter M, Priest L, Kerr A, Zhou C, Oliver TG, Humphries JD, Humphries MJ, Blackhall F, Cannell IG, Pitteri SJ, Hannon GJ, Sage J, Dive C, Simpson KL. Lineage Plasticity in SCLC Generates Non-Neuroendocrine Cells Primed for Vasculogenic Mimicry. J Thorac Oncol. 2023;18(10):1362–85. Epub 20230716. doi: 10.1016/j.jtho.2023.07.012. PubMed PMID: 37455012; PMCID: PMC10561473.

53. Wooten DJ, Groves SM, Tyson DR, Liu Q, Lim JS, Albert R, Lopez CF, Sage J, Quaranta V. Systems-level network modeling of Small Cell Lung Cancer subtypes identifies master regulators and destabilizers. PLoS Comput Biol. 2019;15(10):e1007343. Epub 20191031. doi: 10.1371/journal.pcbi.1007343. PubMed PMID: 31671086; PMCID: PMC6860456.

54. Augustyn A, Borromeo M, Wang T, Fujimoto J, Shao C, Dospoy PD, Lee V, Tan C, Sullivan JP, Larsen JE, Girard L, Behrens C, Wistuba, II, Xie Y, Cobb MH, Gazdar AF, Johnson JE, Minna JD. ASCL1 is a lineage oncogene providing therapeutic targets for high-grade neuroendocrine lung cancers. Proc Natl Acad Sci U S A. 2014;111(41):14788–93. Epub 2014/10/01. doi: 10.1073/pnas.1410419111. PubMed PMID: 25267614; PMCID: 4205603.

55. Cai L, Wu F, Zhou Q, Gao Y, Yao B, DeBerardinis RJ, Acquaah-Mensah GK, Aidinis V, Beane JE, Biswal S, Chen T, Concepcion-Crisol CP, Gruner BM, Jia D, Jones RA, Kurie JM, Lee MG, Lindahl P, Lissanu Y, Lorz C, MacPherson D, Martinelli R, Mazur PK, Mazzilli SA, Mii S, Moll HP, Moorehead RA, Morrisey EE, Ng SR, Oser MG, Pandiri AR, Powell CA, Ramadori G, Santos M, Snyder EL, Sotillo R, Su KY, Taki T, Taparra K, Tran PT, Xia Y, van Veen JE, Winslow MM, Xiao G, Rudin CM, Oliver TG, Xie Y, Minna JD. The Lung Cancer Autochthonous Model Gene Expression Database Enables Cross-Study Comparisons of the Transcriptomic Landscapes Across Mouse Models. Cancer Res. 2025;85(10):1769–83. doi: 10.1158/0008-5472.CAN-24-1607. PubMed PMID: 40298430; PMCID: PMC12081188.

56. Agarwal C, Chandraratna RA, Johnson AT, Rorke EA, Eckert RL. AGN193109 is a highly effective antagonist of retinoid action in human ectocervical epithelial cells. J Biol Chem. 1996;271(21):12209–12. doi: 10.1074/jbc.271.21.12209. PubMed PMID: 8647816.

57. Dann E, Henderson NC, Teichmann SA, Morgan MD, Marioni JC. Differential abundance testing on single-cell data using k-nearest neighbor graphs. Nat Biotechnol. 2022;40(2):245–53. Epub 20210930. doi: 10.1038/s41587-021-01033-z. PubMed PMID: 34594043; PMCID: PMC7617075.

58. Desai P, Takahashi N, Kumar R, Nichols S, Malin J, Hunt A, Schultz C, Cao Y, Tillo D, Nousome D, Chauhan L, Sciuto L, Jordan K, Rajapakse V, Tandon M, Lissa D, Zhang Y, Kumar S, Pongor L, Singh A, Schroder B, Sharma AK, Chang T, Vilimas R, Pinkiert D, Graham C, Butcher D, Warner A, Sebastian R, Mahon M, Baker K, Cheng J, Berger A, Lake R, Abel M, Krishnamurthy M, Chrisafis G, Fitzgerald P, Nirula M, Goyal S, Atkinson D, Bateman NW, Abulez T, Nair G, Apolo A, Guha U, Karim B, El Meskini R, Ohler ZW, Jolly MK, Schaffer A, Ruppin E, Kleiner D, Miettinen M, Brown GT, Hewitt S, Conrads T, Thomas A. Microenvironment shapes small-cell lung cancer neuroendocrine states and presents therapeutic opportunities. Cell Rep Med. 2024;5(6):101610. doi: 10.1016/j.xcrm.2024.101610. PubMed PMID: 38897168; PMCID: PMC11228806.

59. Li Y, Feng H, Gu H, Lewis DW, Yuan Y, Zhang L, Yu H, Zhang P, Cheng H, Miao W, Yuan W, Cheng SY, Gollin SM, Cheng T. The p53-PUMA axis suppresses iPSC generation. Nat Commun. 2013;4:2174. doi: 10.1038/ncomms3174. PubMed PMID: 23873265; PMCID: PMC4394110.

60. Raghavan S, Winter PS, Navia AW, Williams HL, DenAdel A, Lowder KE, Galvez-Reyes J, Kalekar RL, Mulugeta N, Kapner KS, Raghavan MS, Borah AA, Liu N, Vayrynen SA, Costa AD, Ng RWS, Wang J, Hill EK, Ragon DY, Brais LK, Jaeger AM, Spurr LF, Li YY, Cherniack AD, Booker MA, Cohen EF, Tolstorukov MY, Wakiro I, Rotem A, Johnson BE, McFarland JM, Sicinska ET, Jacks TE, Sullivan RJ, Shapiro GI, Clancy TE, Perez K, Rubinson DA, Ng K, Cleary JM, Crawford L, Manalis SR, Nowak JA, Wolpin BM, Hahn WC, Aguirre AJ, Shalek AK. Microenvironment drives cell state, plasticity, and drug response in pancreatic cancer. Cell. 2021;184(25):6119–37 e26. doi: 10.1016/j.cell.2021.11.017. PubMed PMID: 34890551; PMCID: PMC8822455.

61. Burdziak C, Alonso-Curbelo D, Walle T, Reyes J, Barriga FM, Haviv D, Xie Y, Zhao Z, Zhao CJ, Chen HA, Chaudhary O, Masilionis I, Choo ZN, Gao V, Luan W, Wuest A, Ho YJ, Wei Y, Quail DF, Koche R, Mazutis L, Chaligne R, Nawy T, Lowe SW, Pe’er D. Epigenetic plasticity cooperates with cell-cell interactions to direct pancreatic tumorigenesis. Science. 2023;380(6645):eadd5327. Epub 20230512. doi: 10.1126/science.add5327. PubMed PMID: 37167403; PMCID: PMC10316746.

62. Pitter KL, Grbovic-Huezo O, Joost S, Singhal A, Blum M, Wu K, Holm M, Ferrena A, Bhutkar A, Hudson A, Lecomte N, de Stanchina E, Chaligne R, Iacobuzio-Donahue CA, Pe’er D, Tammela T. Systematic Comparison of Pancreatic Ductal Adenocarcinoma Models Identifies a Conserved Highly Plastic Basal Cell State. Cancer Res. 2022;82(19):3549–60. doi: 10.1158/0008-5472.CAN-22-1742. PubMed PMID: 35952360; PMCID: PMC9532381.

63. Kaiser AM, Gatto A, Hanson KJ, Zhao RL, Raj N, Ozawa MG, Seoane JA, Bieging-Rolett KT, Wang M, Li I, Trope WL, Liou DZ, Shrager JB, Plevritis SK, Newman AM, Van Rechem C, Attardi LD. p53 governs an AT1 differentiation programme in lung cancer suppression. Nature. 2023;619(7971):851–9. Epub 20230719. doi: 10.1038/s41586-023-06253-8. PubMed PMID: 37468633; PMCID: PMC11288504.

64. Chan JE, Pan CH, Rub J, Guzman G, Krause K, Brown E, Zhang Z, Styers H, Hartmann G, Li Z, Zhuang X, Lowe SW, Betel D, Yan Y, Tammela T. Critical role for a high-plasticity cell state in lung cancer. Nature. 2026. Epub 20260121. doi: 10.1038/s41586-025-09985-x. PubMed PMID: 41565826.

65. Izzo LT, Reyes T, Meesala S, Ireland AS, Yang S, Sunil HS, Cheng XC, Tserentsoodol N, Hawgood SB, Patz EF, Jr., Witt BL, Tyson DR, O’Donnell KA, Oliver TG. KLF4 promotes a KRT13+ hillock-like state in squamous lung cancer. bioRxiv. 2025. Epub 20250313. doi: 10.1101/2025.03.10.641898. PubMed PMID: 40161723; PMCID: PMC11952405.

66. Karras P, Bordeu I, Pozniak J, Nowosad A, Pazzi C, Van Raemdonck N, Landeloos E, Van Herck Y, Pedri D, Bervoets G, Makhzami S, Khoo JH, Pavie B, Lamote J, Marin-Bejar O, Dewaele M, Liang H, Zhang X, Hua Y, Wouters J, Browaeys R, Bergers G, Saeys Y, Bosisio F, van den Oord J, Lambrechts D, Rustgi AK, Bechter O, Blanpain C, Simons BD, Rambow F, Marine JC. A cellular hierarchy in melanoma uncouples growth and metastasis. Nature. 2022;610(7930):190–8. Epub 20220921. doi: 10.1038/s41586-022-05242-7. PubMed PMID: 36131018; PMCID: PMC10439739.

67. Theunis K, Vanuytven S, Claes I, Geurts J, Rambow F, Brown D, Van Der Haegen M, Marin-Bejar O, Rogiers A, Van Raemdonck N, Leucci E, Demeulemeester J, Sifrim A, Marine JC, Voet T. Single-cell genome and transcriptome sequencing without upfront whole-genome amplification reveals cell state plasticity of melanoma subclones. Nucleic Acids Res. 2025;53(6). doi: 10.1093/nar/gkaf173. PubMed PMID: 40138718; PMCID: PMC11941470.

68. Sumi Y, Yagita K, Yamaguchi S, Ishida Y, Kuroda Y, Okamura H. Rhythmic expression of ROR beta mRNA in the mice suprachiasmatic nucleus. Neurosci Lett. 2002;320(1-2):13–6. doi: 10.1016/s0304-3940(02)00011-3. PubMed PMID: 11849752.

69. Takeda Y, Jothi R, Birault V, Jetten AM. RORgamma directly regulates the circadian expression of clock genes and downstream targets in vivo. Nucleic Acids Res. 2012;40(17):8519–35. Epub 20120629. doi: 10.1093/nar/gks630. PubMed PMID: 22753030; PMCID: PMC3458568.

70. Zhao X, Cho H, Yu RT, Atkins AR, Downes M, Evans RM. Nuclear receptors rock around the clock. EMBO Rep. 2014;15(5):518–28. Epub 20140415. doi: 10.1002/embr.201338271. PubMed PMID: 24737872; PMCID: PMC4210094.

71. Qin K, Gay CM, Byers LA, Zhang J. The current and emerging immunotherapy paradigm in small-cell lung cancer. Nat Cancer. 2025. Epub 20250605. doi: 10.1038/s43018-025-00992-5. PubMed PMID: 40473974.

72. Zugazagoitia J, Osma H, Baena J, Ucero AC, Paz-Ares L. Facts and hopes on cancer immunotherapy for small cell lung cancer. Clin Cancer Res. 2024. Epub 20240417. doi: 10.1158/1078-0432.CCR-23-1159. PubMed PMID: 38630789.

73. Zhang Y, Luo XY, Wu DH, Xu Y. ROR nuclear receptors: structures, related diseases, and drug discovery. Acta Pharmacol Sin. 2015;36(1):71–87. Epub 20141215. doi: 10.1038/aps.2014.120. PubMed PMID: 25500868; PMCID: PMC4571318.

74. Viragova S, Aparicio L, Palmerini P, Zhao J, Valencia Salazar LE, Schurer A, Dhuri A, Sahoo D, Moskaluk CA, Rabadan R, Dalerba P. Inverse agonists of retinoic acid receptor/retinoid X receptor signaling as lineage-specific antitumor agents against human adenoid cystic carcinoma. J Natl Cancer Inst. 2023;115(7):838–52. doi: 10.1093/jnci/djad062. PubMed PMID: 37040084; PMCID: PMC10323906.

75. He B, Nohara K, Park N, Park YS, Guillory B, Zhao Z, Garcia JM, Koike N, Lee CC, Takahashi JS, Yoo SH, Chen Z. The Small Molecule Nobiletin Targets the Molecular Oscillator to Enhance Circadian Rhythms and Protect against Metabolic Syndrome. Cell Metab. 2016;23(4):610–21. doi: 10.1016/j.cmet.2016.03.007. PubMed PMID: 27076076; PMCID: PMC4832569.

76. Huang Z, Zeng L, Ruan Z, Zeng Q, Yan H, Jiang W, Xiong Y, Zhou C, Yang H, Liu L, Dai J, Zou N, Xu S, Wang Y, Wang Z, Deng J, Chen X, Wang J, Xiang H, Li X, Duchemann B, Chen G, Xia Y, Mok T, Scheiermann C, Levi F, Yang N, Zhang Y. Time-of-day immunochemotherapy in non-small cell lung cancer: a randomized phase 3 trial. Nat Med. 2026. Epub 20260202. doi: 10.1038/s41591-025-04181-w. PubMed PMID: 41629425.

77. Gardner EE, Lok BH, Schneeberger VE, Desmeules P, Miles LA, Arnold PK, Ni A, Khodos I, de Stanchina E, Nguyen T, Sage J, Campbell JE, Ribich S, Rekhtman N, Dowlati A, Massion PP, Rudin CM, Poirier JT. Chemosensitive Relapse in Small Cell Lung Cancer Proceeds through an EZH2-SLFN11 Axis. Cancer Cell. 2017;31(2):286–99. doi: 10.1016/j.ccell.2017.01.006. PubMed PMID: 28196596; PMCID: PMC5313262.

78. Sen T, Rodriguez BL, Chen L, Della Corte C, Morikawa N, Fujimoto J, Cristea S, Nguyen T, Diao L, Li L, Fan Y, Yang Y, Wang J, Glisson BS, Wistuba, II, Sage J, Heymach JV, Gibbons DL, Byers LA. Targeting DNA damage response promotes anti-tumor immunity through STING-mediated T-cell activation in small cell lung cancer. Cancer Discov. 2019. doi: 10.1158/2159-8290.CD-18-1020. PubMed PMID: 30777870.

79. Kim D, Paggi JM, Park C, Bennett C, Salzberg SL. Graph-based genome alignment and genotyping with HISAT2 and HISAT-genotype. Nat Biotechnol. 2019;37(8):907–15. Epub 20190802. doi: 10.1038/s41587-019-0201-4. PubMed PMID: 31375807; PMCID: PMC7605509.

80. Liao Y, Smyth GK, Shi W. featureCounts: an efficient general purpose program for assigning sequence reads to genomic features. Bioinformatics. 2014;30(7):923–30. Epub 20131113. doi: 10.1093/bioinformatics/btt656. PubMed PMID: 24227677.

81. Love MI, Huber W, Anders S. Moderated estimation of fold change and dispersion for RNA-seq data with DESeq2. Genome Biol. 2014;15(12):550. Epub 2014/12/18. doi: 10.1186/s13059-014-0550-8. PubMed PMID: 25516281; PMCID: PMC4302049.

82. Stuart T, Srivastava A, Madad S, Lareau CA, Satija R. Single-cell chromatin state analysis with Signac. Nat Methods. 2021;18(11):1333–41. Epub 20211101. doi: 10.1038/s41592-021-01282-5. PubMed PMID: 34725479; PMCID: PMC9255697.

83. Hao Y, Stuart T, Kowalski MH, Choudhary S, Hoffman P, Hartman A, Srivastava A, Molla G, Madad S, Fernandez-Granda C, Satija R. Dictionary learning for integrative, multimodal and scalable single-cell analysis. Nat Biotechnol. 2024;42(2):293–304. Epub 20230525. doi: 10.1038/s41587-023-01767-y. PubMed PMID: 37231261; PMCID: PMC10928517.

84. McGinnis CS, Murrow LM, Gartner ZJ. DoubletFinder: Doublet Detection in Single-Cell RNA Sequencing Data Using Artificial Nearest Neighbors. Cell Syst. 2019;8(4):329–37 e4. Epub 20190403. doi: 10.1016/j.cels.2019.03.003. PubMed PMID: 30954475; PMCID: PMC6853612.

85. Zhang Y, Liu T, Meyer CA, Eeckhoute J, Johnson DS, Bernstein BE, Nusbaum C, Myers RM, Brown M, Li W, Liu XS. Model-based analysis of ChIP-Seq (MACS). Genome Biol. 2008;9(9):R137. Epub 2008/09/19. doi: 10.1186/gb-2008-9-9-r137. PubMed PMID: 18798982; PMCID: 2592715.

86. Korsunsky I, Millard N, Fan J, Slowikowski K, Zhang F, Wei K, Baglaenko Y, Brenner M, Loh PR, Raychaudhuri S. Fast, sensitive and accurate integration of single-cell data with Harmony. Nat Methods. 2019;16(12):1289–96. Epub 20191118. doi: 10.1038/s41592-019-0619-0. PubMed PMID: 31740819; PMCID: PMC6884693.

87. Hafemeister C, Satija R. Normalization and variance stabilization of single-cell RNA-seq data using regularized negative binomial regression. Genome Biol. 2019;20(1):296. Epub 20191223. doi: 10.1186/s13059-019-1874-1. PubMed PMID: 31870423; PMCID: PMC6927181.

88. Sun H, Zhou Y, Fei L, Chen H, Guo G. scMCA: A Tool to Define Mouse Cell Types Based on Single-Cell Digital Expression. Methods Mol Biol. 2019;1935:91–6. doi: 10.1007/978-1-4939-9057-3_6. PubMed PMID: 30758821.

89. Kolberg L, Raudvere U, Kuzmin I, Adler P, Vilo J, Peterson H. g:Profiler-interoperable web service for functional enrichment analysis and gene identifier mapping (2023 update). Nucleic Acids Res. 2023;51(W1):W207–W12. doi: 10.1093/nar/gkad347. PubMed PMID: 37144459; PMCID: PMC10320099.

90. Castanza AS, Recla JM, Eby D, Thorvaldsdottir H, Bult CJ, Mesirov JP. Extending support for mouse data in the Molecular Signatures Database (MSigDB). Nat Methods. 2023;20(11):1619–20. doi: 10.1038/s41592-023-02014-7. PubMed PMID: 37704782; PMCID: PMC11397807.

91. Hanzelmann S, Castelo R, Guinney J. GSVA: gene set variation analysis for microarray and RNA-seq data. BMC Bioinformatics. 2013;14:7. Epub 20130116. doi: 10.1186/1471-2105-14-7. PubMed PMID: 23323831; PMCID: PMC3618321.

92. Zhang L, Zhang J, Nie Q. DIRECT-NET: An efficient method to discover cis-regulatory elements and construct regulatory networks from single-cell multiomics data. Sci Adv. 2022;8(22):eabl7393. Epub 20220601. doi: 10.1126/sciadv.abl7393. PubMed PMID: 35648859; PMCID: PMC9159696.

93. Greenwald NF, Miller G, Moen E, Kong A, Kagel A, Dougherty T, Fullaway CC, McIntosh BJ, Leow KX, Schwartz MS, Pavelchek C, Cui S, Camplisson I, Bar-Tal O, Singh J, Fong M, Chaudhry G, Abraham Z, Moseley J, Warshawsky S, Soon E, Greenbaum S, Risom T, Hollmann T, Bendall SC, Keren L, Graf W, Angelo M, Van Valen D. Whole-cell segmentation of tissue images with human-level performance using large-scale data annotation and deep learning. Nat Biotechnol. 2022;40(4):555–65. Epub 20211118. doi: 10.1038/s41587-021-01094-0. PubMed PMID: 34795433; PMCID: PMC9010346.

94. Rumberger JL, Greenwald NF, Ranek JS, Boonrat P, Walker C, Franzen J, Varra SR, Kong A, Sowers C, Liu CC, Averbukh I, Piyadasa H, Vanguri R, Nederlof I, Wang XJ, Van Valen D, Kok M, Bendall SC, Hollmann TJ, Kainmueller D, Angelo M. Automated classification of cellular expression in multiplexed imaging data with Nimbus. Nat Methods. 2025;22(10):2161–70. Epub 20251008. doi: 10.1038/s41592-025-02826-9. PubMed PMID: 41062826.

95. Van Gassen S, Callebaut B, Van Helden MJ, Lambrecht BN, Demeester P, Dhaene T, Saeys Y. FlowSOM: Using self-organizing maps for visualization and interpretation of cytometry data. Cytometry A. 2015;87(7):636–45. Epub 20150108. doi: 10.1002/cyto.a.22625. PubMed PMID: 25573116.

